# BrainScale, Enabling Scalable Online Learning in Spiking Neural Networks

**DOI:** 10.1101/2024.09.24.614728

**Authors:** Chaoming Wang, Xingsi Dong, Zilong Ji, Jiedong Jiang, Xiao Liu, Si Wu

**Affiliations:** Guangdong Institute of Intelligence Science and Technology, Guangdong, China; School of Psychological and Cognitive Sciences, Peking University, Beijing, China; Center of Quantitative Biology, Peking-Tsinghua Center for Life Sciences, Academy for Advanced Interdisciplinary Studies, Peking University, Beijing, China; PKU-IDG/McGovern Institute for Brain Research, Peking University, Beijing, China; Institute of Cognitive Neuroscience, University College London, London, United Kingdom; Beijing International Center for Mathematical Research, Peking University, Beijing, China; Janelia Research Campus, Howard Hughes Medical Institute, Ashburn, VA, USA

## Abstract

Spiking neural networks (SNNs) represent a promising paradigm for understanding brain functions [1, 2] and developing neuromorphic intelligence [3, 4]. However, their potential remains largely unrealized due to a fundamental limitation: the lack of a scalable online learning system capable of supporting large-scale training of complex brain dynamics over long timescales. Current approaches are either limited to offline learning [5], impeding long-time task training; constrained by high memory complexity [6, 7], hindering network scaling; or using oversimplified models [8], failing to capture complex brain dynamics. Here, we introduce Brain-Scale, a system that synergizes model universality, computational efficiency, and engineering usability to enable scalable online learning in SNNs. First, BrainScale introduces standard model abstractions to support the training of diverse spiking networks. Second, BrainScale implements an online learning algorithm with linear memory complexity by exploiting intrinsic properties of SNN dynamics. Third, BrainScale provides an online learning compiler that automates the implementation of online learning for any user-defined models. Extensive evaluations across diverse SNN dynamics and computational tasks demonstrate that BrainScale consistently delivers strong training performance while maintaining extremely low memory usage and high computational efficiency. These properties enable the online training of a whole-brain-scale SNN of the *Drosophila* brain, accurately capturing functional activities across brain regions. These results position BrainScale as a foundational platform for advancing large-scale brain simulation and neuromorphic computing.

## Introduction

Spiking neural networks (SNNs) are a type of neural network that captures the spike-based, event-driven communication of biological neurons. By reproducing the precise timing of action potentials, SNNs provide a biologically interpretable framework for probing cognitive functions such as sensory processing, memory formation, and decision-making [1, 2, 9]. Their event-driven architecture also delivers substantial gains in energy efficiency and spatiotemporal information processing compared with conventional artificial neural networks (ANNs) [10–12]. Together, these strengths position SNNs as a critical bridge between fundamental neuroscience and practical applications in neuromorphic engineering [1–3, 12].

Despite their theoretical advantages, SNNs encounter fundamental scalability challenges. Specifically, when scaling SNN complexity to match the neuron number and neuronal dynamics found in biological brains, problems like network optimization become intractable. The dominant training approach, backpropagation through time (BPTT) algorithm with surrogate gradients [5, 13, 14], is an offline training method and incurs memory costs that increase linearly with the number of time steps *T*. This creates prohibitive memory demands when modeling biological timescales, where a single second may contain thousands of discrete time steps, thus limiting its applications on long time sequences and large-scale SNNs [11]. Recent online learning algorithms for SNNs attempt to address memory constraints by training models in a forward-time manner, limiting memory consumption to a single time step rather than storing entire temporal sequences [15]. While promising, these algorithms typically face one or more of the following three fundamental limitations that must be addressed for the scalability issue.

1. **Computational inefficiency**. State-of-the-art online learning methods that approach BPTT performance, such as e-prop [7] and OSTL [6], suffer from 𝒪(*N* ^2^) quadratic memory complexity (where *N* is the number of neurons), making them impractical for large-scale simulations.
2. **Limited model generalization**. While some algorithms achieve 𝒪(*N*) memory complexity [8, 16, 17], they are specifically designed for very simple neuron models, e.g., leaky integrate-and-fire (LIF) neurons, and therefore lack the generalizability to more complex biological neuronal dynamics.
3. **Poor engineering implementation**. The practical success of a learning algorithm relies heavily on a robust programming framework that supports its automatic implementation. For instance, the widespread adoption of automatic differentiation (AD) frameworks for ANNs like PyTorch [18] and TensorFlow [19] has significantly reduced the effort required to integrate models with backpropagation. In contrast, current online learning algorithms lack such automatic and user-friendly systems, limiting their broader applicability and real-world impact.

In this study, we introduce the BrainScale system, designed to support the training of large-scale SNNs by balancing model universality, computational efficiency, and engineering usability. In terms of model universality, BrainScale simplifies general SNNs into two fundamental models: `AlignPre` and `AlignPost`. In terms of computational efficiency, BrainScale leverages the sparse, event-driven nature of spiking neurons to reformulate real-time recurrent learning (RTRL) [20], which achieves linear memory scaling relative to the number of neurons while maintaining gradient approximation accuracy. In terms of engineering usability, BrainScale provides an online learning compiler that automatically generates efficient online learning code for any complex SNNs, facilitating implementations across various hardware platforms, including CPUs, GPUs, and TPUs.

To validate BrainScale’s effectiveness, we perform extensive experiments across diverse SNN models, neuromorphic datasets, and computational tasks. Our results demonstrate that BrainScale achieves extremely low memory consumption and computational efficiency, compared to offline BPTT-based and online algorithms with quadratic memory complexity. For standard neuromorphic tasks, BrainScale achieves training performance comparable to BPTT. In brain simulation models, it enables the training of SNNs in simulating brain dynamics involved in typical cognitive tasks. Remarkably, BrainScale facilitates the training of a whole-brain-scale SNN model of the *Drosophila* brain [21, 22], successfully fitting the functional activities across the entire *Drosophila* brain [23, 24]. By addressing the critical challenges of scalable online learning for SNNs, BrainScale represents an important advancement in understanding brain dynamics at scale and facilitating the development of large-scale neuromorphic intelligent systems.

## Results

### BrainScale supports a wide range of spiking networks

BrainScale supports a wide range of SNNs with the following discretized dynamics (SI. A):

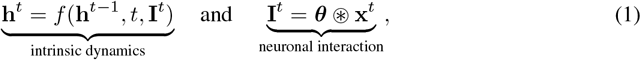

where **h**^*t*^ ∈ℝ^*Hd*^ represents the hidden states of *H* neurons, with each neuron quantified by *d* state variables; **I**^*t*^ ∈ℝ^*H*^ represents the synaptic currents; and *t* represents the time stamp; **x**^*t*^ ∈ ℝ^*I*^ represents synaptic inputs (e.g., presynaptic spikes); and ***θ*** ∈ℝ^*P*^ denotes connection weights between pre- and postsynaptic neurons (Fig. 1).

**Figure 1.**
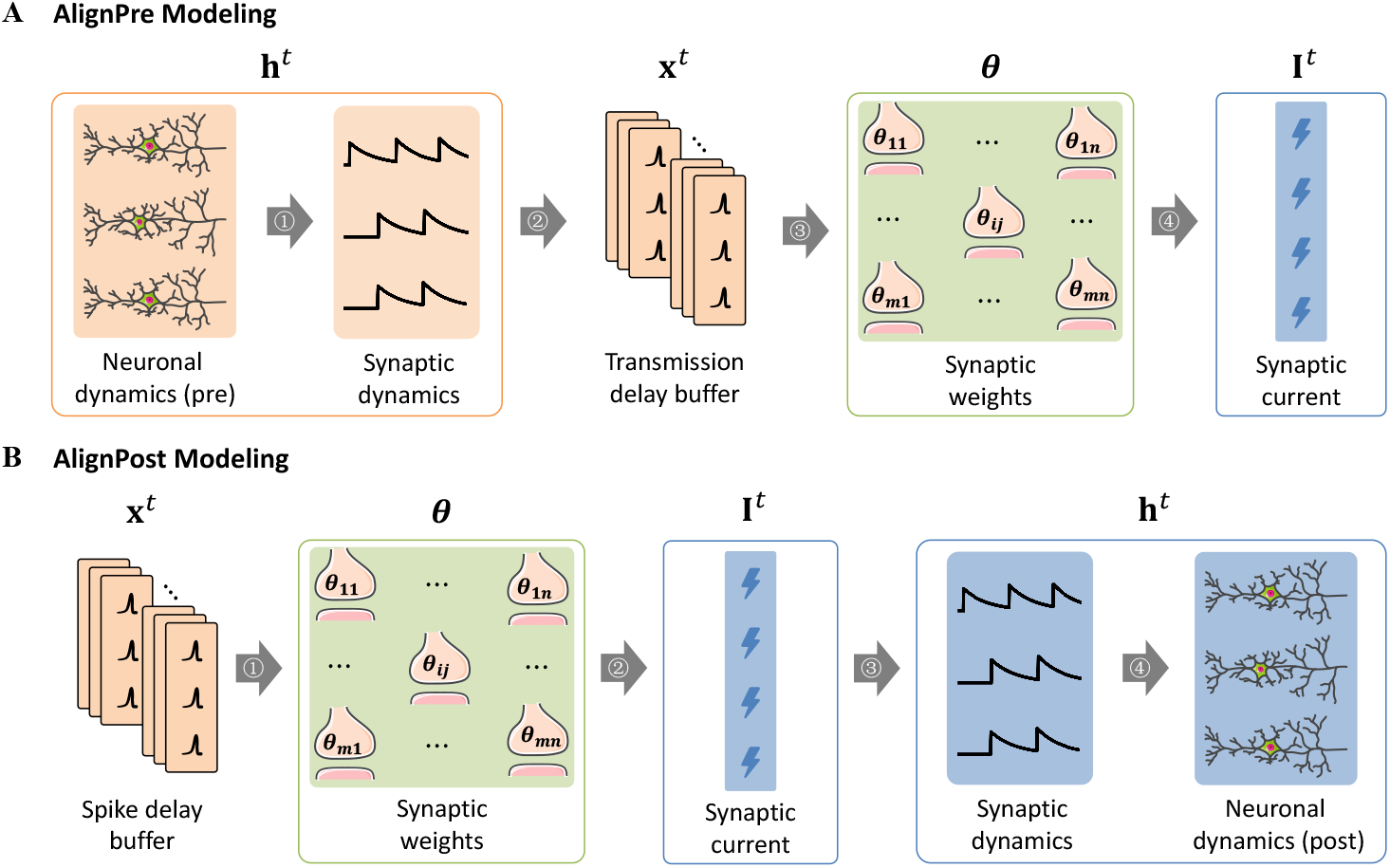
AlignPre **and** AlignPost abstractions for general spiking networks. **A**. `AlignPre` modeling, where synaptic variables are aligned with the presynaptic neuronal dimensions. Presynaptic spikes drive the updates of synapses ➀. The resulting delayed synaptic dynamics ➁ are then transmitted to postsynaptic sites ➂, generating postsynaptic currents ➃, which in turn influence the evolution of postsynaptic neuron dynamics. **B**. `AlignPost` modeling, where synaptic variables are aligned with the postsynaptic neuronal dimensions. The delayed presynaptic spikes ➀ determine the postsynaptic inputs ➁. These inputs drive the updates of synaptic dynamics ➂, which then influence the evolution of postsynaptic neuron dynamics ➃. In `AlignPre`, the input **x** governing neuronal interactions in Eq. 1 represents conductance of synaptic dynamics; while in `AlignPost`, the variable **x** in Eq. 1 corresponds to presynaptic spikes. The orange color represents the presynaptic dynamics with dimension ℝ^*m*^, the blue color denotes the postsynaptic dynamics with dimension ℝ^*n*^, and the green color represents the synaptic interaction ***θ*** with dimension ℝ^*m×n*^.

SNNs described by Eq. 1 are implemented using the `AlignPre` and `AlignPost` frameworks [25, 26] (Fig. 1; see examples in SI. B). Specifically, in `AlignPre`, synapses with identical parameters that share the same presynaptic neuron exhibit identical dynamical trajectories (Fig. S6), enabling synaptic variables to be aligned with the presynaptic neuronal dimension (Fig. 1A). In `AlignPost`, when synapses feature exponential dynamics with identical time constants, multiple synapses from presynaptic neurons converging onto a single postsynaptic neuron can be collectively represented by a single shared exponential synapse. This shared representation captures the combined input to the postsynaptic neuron (Fig. S6), allowing synaptic variables to be aligned along the postsynaptic dimension (Fig. 1B).

`AlignPre` and `AlignPost` decompose an SNN into two parts (Eq. 1): (1) an *intrinsic dynamics* part that dictates the temporal evolution of neuronal states through the element-wise activation function *f*, and (2) a *neuronal interaction* part that converts presynaptic inputs into postsynaptic currents via the interaction operator ⊛. The decomposition can largely reduce computational complexity and memory consumption, enabling a simplified algorithm for SNN online learning (described in the next section).

### BrainScale achieves linear-memory online learning algorithm

The objective of SNN training is to find a group of synaptic weights ***θ*** that minimize a loss function ℒ that quantifies the discrepancy between target values and network outputs. When target values are provided at discrete time points *t* ∈ 𝒯, we compute the weight gradient by:

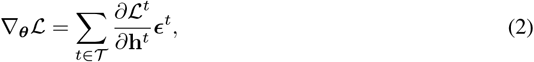

where ℒ^*t*^ represent the instantaneous loss at time *t, ∂* ℒ^*t*^*/∂***h**^*t*^ denotes the learning signal that arrival at the hidden state **h**^*t*^, and ***ϵ***^*t*^ is the eligibility trace indicating how parameters influence the hidden state, which is computed as [20]:

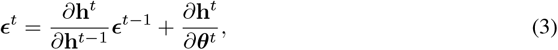

where the hidden Jacobian *∂***h**^*t*^*/∂***h**^*t−*1^ = **D**^*t*^ + **J**^*t*^ consists of a diagonal matrix **D**^*t*^ derived from intrinsic dynamics *f*, and a square matrix **J**^*t*^ derived from neuronal interactions ⊛ (see SI. C). The hidden-to-weight Jacobian *∂***h**^*t*^*/∂****θ***^*t*^ captures the effect of weight changes on hidden states. Importantly, Eq. 3 imposes a memory overhead of 𝒪 (*H*^3^) (Fig. 2A).

**Figure 2.**
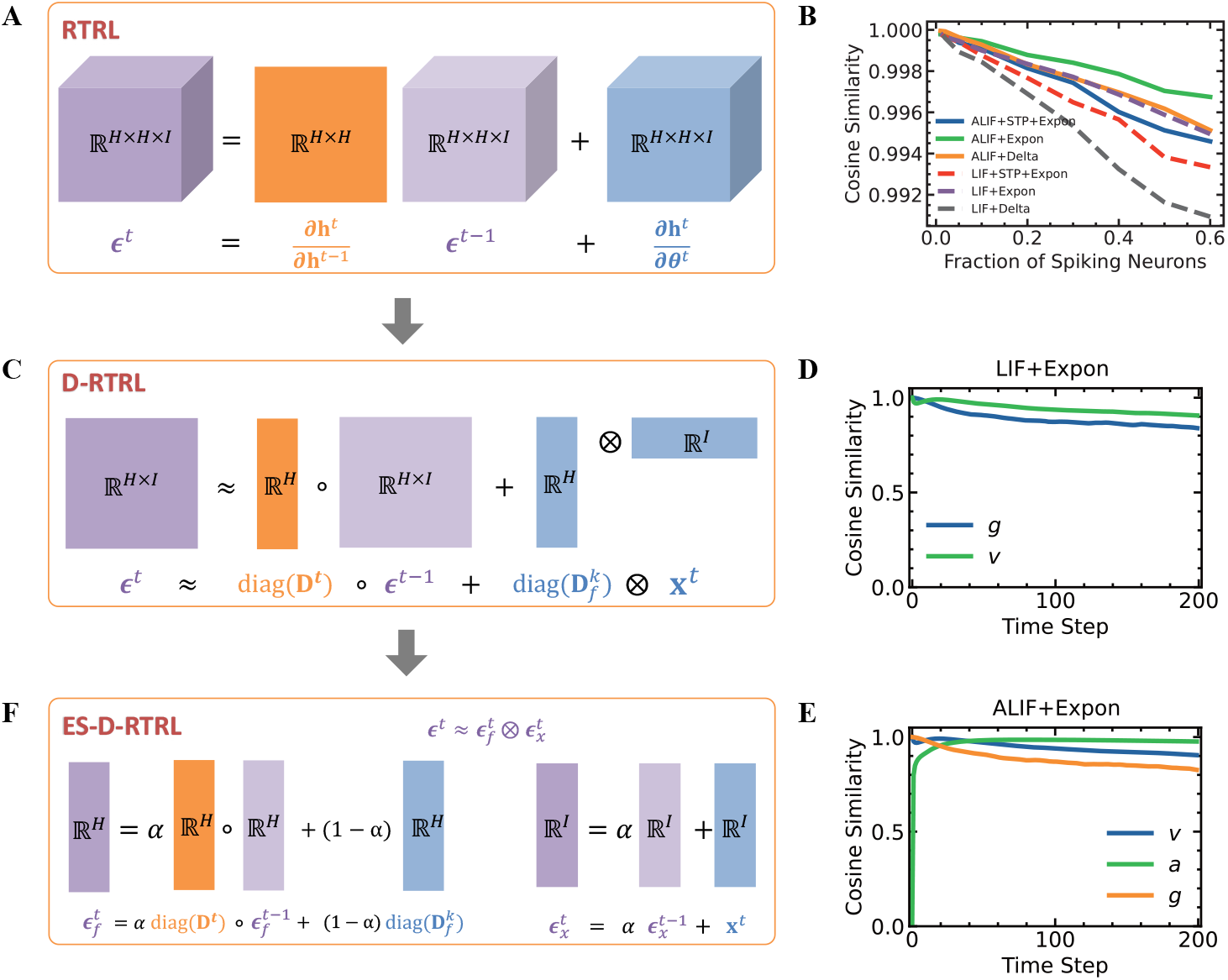
Reducing the complexity of online learning algorithms. **A**. The visualization of RTRL computation complexity reveals that ***ϵ*** and the parameter Jacobian *∂***h**^*t*^*/∂****θ***^*t*^ exhibit cubic memory complexity, while the hidden Jacobian *∂***h**^*t*^*/∂***h**^*t−*1^ demonstrates quadratic memory complexity. **B**. Cosine similarity between the diagonal Jacobian **D**^*t*^ induced by intrinsic dynamics in Eq. 1 and the full hidden Jacobian *∂***h**^*t*^*/∂***h**^*t−*1^, over different network dynamics and fraction of spiking neurons. **C**. In D-RTRL computation, the eligibility trace ***ϵ*** is reduced to exhibit quadratic memory complexity. **D**. Cosine similarity between the rank-one approximation (Eq. 5) and D-RTRL eligibility trace (Eq. 4) for LIF neurons with exponential synapse (Eq. S46). Similarity is computed for membrane potential (*v*) and synaptic conductance (*g*). **E**. Cosine similarity between the rank-one approximation (Eq. 5) and D-RTRL eligibility trace (Eq. 4) for ALIF neurons with exponential synapse (Eq. S47). Similarity is computed for membrane potential (*v*), synapse (*g*), and adaptation (*a*) variables. **F**. In ES-D-RTRL computation, the eligibility trace ***ϵ*** is approximated using two variables with linear memory complexity that separately track pre-synaptic and post-synaptic neuronal activities.

By leveraging SNN’s inherent properties (SI. C), we can reduce the complexity to 𝒪 (*H*). Without loss of generality, we consider each neuron has only 1 state variable with *d* = 1 (but see SI. G for generalized results for *d >* 1). First, we found that a diagonal approximation of the hidden Jacobian captures over 99% of the cosine similarity with the full Jacobian (*∂***h**^*t*^*/∂***h**^*t−*1^ ≈ **D**^*t*^, Fig. 2B). Second, we proved (SI. D) that the hidden-to-weight Jacobian can be decomposed as 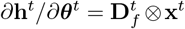, where 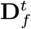 is a diagonal matrix with entries 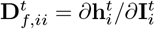, and ⊗ represents the Kronecker product. These observations enable us to simplify the eligibility trace in Eq. 3 to:

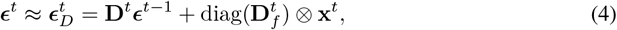

which we referred to as the diagonal-approximated RTRL (D-RTRL) (Fig. 2C). This diagonal approximation reduced the memory complexity from 𝒪 (*H*^3^) to 𝒪 (*H*^2^).

We further observed that 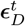 can be expressed as a summation of *t* rank-one matrices: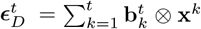, where 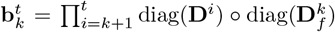 (Fig. S7). Since inputs **x**^*t*^ maintain consistent signs across time steps—whether as binary spikes in `AlignPost` or synaptic conductance in `AlignPre` (Fig. 1), we proved that the sum of multiple rank-one matrices can be approximated as a single rank-one matrix between summed vectors (SI. E). Consequently, the eligibility trace can be simplified to:

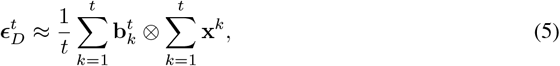

which still preserves a high cosine similarity with the full eligibility trace across LIF (Fig. 2D), ALIF (Fig. 2E), and other SNN architectures (Fig. S1).

In summary, this approximation gives the exponentially smoothed diagonal approximated RTRL (ES-D-RTRL) algorithm (Fig. 2F):

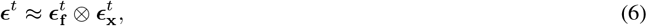

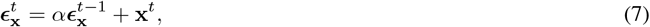

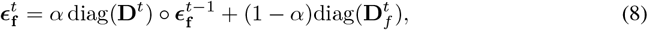

Where ∘ denotes element-wise multiplication, and *α* (0 *< α <* 1) is the smoothing factor corresponding to time constant τ = −Δ*t/* ln(*α*). ***ϵ***_**x**_∈ ℝ^*I*^ and ***ϵ***_**f**_∈ ℝ^*H*^ represent eligibility traces tracking pre-and postsynaptic activity, respectively. The eligibility trace in ES-D-RTRL operates temporally, locally, and in an unsupervised way, effectively achieving memory complexity of 𝒪 (*H*).

### BrainScale facilitates automated online learning compilation

The widespread success of ANNs primarily stems from the automation achieved in frameworks such as PyTorch [18] and TensorFlow [19] for their training algorithms, namely backpropagation and BPTT algorithms. To provide similar training convenience for SNNs, we introduced a compiler (Fig. 3A-E) that automates the translation of SNN models into efficient online learning implementations of D-RTRL and ES-D-RTRL, and subsequently deploys both models and generated algorithms across various hardware backends, including CPUs, GPUs, and TPUs (Fig. 3).

**Figure 3.**
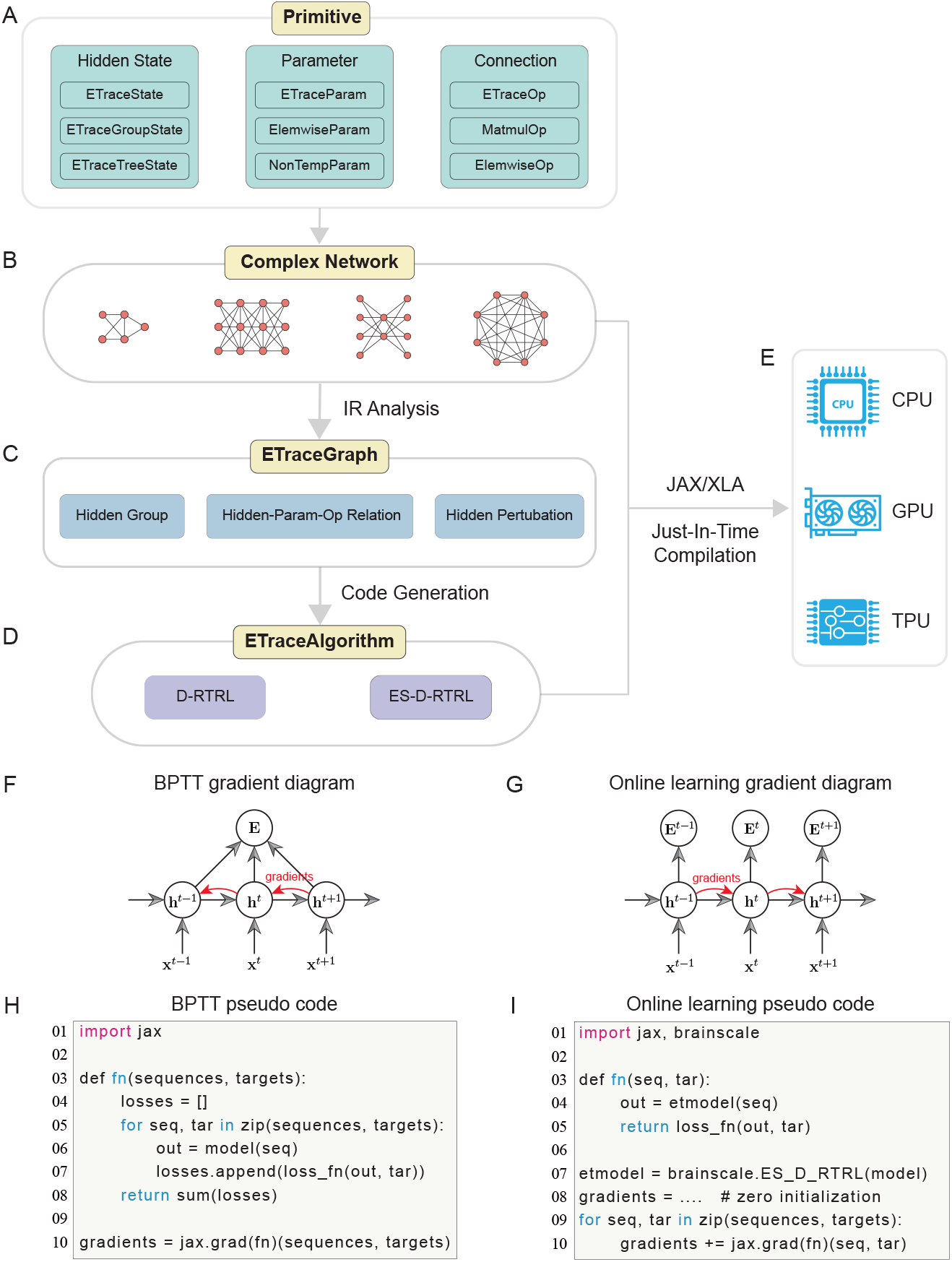
The architecture of online learning compilation. **A**. Three types of primitives are defined to structure spiking network modeling: `Hidden State` for neuronal and synaptic states, `Parameter` for trainable parameters, and `Connection` for synaptic connectivity types. **B**. Three primitives enable the construction of spiking networks with any complex network dynamics and topologies. **C**. BrainScale uses IR analysis to abstract Python-defined models into symbolic representations, then analyzes the graphs between hidden groups, weight parameters, and synaptic connections. **D**. Based on the analyzed graphs, the compiler generated code implementing the user-selected algorithm—either D-RTRL or ES-D-RTRL. **E**. Using the JIT compilation capability of JAX and XLA, both user-coded models and compiler-generated algorithms are compiled and executed on CPUs, GPUs, and TPUs. **F**. Gradient computation diagram for BPTT autograd implementation, showing backward gradient accumulation. **G**. Gradient computation diagram for online D-RTRL and ES-D-RTRL implementation, showing forward gradient accumulation. **H**. Pseudo code for the implementation of BPTT gradient diagram, where the `jax.grad` function processes the complete sequence at once. **I**. Pseudo code for the implementation of online learning gradient diagram, where `jax.grad` is applied at each step, and gradient computation is performed in forward mode as the sequence unrolls temporally.

At the `Primitive` layer, the system encapsulates three primitives that capture the essential components of general SNNs (as formalized in Eq. 1, Fig. 3A). Specifically, `Hidden State` primitives represent internal neuronal and synaptic variables, such as membrane potentials and synaptic conductances; `Parameter` primitives denote trainable parameters, such as synaptic weights; while `Connection` primitives describe network connectivities, including dense, convolutional, sparse connections, and element-wise operations. By flexibly composing these primitives, the `Complex Network` layer supports the construction of complex SNNs with sophisticated dynamics and topologies (Fig. 3B).

The core innovation of the compiler lies in `ETraceGraph` and `ETraceAlgorithm` layers. At the `ETraceGraph` stage, user-defined Python-based models are transformed into symbolic intermediate representations (IRs), which explicitly encode data dependencies between model parameters and neuronal states. The compiler analyzes these IRs to extract higher-level computational graphs that describe how each model parameter, ***θ***, influences the hidden state through specific connections ⊛. The resulting graph captures all critical information necessary for computing eligibility traces, including the hidden-to-hidden Jacobian (diag(**D**^*t*^)) and hidden-to-weight Jacobians (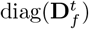 and **x**^*t*^) (Fig. 3C).

Subsequently, at the `ETraceAlgorithm` layer, the compiler automatically generates optimized online learning implementations based on the user-selected algorithm (Fig. 3D). The generated code includes both the logic for updating eligibility traces during state evolution (see Eq. 4 for D-RTRL and Eqs. 6-8 for ES-D-RTRL) and the gradient computation triggered by learning signals (see Eq. 2). During this code generation process, the compiler also performs advanced optimizations, such as reusing shared eligibility traces within the same neuron population to minimize memory footprint, and vectorizing hidden-group Jacobian computations to maximize computational efficiency (SI. L).

Finally, leveraging just-in-time (JIT) compilation via JAX and XLA [27], both user-defined models (Fig. 3B) and compiler-generated algorithms (Fig. 3D) are automatically deployed onto the target hardware platform (Fig. 3E). The hardware-agnostic nature of the intermediate representations ensures that the compiled online learning code inherently supports cross-platform deployment onto CPUs, GPUs, and TPUs.

Through this hierarchical online learning compilation, BrainScale provides a unified and highly automated solution for computing forward gradients (see comparison of Fig. 3F for BPTT and Fig. 3G for online learning) without introducing additional programming difficulties (see comparison of Fig. 3H for BPTT and Fig. 3I for online learning; SI. P). This end-to-end online learning system will greatly lower the barrier to entry for using online learning in SNNs, making advanced algorithms more accessible and deployable in real-world applications.

### Algorithmic evaluations on approximation, long-term dependency, memory, and speed

ES-D-RTRL incorporates different approximations for computing parameter gradients. To assess its effectiveness in approximating the hidden Jacobian and weight gradients on real-world neuromorphic datasets, we conducted experiments using IBM DVS Gesture [28], Spiking Heidelberg Digits (SHD) [29], and Neuromorphic-MNIST (N-MNIST) [30] datasets. We explored various SNN dynamics, ranging from LIF to ALIF neurons, and synapses including Delta, exponential, short-term depression (STD), and short-term plasticity (STP) (SI. B). We first evaluated the cosine similarity between the single-step hidden Jacobian (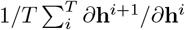 Fig. 4A) and the long-range hidden Jacobian (*∂***h**^*T*^ */∂***h**^1^; Fig. 4B), where *T* is the dataset length. We observed consistently high cosine similarity between the two. Notably, high similarity values were especially prominent in networks with complex intrinsic dynamics, such as ALIF-based architectures (Fig. 4B). We then assessed parameter gradient approximations by comparing the cosine similarity between gradients computed by our online learning algorithms and BPTT. The gradients computed by D-RTRL exhibited a cosine similarity (Fig. 4C) comparable to its hidden Jacobian approximation (Fig. 4A). ES-D-RTRL, however, demonstrated a relatively lower gradient similarity compared to D-RTRL (Fig. 4D), due to the additional approximations involved in computing eligibility traces (SI. G). Moreover, both algorithms exhibited lower gradient similarities on N-MNIST (Fig. 4C-D), indicating some dataset dependence. Despite this, both D-RTRL and ES-D-RTRL gradients maintained high approximation, demonstrating their effectiveness in capturing essential gradients for training SNNs on neuromorphic datasets. Further investigation revealed several key insights about factors affecting the approximation performance (SI. H). Network size showed no significant impact on approximation accuracy (Fig. S9). However, approximation accuracy declined greatly when either feedforward (Fig. S10) or recurrent (Fig. S11) weight strength increased to levels that produced unrealistically high firing rates. Additionally, deeper recurrent layers led to lower approximation accuracies (Fig. S12), suggesting a preference for shallow network architectures in our algorithms.

**Figure 4.**
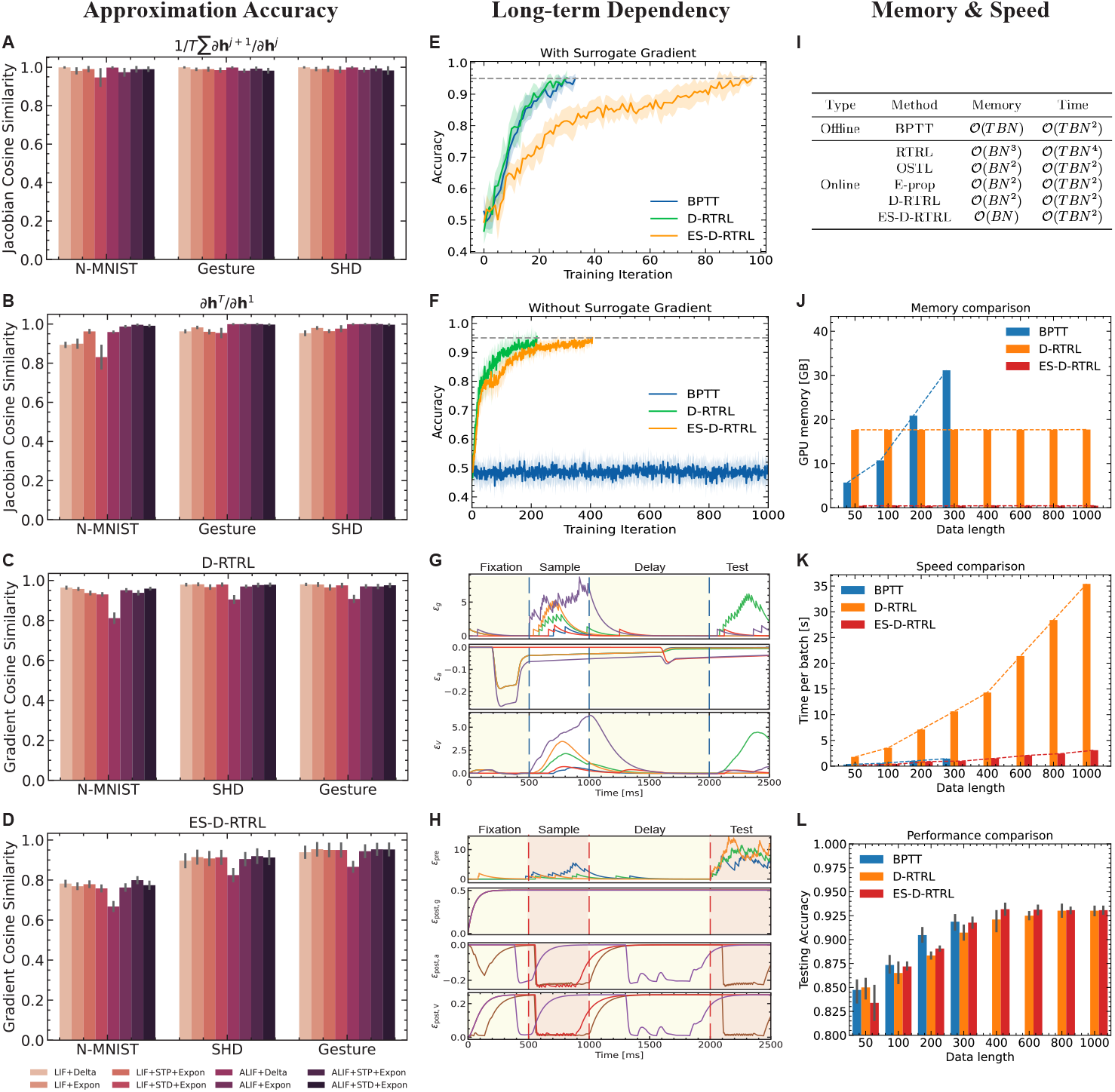
Evaluation of learning algorithms on approximation accuracy, long-term dependency learning, memory usage, and computational speed. **A-D**. Approximation accuracy evaluations across three neuromorphic datasets and eight network dynamics. For each dataset, we randomly selected 500 data points, computed the Jacobian cosine similarity for each data point, and reported the average cosine similarity across all samples. **A**. Cosine similarity between approximated and exact single-step hidden Jacobians: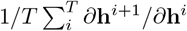. **B**. Cosine similarity between approximated and exact long-range hidden Jacobians: *∂***h**^*T*^ */∂***h**^1^. **C**. Cosine similarity between weight gradients computed by D-RTRL and BPTT algorithms. **D**. Cosine similarity between weight gradients computed by ES-D-RTRL and BPTT algorithms. **E-H**. Evaluation of long-term dependency learning capability using the DMTS task. **E**. Comparison of testing accuracy for different algorithms on networks utilizing surrogate gradients. **F**. Comparison of testing accuracy for different algorithms on networks without surrogate gradients. **G**. Visualization of the eligibility trace in the D-RTRL algorithm, showing data from five randomly sampled positions. **H**. Visualization of the eligibility trace in the ES-D-RTRL algorithm, displaying data from three randomly sampled positions. **I-L**. Comparison of memory efficiency and running speed between different learning algorithms. All evaluations were performed using the Gesture dataset [28]. **I**. Theoretical analysis of memory and computational complexity. **J**. Comparison of GPU memory consumption between our online learning algorithms and BPTT during one batch of training. **K**. Comparison of training speed for one batch between our online learning algorithms and BPTT during one batch training. **L**. Comparison of final testing accuracy over different data lengths.

Spiking networks hold promise for modeling long-term dependencies in datasets. Therefore, we evaluated whether ES-D-RTRL is sufficient for effectively learning very long sequences. We trained recurrent spiking networks (Eq. S47) on the delayed match-to-sample (DMTS) task [31], in which the network is required to maintain directional information from the sample phase over a long delay period (>1000 ms) before making a decision based on the test period stimulus (SI. I). We initially applied surrogate spiking gradients [5] on the network training. Experiments demonstrated that D-RTRL, ES-D-RTRL, and BPTT all effectively solve the task, with BPTT and D-RTRL showing faster convergence compared to ES-D-RTRL (Fig. 4E). However, without surrogate gradients, BPTT failed to train successfully on this task, with its training accuracy fluctuating around the 50% random baseline (Fig. 4F). In contrast, D-RTRL and ES-D-RTRL were capable of learning normally without compromising training accuracy. These findings highlight that utilizing intrinsic dynamics to train spiking networks can effectively capture long-term dependency information from spiking data. Then, we investigated how our algorithm achieves long-term dependency learning. We found that D-RTRL maintains three eligibility traces: ***ϵ***_**g**_, ***ϵ***_**a**_, and ***ϵ***_**v**_, corresponding to the hidden states **g, a**, and **v** (Eq. S47). Specifically, ***ϵ***_**g**_ tracks spiking inputs, increasing upon arrival of spiking stimuli and gradually decaying during the delay period; *ϵ*_**a**_ tracks the influence Jacobian with slow dynamics, facilitating information transfer across the delay; ***ϵ***_**v**_ integrates information tracked from ***ϵ***_**g**_ and ***ϵ***_**a**_, exhibiting sensitivity to input changes as ***ϵ***_**g**_ while gradually decaying as ***ϵ***_**a**_ (Fig. 4G). In contrast, ES-D-RTRL exhibits markedly different learning behavior (Fig. 4H). The presynaptic eligibility trace ***ϵ***_pre_ behaves similarly to ***ϵ***_**g**_ in D-RTRL but is driven solely by presynaptic spikes. While ***ϵ***_post,**g**_ tracks postsynaptic current information and maintains a constant value throughout. Similar to D-RTRL, ***ϵ***_post,**a**_ and ***ϵ***_post,**v**_ exhibited slow dynamics, delivering long-term postsynaptic Jacobian information over a long time, until the arrival of learning signals during the test period. Finally, we analyzed the differences in synaptic connectivity learned by ES-D-RTRL compared to BPTT and D-RTRL. Our analysis revealed that ES-D-RTRL maintained similar weight distributions with D-RTRL and BPTT, yet it displayed distinct column structures, particularly in recurrent weights (Fig. S15). This pattern, where incoming synaptic weights for a subset of postsynaptic neurons tend to be consistently stronger or weaker together, largely stems from the learning mechanism of ES-D-RTRL (see Eqs. 7-8). Interestingly, similar column structures have been observed in the synaptic connectome of layer 4 in the somatosensory cortex (Fig. S16, adapted from Fig. 3E in [32]), suggesting that ES-D-RTRL may capture biologically relevant connectivity patterns.

Finally, we evaluated the memory and computational complexity of ES-D-RTRL in comparison to other algorithms. Fig. 4I summarize the theoretical analysis. In general, ES-D-RTRL achieves a memory complexity of 𝒪 (*BN*)—scaling linearly with the number of neurons—since it only requires storing two activity traces: one for the presynaptic neurons and one for the postsynaptic neurons. In contrast, other online learning algorithms exhibit at least 𝒪 (*BN* ^2^) memory complexity.

Here, *B* denotes the batch size. For computational complexity, ES-D-RTRL requires 𝒪 (*TBN* ^2^) operations for full gradient computation, but only 𝒪 (*TBN*) operations for computing eligibility traces. This is in contrast to OSTL, E-prop, and D-RTRL, which incur quadratic complexity in computing eligibility traces due to their reliance on more expensive matrix-based updates. We then conducted empirical experiments on the DVS Gesture dataset [28] to compare the memory usage and computational speed of BPTT, D-RTRL, and ES-D-RTRL across varying time steps. The evaluated network consisted of three recurrent layers, featuring LIF dynamics with Delta synapses (Eq. S44). Our experiments showed that BPTT’s GPU memory usage increased linearly with the number of time steps, reaching its limit at 400 time steps due to an out-of-memory error (Fig. 4J). In contrast, D-RTRL and ES-D-RTRL required only constant memory regardless of the data time length. Particularly, ES-D-RTRL consumed less than 0.5 GB of GPU runtime memory throughout training, while D-RTRL required approximately 17.5 GB. Computational time evaluations (Fig. 4K) revealed that all three algorithms’ time consumption scaled linearly with the number of time steps. However, ES-D-RTRL ran significantly faster than BPTT and achieved ten times the acceleration compared to D-RTRL. The observed memory and speed advantages hold consistently across CPU and TPU platforms (Fig. S13 and S14). We also recorded training performance during our evaluations. Fig. 4L presents the mean testing accuracy of ten evaluations, with a 95% confidence interval. All three algorithms performed better as the data sequence length increased. BPTT outperformed D-RTRL and ES-D-RTRL when the data length was 200, while at other lengths, D-RTRL and ES-D-RTRL showed comparable performance.

### Performance evaluations on neuromorphic tasks using task-oriented spiking networks

Spiking networks have been widely used in neuromorphic tasks [3, 11]. To evaluate the effectiveness of our ES-D-RTRL algorithm for neuromorphic applications, we conducted extensive tests using diverse SNN models on three standard neuromorphic benchmarks, including N-MNIST [30], SHD [29], and IBM DVS Gesture [28]. The types of SNN evaluated here include: (1) LIF network [38], LIF neurons connected with delta synapses (Eq. S44) without recurrent connections; (2) RLIF network [38], LIF neurons connected with delta synapses (Eq. S44) with recurrent connections; (3) adLIF network [38], ALIF neurons connected with delta synapses (Eq. S45) without recurrent connections; (4) RadLIF network [38], ALIF neurons connected with delta synapses (Eq. S45) with recurrent connections; (5) and event-based gated recurrent unit (EGRU) [39] (Eq. S56).

The aim of our evaluation is to achieve a fair comparison of the learning performance of different algorithms under identical hyperparameter settings. Table 1 presents the mean and standard deviation of classification accuracy over at least three independent runs for each dataset. The results indicate that ES-D-RTRL matches BPTT’s performance across most tested datasets and network architectures. Conversely, D-RTRL demonstrates poorer performance, which is particularly clear from its reduced training accuracy on SHD. Interestingly, this reduced accuracy occurs despite the gradient computed by D-RTRL having a higher cosine similarity to the BPTT-calculated gradient than ES-D-RTRL’s gradient does (as shown in Fig. 4C-D). This finding suggests a more complex relationship between gradient similarity and task performance than previously assumed, under the context of spiking networks and neuromorphic datasets.

**Table 1.**
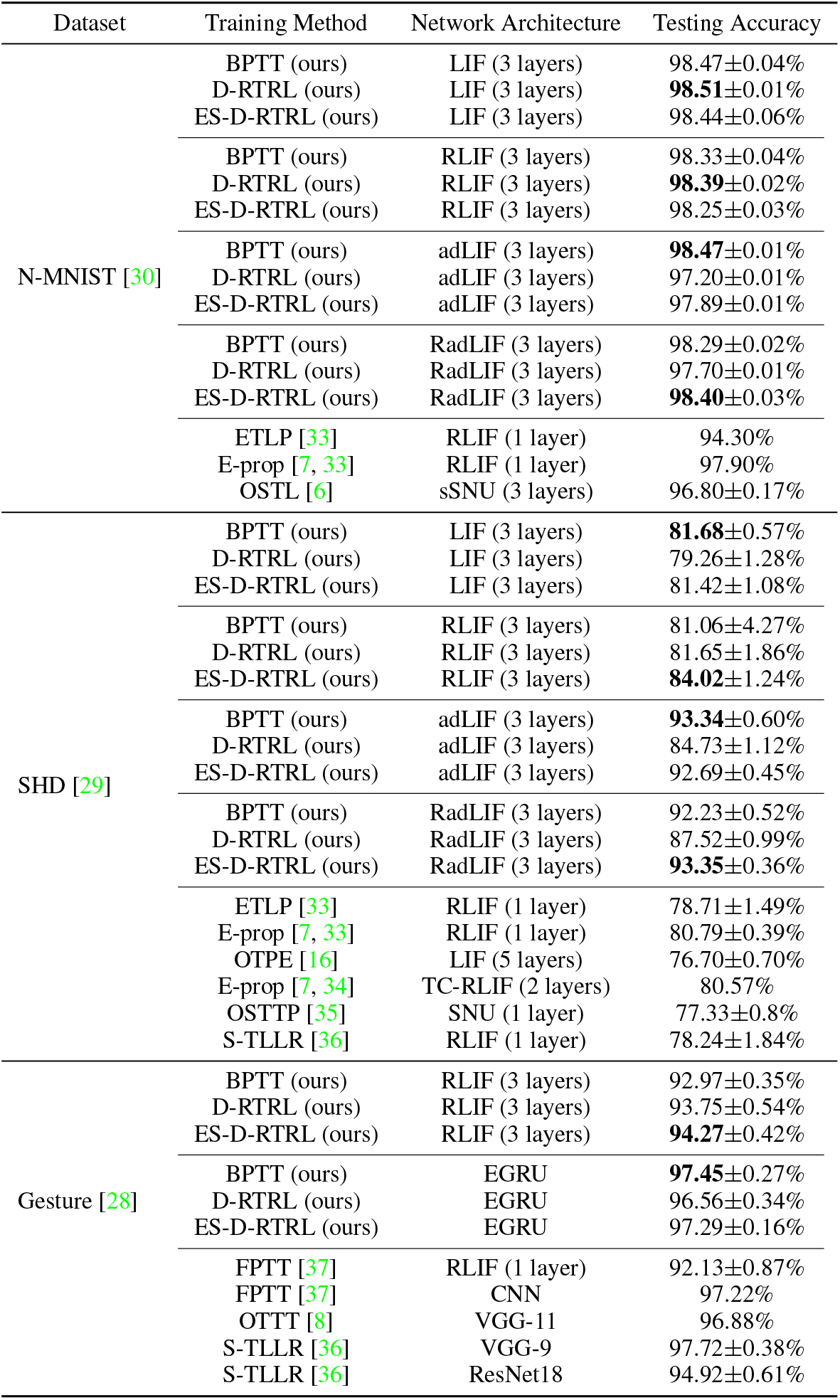
Comparison of training methods and network architectures across different datasets. Results show testing accuracy with standard deviation for various training approaches, including BPTT, D-RTRL, and ES-D-RTRL applied to different network architectures (LIF, RLIF, adLIF, RadLIF) on N-MNIST, SHD, and Gesture datasets. State-of-the-art online learning methods are included for comparison. The bolded data signifies the optimal performance under the specified control parameters.

We also compared our ES-D-RTRL algorithm against several state-of-the-art online learning algorithms, including OSTL [6], E-prop [7], ETLP [33], OSTTP [35], OTPE [16], S-TLLR [36], FPTT [37], and OTTT [8]. ES-D-RTRL consistently outperformed all competing methods across the evaluated datasets, even when using identical network architectures. Notably, on the SHD dataset, ES-D-RTRL achieved a performance improvement of over 4% compared to the best baseline. On the Gesture dataset, both the RLIF-based and CNN-based architectures trained with ES-D-RTRL showed comparable performance to those trained with existing online learning algorithms, demonstrating ES-D-RTRL has superior learning efficiency.

### Performance evaluations on brain simulation tasks using excitatory-inhibitory spiking networks

ES-D-RTRL is theoretically applicable to a broad spectrum of spiking networks, including biologically realistic circuits with complex neuronal dynamics. To illustrate this, we developed a biologically-informed spiking network (Fig. 5B) incorporating: four-variable generalized leaky integrate-and-fire (GIF) neurons [40] for complex neuronal dynamics, distinct excitatory and inhibitory (E/I) populations, conductance-based synapses [41], and sparse connectivity [42]. Our network comprised 800 GIF neurons, with a 4:1 excitatory-to-inhibitory ratio and a 10% random connection probability, each featuring conductance-based exponential synapse dynamics (Eq. S55). We trained this network on an evidence-accumulation task [43], where animals navigate a virtual linear track and encounter seven visual cues (Fig. 5A; SI. I). The task required animals to integrate left/right visual cues and, upon reaching a T-junction, select the corresponding direction. Correct decisions based on majority cue distribution were rewarded.

**Figure 5.**
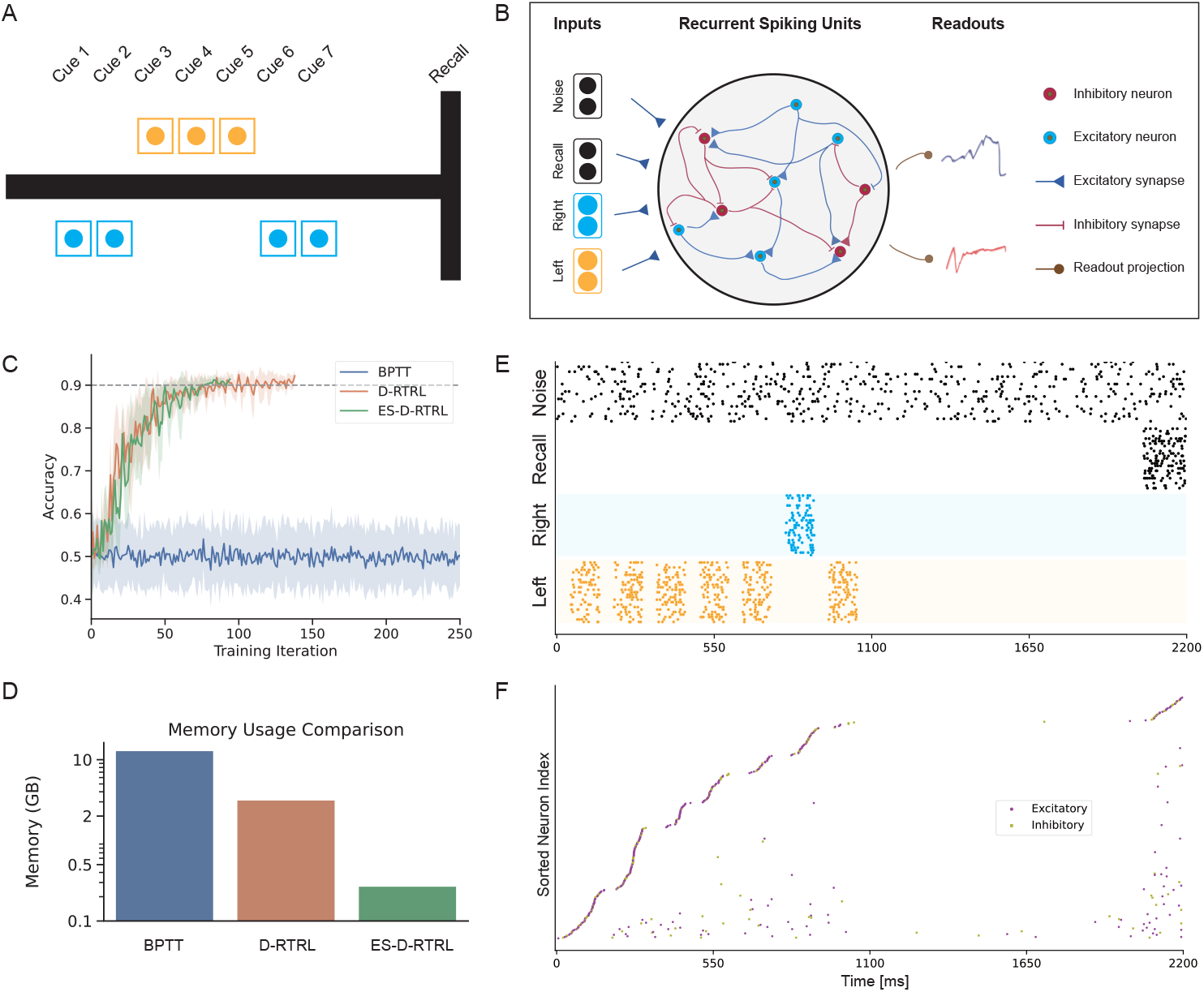
Training and analysis of biologically-informed excitatory-and-inhibitory network for cognitive task performance. **A**. Schematic of a virtual T-maze trial. Seven visual cues are presented (three left, four right), with the majority direction (right) indicating the correct choice for reward. **B**. Architecture of the recurrent excitatory-inhibitory spiking network implementing the T-maze task. The network transforms time-varying sensory inputs encoding cues and task rules into binary motor outputs representing left or right decisions. **C**. Testing accuracy comparison across training algorithms: BPTT, D-RTRL, and ES-D-RTRL. **D**. Device memory usage comparison during network training across algorithms. **E**. Example input spike trains for the evidence accumulation task, with time on the *x*-axis and neuron index on the *y*-axis. **F**. Network spiking activity following ES-D-RTRL training in response to the input pattern shown in **E**. Raster plot displays sorted recurrent network spikes after training completion.

Fig. 5C compares training performance for BPTT, D-RTRL, and ES-D-RTRL. Without surrogate gradients, BPTT failed to solve the task, whereas D-RTRL and ES-D-RTRL both achieved robust convergence. Despite the network’s relatively small neuron size, BPTT consumed over 10 GB of device memory and D-RTRL required more than 2 GB, due to their high memory complexity (Fig. 5D and Fig. 4I). In contrast, ES-D-RTRL sustained high performance while consuming less than 0.2 GB.

Post-training analysis of ES-D-RTRL network dynamics revealed highly selective, temporally precise spiking patterns. Fig. 5E shows input spike trains from a representative trial. During the evidence accumulation phase (0 - 1100 ms), neurons trained by ES-D-RTRL exhibited high selectivity to specific visual cues. Sequentially presented visual cues induced orderly transitions between transient and clustered neuron population activities. Neurons in each population predominantly utilized a time-to-first-spike coding strategy for sensory processing. Notably, within each selective population, neuronal responses to external stimuli displayed clear rank ordering, with neurons firing sequentially in response to stimuli (Fig. 5F). Remarkably, these neural dynamics closely align with neuronal activities observed in the posterior parietal cortex of mice performing similar evidence accumulation tasks (Fig. S17, adapted from Fig. 2a in [43]).

### Modeling *Drosophila* resting-state functional connectivity via whole-brain spiking network training

The linear-memory complexity of ES-D-RTRL provides a crucial advantage for training large-scale spiking networks. To demonstrate this, we trained a whole-brain spiking network of the *Drosophila* brain constrained by the FlyWire anatomical connectome [21], to reproduce both resting-state whole-brain calcium imaging signals and their inter-regional functional connectivity [23, 24]. Although the FlyWire connectome [21] offers a detailed anatomical map of neuronal wiring, direct implementation of this connectivity in spiking network models [22] fails to accurately capture either the resting-state neural activity in each neuropil (Fig. 6G) or the cross-neuropil correlations (Fig. 6K) recorded in experiments. We therefore treated the connectome-based network as an architecture scaffold [22] and trained a low-rank synaptic weight to account for individual variability using our ES-D-RTRL algorithm.

**Figure 6.**
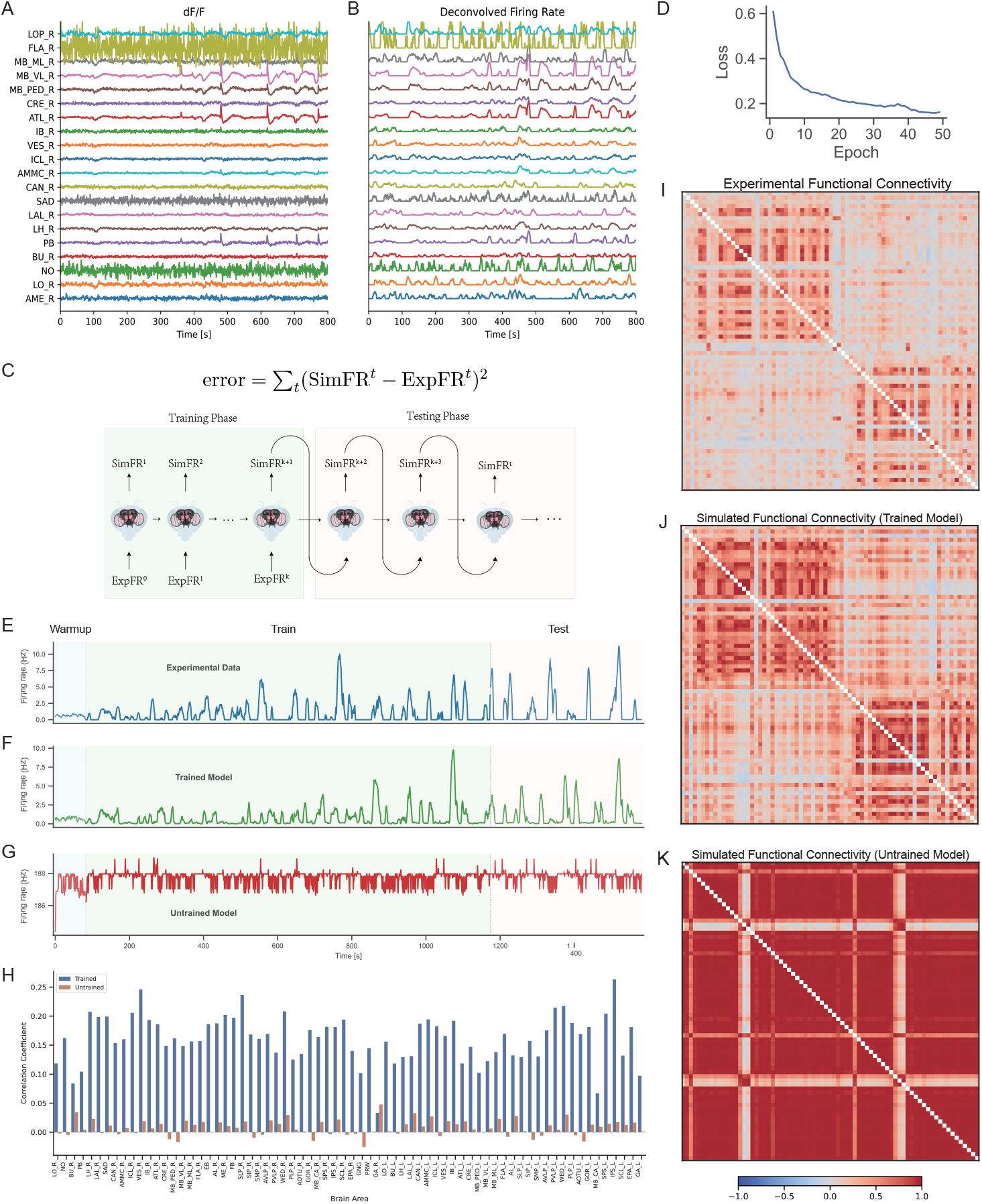
Fitting *Drosophila* resting-state neural activity by training whole-brain spiking networks. **A**. Representative ΔF*/*F traces from atlas ROIs in a single fly over 800 seconds, obtained from calcium imaging experiments [23, 24]. **B**. Deconvolved firing rates for the corresponding neuropils shown in **A. C**. Schematic of the training and testing paradigm. During the training phase, the model receives the experimental firing rate at time t (ExpFR^*t*^) and predicts the rate at time t + 1 (SimFR^*t*^) by minimizing the mean squared error. In the testing phase, the model autonomously generates sequential activations from its most recent prediction. **D**. Loss convergence of whole-brain neural activity fitting using the ES-D-RTRL algorithm. **E-G**. Firing rate dynamics for the Mushroom body neuropil: (**E**) experimental data, (**F**) outputs from the trained model, and (**G**) outputs from an untrained control. The initial `Warmup` phase (80 seconds) initializes network activity with experimental data. In the subsequent `Train` phase, the network generates activity at time *t* + 1 based on its prior output at time *t*. The `Test` phase evaluates the network’s ability to generate activity over unseen periods. **H**. Quantitative comparison of similarity between experimental data and model-generated outputs, contrasting trained versus untrained models. **I–K**. Functional connectivity matrices derived from (**I**) experimental recordings, (**J**) the trained whole-brain network, and (**K**) the untrained network, underscoring the efficacy of the network in capturing intrinsic connectivity patterns.

We first converted the 17 minutes of calcium signals recorded from each brain region (Fig. 6A) into deconvolved firing rates (Fig. 6B), and divided the entire dataset into a training set of 80% and a testing set of 20%. During training, the network received the experimental deconvolved firing rate at time *t* (ExpFR^*t*^) as input and predicted the firing rate at time *t* + 1 (SimFR^*t*+1^). The loss function was defined as the mean squared error between simulated and experimental firing rates (Fig. 6C). ES-D-RTRL then continuously adjusted synaptic weights online according to the error gradients, enabling the network to autonomously learn the complex spatiotemporal patterns underlying resting-state dynamics. This model consumes approximately 8.9 GB of GPU memory when using ES-D-RTRL, yet training with D-RTRL and BPTT exceeds the capacity of a single 32 GB GPU (Fig. S20). Fig. 6D shows the convergence of the loss function during the ES-D-RTRL training.

We evaluated the model’s dynamics in the mushroom body to assess training-induced improvements (Fig. 6E–G). In the warmup phase, we initialized whole-brain dynamics by feeding the first 84 seconds of experimental firing rates into the model; thereafter, with no external inputs, the model generated ongoing neural activity solely from its previous timestep’s neuropil state. Before training, the model failed to reproduce any experimentally observed spontaneous fluctuations (Fig. 6G); after training, however, its output not only exhibited activation patterns highly consistent with the experimental recordings (“Train” phase in Fig. 6F) but also sustained similar oscillation patterns on previously novel test data (“Test” phase in Fig. 6F). These results demonstrate the model’s ability to generalize to untrained neural dynamics. To quantify improvements across the entire *Drosophila* brain, we computed Pearson correlations between model-generated and experimental firing rates for each of 68 regions, both pre- and post-training. Training produced a marked uplift in regional fit, and statistical tests confirmed that nearly every region exhibited a significant post-training correlation increase (Fig. 6H).

We further compared whole-brain functional connectivity across experimental recordings, the un-trained model’s spontaneous activity, and the trained model’s output (Fig. 6I–K). Prior to training, neurons within each neuropil exhibited spontaneous fluctuations with similar activity patterns (Fig. S18), leading to high correlations across neuropils (Fig. 6K). After training, the model dynamics generated highly consistent functional connectivity strengths and patterns (Fig. 6J, Fig. S19) compared to experimental data, regardless of whether during the training or testing phases.

## Discussion

As next-generation neural networks [44], SNNs can benefit from the lessons learned in the development of ANNs [45]. The success of ANNs is largely attributed to the synergistic integration of the backpropagation algorithm and AD frameworks, in which backpropagation provides the theoretical foundation for ANN training, and AD frameworks ensure convenience in implementation by automating the differentiation process. In contrast, the current paradigms for SNN online learning have not yet achieved similar co-optimization between algorithms and frameworks, limiting SNNs from fully leveraging the theoretical advantages of online learning in real-world applications. In this work, we designed the *BrainScale* system, which simultaneously optimizes three critical dimensions to enable scalable online learning for SNNs:

1. **Model universality**: BrainScale introduces `AlignPre` and `AlignPost` abstractions to comprehensively support SNNs that capture essential characteristics of neural systems.
2. **Computational efficiency**: BrainScale leverages the inherent sparse and event-driven nature of SNNs to reduce the complexity of RTRL, achieving an online learning algorithm with memory complexity that scales linearly with the number of neurons.
3. **Engineering usability**: BrainScale introduces an online learning compiler that automatically adapts to user-defined SNN models, enabling end-to-end automation from mathematical modeling to hardware deployment.

Our comprehensive evaluations demonstrate that BrainScale delivers effective gradient approximation while substantially improving both memory efficiency and computational speed over existing approaches. BrainScale demonstrates robust performance across diverse applications, from neuromorphic datasets to cognitive tasks and *Drosophila* whole-brain neural activity fitting.

The ES-D-RTRL algorithm advances recent efforts to scale spiking networks [46, 47] by combining biological plausibility with linear-memory training efficiency. It builds on diagonal approximation strategies [7], exploiting the inherent sparsity of spiking activity—biological neurons typically fire at low rates (0.2–5 Hz) [48], with only 1–2% active at any given time in cortical circuits [49]. Crucially, ES-D-RTRL introduces a rank-one approximation based on the consistent sign of spike-based inputs—a biologically grounded yet previously untapped property that enables reduction of memory complexity from quadratic to linear while preserving learning dynamics.

Beyond efficiency, ES-D-RTRL offers insights into a fundamental question in computational neuroscience: What advantages do SNNs offer over conventional rate-based models? While the energy efficiency of event-driven computation is well established [3], our results reveal an additional benefit in the context of learning. We show that the dynamics of spiking neurons permit an accurate, online approximation of gradients with linear complexity, highlighting a structural advantage of SNNs for online learning.

Built on our BrainPy ecosystem for simulating general spiking networks [50, 26, 51], BrainScale extends this framework with automated online learning capabilities. Its automated online learning compiler enables researchers to concentrate on model design and scientific questions rather than low-level implementation, a crucial advantage in modern deep learning where rapid prototyping and hardware portability are essential for progress [18, 19]. In contrast, existing online learning approaches typically rely on predefined neuron and synapse models, limiting users to assembling networks from a fixed library of components [8, 36], which restricts the flexibility in customizing dynamics or connectivity. This rigidity also impedes crucial optimizations for efficient and scalable training, such as eligibility trace reuse and vectorized Jacobian computation (SI. L).

The scalability of BrainScale is demonstrated by our application to whole-brain *Drosophila* neural activity fitting. As neuromorphic computing enters a critical phase where scalability is essential [47], the linear memory scaling of ES-D-RTRL offers a practical and efficient solution for brain-scale simulations that were previously computationally intractable. Particularly, our *Drosophila* connectome model—containing more than 125,000 neurons and 50 million synaptic connections—poses substantial memory demands: conventional methods such as D-RTRL and BPTT require over 100 GB of memory at peak usage. In contrast, ES-D-RTRL completes the same task using only 8.9 GB (Fig. S20). This highlights ES-D-RTRL’s superior scalability and efficiency for large-scale network training.

While our results highlight several compelling advantages of BrainScale, a number of challenges and limitations remain open for future work. First, although the ES-D-RTRL algorithm has proven effective across multiple benchmarks, its gradient approximation, especially in networks with high firing rates or deeper recurrent layers, still exhibits dataset and architecture sensitivity. A deeper theoretical exploration of these approximations could help define tighter bounds for their applicability. Second, extending BrainScale to incorporate plasticity mechanisms [52] that capture additional facets of synaptic adaptation may further enhance its biological fidelity. Third, exploring hybrid approaches that combine the computational efficiency of ANN-SNN conversion methods [53] with online learning capabilities represents a promising research direction. While conversion methods typically excel in inference speed and energy efficiency, they currently lack the adaptive capacity for continuous learning that our framework provides. Developing techniques that leverage pre-trained ANN knowledge as initialization while maintaining online adaptation could potentially combine the best of both paradigms. Finally, while our compiler’s automated approach currently supports many hardware platforms, further integration with emerging neuromorphic hardware architectures [54, 47] could unlock even greater efficiency and energy savings, particularly in low-power, embedded applications.

## Data availability

The datasets used in this study are publicly available and open source. The N-MNIST dataset is freely available at https://www.garrickorchard.com/datasets/n-mnist [30] (accessed on June 20, 2025). The SHD dataset is publicly available at https://zenkelab.org/resources/spiking-heidelberg-datasets-shd [29] (accessed on June 20, 2025). The IBM DVS Gesture dataset can be downloaded from https://ibm.ent.box.com/s/3hiq58ww1pbbjrinh367ykfdf60xsfm8/folder/50167556794 [28] (accessed on June 20, 2025). The DMTS and evidence accumulation tasks are generated in this study and can be found in the publicly available GitHub repository https://github.com/chaobrain/brainscale-exp-for-snns. The whole-brain calcium imaging data of *Drosophila* can be obtained in https://doi.org/10.6084/m9.figshare.13349282 [24] (accessed on June 20, 2025).

## Code availability

BrainScale is distributed via the PyPI package index (https://pypi.org/project/brainscale) and publicly released on GitHub (https://github.com/chaobrain/brainscale) under the license of Apache License v2.0. Its documentation will be hosted on the free documentation hosting platform Read the Docs (https://brainscale.readthedocs.io/). BrainScale can be used in Windows, macOS, and Linux operating systems. The code to reproduce the experimental evaluations of Fig. 4, Fig. 5, and Table 1 is publicly available from the GitHub repository https://github.com/chaobrain/brainscale-exp-for-snns. The code to reproduce the modeling of Fig. 6 is publicly available from the GitHub repository https://github.com/chaobrain/brainscale_fitting_drosophila_whole_brain_activity.

## Acknowledgments

This work was supported by the Young Scientists Fund of the National Natural Science Foundation of China (No. 3240070449, CMW), the Science and Technology Innovation 2030-Brain Science and Brain-inspired Intelligence Project (No. 2021ZD0200204, SW), and the National Natural Science Foundation of China (No. T2421004, SW).

## Author contributions

Conceptualization and Methodology: CMW, XSD, JDJ, SW. Software and Investigation: CMW. Analysis: CMW, XSD, ZLJ, JDJ, XL. Theorem Proof: XSD, WCM. Visualization: CMW, XL, ZLJ. Writing: CMW, ZLJ, SW. Writing (Review & Editing): CWM, ZLJ, XSD, JDJ, XL, SW. Funding Acquisition: CMW, SW.

