## Supplementary material for "BrainScale, Enabling Scalable Online Learning in Spiking Neural Networks": SI

**D Proof of Theorem C.2** **14**

**E Proof of Theorem C.3** **15**

**F Experimental validations of Theorem C.3** **17**

**O Environment settings** **45**

**P Code examples of BrainScale programming interface** **46**

**Q Supplementary figures** **50**

**R Supplementary tables** **69**

### A Model abstraction in BrainScale

BrainScale supports the online learning of models constructed based on the `AlignPre` and `AlignPost` projection principles (please refer to BrainScale supports a wide range of spiking networks). These principles, previously utilized in our BrainPy simulator [1] to create efficient and standardized SNN models, are now formally defined here. Our formulation covers neuronal dynamics (SI. A.1), synaptic dynamics (SI. A.2), `AlignPre` projection (SI. A.3) and `AlignPost` projection (SI. A.4). We also provide examples of SNN models employing this abstraction in SI. B.

#### A.1 Neuronal dynamics

The dynamics of a neuron are regulated by a complex interplay of ionic channels, neurotransmitter receptors, and intracellular signaling cascades, which collectively govern the neuron's membrane potential and excitability [2]. Moreover, neuronal dynamics are influenced by synaptic inputs, and neuromodulatory substances that can modulate the neuron's responsiveness [2]. As a result of these intricate processes, individual neurons exhibit diverse and complex spiking behaviors, displaying highly variable firing patterns in response to identical step currents [3]. To capture this diversity, a range of spiking neuron models has been proposed, including detailed compartmental models, reduced compartmental models, and point neuron models [4].

In general, we denote the neuronal state as $\mathbf{v}$ and the occurrence of a neuronal spike as $\mathbf{z}$, with their dynamics governed by the following equations:

$$\frac{\mathrm{d}\mathbf{v}}{\mathrm{d}t} = f_n(\mathbf{v}, t, \mathbf{I}), \tag{S9}$$

$$\mathbf{z}^t = \mathcal{H}(\mathbf{v}^t - v_{\text{th}}), \tag{S10}$$

where $\mathbf{I}$ represents the external current, $f_n$ typically denotes an element-wise function describing the evolution of each neuronal state, $v_{\text{th}}$ is the threshold membrane potential for spike generation, and $\mathcal{H}$ represents the Heaviside function used to determine whether a spike is generated at the current time step. Note that we treat $\mathbf{z}$ as an intermediate variable rather than one of the hidden states.

***One-dimensional point neuron model:*** The simplest approach for describing neuronal excitability involves one-dimensional dynamical systems, which consist of a single variable. An example of such a system is the leaky integrate-and-fire (LIF) model [5], whose dynamics is given by:

$$\tau\frac{\mathrm{d}\mathbf{v}}{\mathrm{d}t} = -(\mathbf{v} - V_{\text{rest}}) + \mathbf{I}, \tag{S11}$$

where $\tau$ is the membrane time constant, and $V_{\text{rest}}$ is the resting potential. Whenever the membrane potential reaches the spike threshold $V_{\text{th}}$ (i.e., $\mathbf{v}_i(t) \geq V_{th}$), the neuron fires a spike, and the membrane potential is reset to the reset potential (i.e., $\mathbf{v}_i(t) \leftarrow V_{\text{reset}}$). Using the exponential Euler integration, the LIF neuron can be discretized as:

$$\bar{\mathbf{v}}^{t-1} = \mathbf{v}^{t-1} + \mathbf{z}^{t-1}(V_{\text{reset}} - \mathbf{v}^{t-1}), \tag{S12}$$

$$\mathbf{v}^t = e^{-\Delta t/\tau}\bar{\mathbf{v}}^{t-1} + (1 - e^{-\Delta t/\tau})(V_{\text{rest}} + \mathbf{I}^t). \tag{S13}$$

Due to its simplicity and computational efficiency, the LIF model is widely utilized in spiking network modeling but it cannot precisely model the subthreshold dynamics. To capture subthreshold fluctuations of the membrane potential more accurately, quadratic [6] or exponential [7] extensions of the LIF model can also be employed.

***Two-dimensional point neuron model:*** However, one-dimensional models are limited in their ability to reproduce certain significant features of single neuron dynamics, such as adaptation and bursting. Models with at least two-dimensional states are required to faithfully replicate all the neurocomputational characteristics of neurons. Prominent examples include the adaptive exponential integrate-and-fire (AdEx) model [8] and the Izhikevich neuron model [9]. In this work, we are using the adaptive LIF (ALIF) model [10], whose dynamics is given by:

$$\tau_{\mathbf{a}}\frac{\mathrm{d}\mathbf{a}}{\mathrm{d}t} = -\mathbf{a}, \tag{S14}$$

$$\tau\frac{d\mathbf{v}}{dt} = -\mathbf{v} + V_{\text{rest}} + \mathbf{a} + \mathbf{I}, \tag{S15}$$

where $\mathbf{a}$ denotes the internal adaption current and $\tau_{\mathbf{a}}$ is the time constant of the adaption current. When $\mathbf{v}^i$ of $i$-th neuron meets $V_{th}$, the model fires:

$$\mathbf{a}_i \leftarrow \mathbf{a}_i + A, \tag{S16}$$

$$\mathbf{v}_i \leftarrow V_{\text{reset}}, \tag{S17}$$

where $A$ is the spike-triggered current.

Using the exponential Euler integration, the ALIF neuron can be discretized as:

$$\bar{\mathbf{v}}^{t-1} = \mathbf{v}^{t-1} + \mathbf{z}^{t-1}(V_{\text{reset}} - \mathbf{v}^{t-1}), \tag{S18}$$

$$\mathbf{a}^t = e^{-\Delta t/\tau_{\mathbf{a}}}\mathbf{a}^{t-1} + \mathbf{z}^{t-1}A, \tag{S19}$$

$$\mathbf{v}^t = e^{-\Delta t/\tau}\bar{\mathbf{v}}^{t-1} + (1 - e^{-\Delta t/\tau})(V_{\text{rest}} + \mathbf{I}^t + \mathbf{a}^t). \tag{S20}$$

***Multi-compartment neuron model:*** Recently, dendritic mechanisms have been incorporated into simplified point-based neuron models [11] to account for important contributions of dendrites to neuronal integration and output, thereby enhancing the computational power of realistic neurons. These models employ phenomenological equations to describe dendritic compartments and spikes, and can be easily trained using gradient-based learning methods. Dendritic mechanisms operate across multiple timescales, ranging from a few milliseconds to hundreds of milliseconds, enabling complex computations such as coincidence detection, low-pass filtering, input segregation/amplification, parallel nonlinear processing, and logical operations [12–18, 13, 19, 20]. Recent SNN networks that incorporate similar dendritic mechanisms have demonstrated superior advantages in temporal integration [21] and long-term dependency learning [22].

##### A.2 Synaptic dynamics

Synaptic dynamics, on the other hand, describe the processes that occur at the synapses, which facilitate communication between neurons. Synaptic dynamics typically involve three processes: neurotransmitter release from the presynaptic neuron, binding of these neurotransmitters to receptors on the postsynaptic cell (which is further detailed in Section A.3 and A.4), and subsequent generation of postsynaptic excitatory or inhibitory currents.

**1. Presynaptic neurotransmitter releasing.** Firstly, synaptic kinetics are defined by presynaptic efficiency, such as the number of neurotransmitters released from the presynaptic cell [23, 24]. This phenomenon is known as short-term plasticity (STP), although it can be disregarded in modeling when the STP effect is weak. STP can take two forms: short-term facilitation (STF) and short-term depression (STD), which respectively increase and decrease synaptic efficiency with repeated presynaptic activity. In the BrainScale framework, the STP effect can be modeled using the `AlignPre` projection, where each neuron has a corresponding STP state.

***STD model:*** The STD effect is modeled using a normalized variable $\mathbf{x}$ (where each element $0 \leq \mathbf{x}_i \leq 1$), representing the fraction of available resources after neurotransmitter depletion [25]:

$$\frac{\mathrm{d}\mathbf{x}}{\mathrm{d}t} = \frac{1-\mathbf{x}}{\tau_d} - U\mathbf{x}\delta(t - t^{\text{pre}}), \tag{S21}$$

where $U$ is the fraction of resources used per spike, $\tau_d$ is the time constant for recovery of synaptic vesicles, $t^{\text{pre}}$ is the arrival time of the presynaptic spike, and $\delta(*)$ is the Dirac delta function representing spike occurrences. For numerical integration, the exponential Euler method can be used to accurately solve the STD model:

$$\mathbf{x}^t = e^{-\Delta t/\tau_d}\mathbf{x}^{t-1} + (1 - e^{-\Delta t/\tau_d}) - U\mathbf{x}^{t-1}\mathbf{z}^{\text{pre},t-t_d}, \tag{S22}$$

where $\Delta t$ is the numerical integration time step and $t_d$ is the spike transmission delay.

***STP model:*** The STF effect is modeled using a utilization parameter $\mathbf{u}$, representing the fraction of available resources ready for use (release probability). The dynamics of $\mathbf{u}$ and $\mathbf{x}$ are governed by the following equations:

$$\frac{\mathrm{d}\mathbf{u}}{\mathrm{d}t} = -\frac{\mathbf{u}}{\tau_f} + U(1 - \mathbf{u}^-)\delta(t - t^{\text{pre}}), \tag{S23}$$

$$\frac{\mathrm{d}\mathbf{x}}{\mathrm{d}t} = \frac{1-\mathbf{x}}{\tau_d} - \mathbf{u}^+\mathbf{x}^-\delta(t - t^{\text{pre}}), \tag{S24}$$

where $\mathbf{u}^-$ and $\mathbf{x}^-$ are the corresponding variables just before the arrival of the spike, and $\mathbf{u}^+$ refers to the moment just after the spike. In the parameter regime of $\tau_d \gg \tau_f$ and large $U$, an initial spike leads to a significant drop in $\mathbf{x}$ that takes a long time to recover, resulting in STD dominance. In the regime of $\tau_f \gg \tau_d$ and small $U$, the synaptic efficacy gradually increases with spikes, leading to STF dominance. Using the exponential Euler integration method, the STP model can be discretized as follows:

$$\mathbf{u}^t = e^{-\Delta t/\tau_f}\mathbf{u}^{t-1} + U(1-\mathbf{u}^{t-1})\mathbf{z}^{\mathrm{pre},t-t_d}, \tag{S25}$$

$$\mathbf{x}^t = e^{-\Delta t/\tau_d}\mathbf{x}^{t-1} + (1-e^{-\Delta t/\tau_d}) - \mathbf{u}^t\mathbf{x}^{t-1}\mathbf{z}^{\mathrm{pre},t-t_d}. \tag{S26}$$

**2. Postsynaptic neurotransmitter binding.** Following the neurotransmitter release, synaptic efficiency is commonly characterized by the postsynaptic neurotransmitter binding ratio. This binding is denoted as the synaptic states $\mathbf{g}$, and their dynamics are governed by the equation:

$$\frac{\mathrm{d}\mathbf{g}}{\mathrm{d}t} = f_g(\mathbf{g}, t), \tag{S27}$$

where $f_g$ represents an element-wise function that specifies the evolution of each synaptic state based on neurotransmitter binding.

The specific formulation for modeling postsynaptic neurotransmitter binding is not provided in this section since the modeling approaches differ between the `AlignPre` and `AlignPost` projections in BrainScale. For detailed information regarding neurotransmitter binding, please refer to SI. A.3 and SI. A.4.

**3. Postsynaptic current generation.** Finally, once the postsynaptic conductance is obtained, the changes in current at the postsynaptic site can be computed using Ohm's law. The postsynaptic conductance induced by synaptic connections is denoted as $\mathbf{g}^{\mathrm{post},t}$. Please refer to SI. A.3 and SI. A.4 for detailed information on how it is calculated in `AlignPre` and `AlignPost` projections, respectively.

***COBA:*** Based on Ohm's law, the current generated in the postsynaptic cells is expressed as the conductance-based output (COBA):

$$\mathbf{I}^{\mathrm{post},t} = \mathbf{g}^{\mathrm{post},t} \circ (E - \mathbf{v}^{\mathrm{post},t}), \tag{S28}$$

where $E$ represents the reversal potential.

***CUBA:*** However, since the membrane voltage typically fluctuates around the resting membrane potential $v_{\mathrm{rest}}$, the above equation can be simplified as:

$$\mathbf{I}^{\mathrm{post},t} = \mathbf{g}^{\mathrm{post},t}(E - v_{\mathrm{rest}}) \propto \mathbf{g}^{\mathrm{post},t}, \tag{S29}$$

where $E - v_{\mathrm{rest}}$ is a constant that can be absorbed into the synaptic weight $\boldsymbol{\theta}$. This model is referred to as current-based output (CUBA).

***NMDA:*** In the case of NMDA synapses, the current flux is also influenced by magnesium. Given the synaptic conductance, the NMDA synapses generate the postsynaptic current using the following equation:

$$\mathbf{I}^{\mathrm{post},t} = \mathbf{g}^{\mathrm{post},t} \circ (E - \mathbf{v}^{\mathrm{post},t}) \circ g_\infty(\mathbf{v}^{\mathrm{post},t}), \tag{S30}$$

where the fraction of channels $g_\infty$ not blocked by magnesium can be fitted as $g_\infty(\mathbf{v}) = (1 + e^{-\alpha\mathbf{v}}\frac{[\mathrm{Mg}^{2+}]}{\beta})^{-1}$. Here, $[\mathrm{Mg}^{2+}]$ represents the extracellular magnesium concentration.

##### A.3 `AlignPre` synaptic projections

In the `AlignPre` projection (Figure 1C), the synaptic state $\mathbf{g}$ represents the changes in conductance on postsynaptic neurons induced by each presynaptic neuronal spike. Therefore, $\mathbf{g} \in \mathbb{R}^n$ has dimensions aligned with the presynaptic neurons. Two models are introduced below: the Alpha synapse and Markov synapse models.

***Alpha model:*** The Alpha synapse is a commonly used model, described analytically as $\mathbf{g}^t = \frac{t-t^{\mathrm{pre}}}{\tau_{\mathbf{g}}}\exp\left(-\frac{t-t^{\mathrm{pre}}}{\tau_{\mathbf{g}}}\right)$. However, this equation is challenging to implement directly. It can be transformed into differential forms for numerical integration:

$$\frac{d\mathbf{g}}{dt} = -\frac{\mathbf{g}}{\tau_{\mathbf{g}}} + \frac{\mathbf{s}}{\tau_{\mathbf{g}}}, \tag{S31}$$

$$\frac{d\mathbf{s}}{dt} = -\frac{\mathbf{s}}{\tau_{\mathbf{g}}} + \delta\left(t - t^{\text{pre}}\right), \tag{S32}$$

where $\mathbf{s}$ is an intermediate variable. By employing the exponential Euler integration method, the Alpha synapse model can be discretized as follows:

$$\mathbf{s}^t = e^{-\Delta t/\tau_{\mathbf{g}}}\mathbf{s}^{t-1} + \mathbf{z}^{\text{pre},t-t_d}, \tag{S33}$$

$$\mathbf{g}^t = e^{-\Delta t/\tau_{\mathbf{g}}}\mathbf{g}^{t-1} + (1 - e^{-\Delta t/\tau_{\mathbf{g}}})\mathbf{s}^t. \tag{S34}$$

***Synaptic interaction:*** The aforementioned `AlignPre` synaptic models describe how conductance changes on postsynaptic neurons are driven by each presynaptic neuron. In order to combine the effects of all presynaptic connections onto a single postsynaptic neuron, we need to compute the synaptic interaction. Typically, a linear transformation is used to calculate the aggregated postsynaptic conductance:

$$\mathbf{g}^{\text{post},t} = \boldsymbol{\theta} \circledast \mathbf{g}^t, \tag{S35}$$

Here, $\boldsymbol{\theta} \in \mathbb{R}^{n\times m}$ represents the synaptic connectivity, which can be a dense or sparse matrix. It is worth noting that the transformation $\boldsymbol{\theta}$ can also be implemented as a convolution or other operations commonly used in deep learning.

By incorporating the STP effect, the computation of the postsynaptic conductance can be modified as follows:

$$\mathbf{g}^{\text{post},t} = \begin{cases} \boldsymbol{\theta} \circledast (\mathbf{u}^t \circ \mathbf{x}^t \circ \mathbf{g}^t), & \text{for STP,} \\ \boldsymbol{\theta} \circledast (\mathbf{x}^t \circ \mathbf{g}^t), & \text{for STD.} \end{cases} \tag{S36}$$

In the case of STP, the conductance is computed by element-wise multiplication of vectors $\mathbf{u}^t$, $\mathbf{x}^t$, and $\mathbf{g}^t$, followed by the linear transformation $\boldsymbol{\theta}$. On the other hand, in the case of STD, only vectors $\mathbf{x}^t$ and $\mathbf{g}^t$ are element-wise multiplied before applying the transformation $\boldsymbol{\theta}$.

##### A.4 `AlignPost` synaptic projections

In the `AlignPost` projection (Figure 1D), the synaptic state $\mathbf{g} \in \mathbb{R}^m$ represents the summarized conductance effect on the specific postsynaptic site. Unlike `AlignPre` projections, `AlignPost` models initially calculate the aggregated conductance changes based on the connected presynaptic inputs (refer to the next paragraph "Synaptic interaction"). Subsequently, the synaptic conductance evolves in response to the summarized conductance inputs using exponential-family synapse models. Notably, the `AlignPost` projection can only apply to exponential-family synapse models.

***Synaptic interaction:*** In contrast to the `AlignPre` projection, the synaptic interactions in `AlignPost` projections should be first computed. It transforms the presynaptic spikes $\mathbf{z}^{\text{pre},t-t_d}$ into postsynaptic conductance changes $\Delta\mathbf{g}^{\text{post}}$ using the following equation:

$$\Delta\mathbf{g}^{\text{post},t} = \boldsymbol{\theta} \circledast \mathbf{z}^{\text{pre},t-t_d}. \tag{S37}$$

Since $\mathbf{z}$ represents binary spikes, this transformation can be efficiently performed using event-driven operators provided in the BrainPy simulator [26, 1]. However, when the STP effect is considered (see SI. A.2), `AlignPost` projections calculate postsynaptic conductance changes as follows:

$$\Delta\mathbf{g}^{\text{post},t} = \begin{cases} \boldsymbol{\theta} \circledast (\mathbf{u}^t \circ \mathbf{x}^t \circ \mathbf{z}^{\text{pre},t-t_d}), & \text{for STP,} \\ \boldsymbol{\theta} \circledast (\mathbf{x}^t \circ \mathbf{z}^{\text{pre},t-t_d}), & \text{for STD,} \end{cases} \tag{S38}$$

where $\mathbf{u}$ and $\mathbf{x}$ are variables representing short-term facilitation and depression, respectively.

Once the postsynaptic conductance changes are computed, the postsynaptic conductance $\mathbf{g}^{\text{post},t}$ can be updated according to the following models.

***Exponential model:*** Assuming the binding of neurotransmitters is instantaneous, the single exponential model only models the decay phase of a neurotransmitter binding as $\mathbf{g}^{\mathrm{post},t} = \exp\left(-\left(t - t^{\mathrm{pre}}\right)/\tau_{\mathbf{g}}\right)$, where $\tau_{\mathbf{g}}$ is the decay time constant. This equation can be expressed in a differential equation form:

$$\frac{\mathrm{d}\mathbf{g}^{\mathrm{post}}}{\mathrm{d}t} = -\frac{\mathbf{g}^{\mathrm{post}}}{\tau_{\mathbf{g}}} + \sum_k \delta(t - t_k^{\mathrm{sp}}). \tag{S39}$$

Using the exponential Euler integration method, the exponential synapse model can be discretized as:

$$\mathbf{g}^{\mathrm{post},t} = e^{-\Delta t/\tau_{\mathbf{g}}} \mathbf{g}^{\mathrm{post},t-1} + \Delta \mathbf{g}^{\mathrm{post},t}. \tag{S40}$$

***Double exponential model:*** Considering both the decay and rice phases, the double exponential model reproduces the behavior of the neurotransmitter binding well: $\mathbf{g}^{\mathrm{post},t} = B\left(\exp\left(-\frac{t-t^{\mathrm{pre}}}{\tau_d}\right) - \exp\left(-\frac{t-t^{\mathrm{pre}}}{\tau_r}\right)\right)$, where $\tau_{\mathbf{g}}$ is the decay time constant, $\tau_r$ is the rise time constant, and $B = \frac{\tau_{\mathbf{g}}}{\tau_{\mathbf{g}} - \tau_r}\left(\frac{\tau_r}{\tau_{\mathbf{g}}}\right)^{\frac{\tau_r}{\tau_r - \tau_{\mathbf{g}}}}$ is the normalizing constant so that the maximal value of $\mathbf{g}^{\mathrm{post}}$ is 1 for a single spike. The double exponential model can be expressed as:

$$\mathbf{g}^{\mathrm{post},t} = B\left(\mathbf{s}_d^t - \mathbf{s}_r^t\right), \tag{S41}$$

$$\mathbf{s}_d^t = e^{-\Delta t/\tau_{\mathbf{g}}} \mathbf{s}_d^{t-1} + \Delta \mathbf{g}^{\mathrm{post},t}, \tag{S42}$$

$$\mathbf{s}_r^t = e^{-\Delta t/\tau_r} \mathbf{s}_r^{t-1} + \Delta \mathbf{g}^{\mathrm{post},t}. \tag{S43}$$

#### B Spiking networks evaluated in this work

In this section, we present the spiking networks evaluated in this study. We provide a detailed description of these networks through their discretized equations, which form the foundation of our analysis. A key focus is placed on the neuronal interaction (see SI. C), highlighted using the notation `neuronal interaction` with an under bracket. Other element-wise operations in the equations describe the intrinsic dynamics of individual neurons and synapses within the network. Additionally, we explicitly point out the concrete implementation of the hidden state $\mathbf{h}^t$ in Eq. 1, its diagonal elements of $\mathbf{D}^t$ and the hidden-to-weight diagonal matrix $\mathbf{D}_f^t$ in Eq. 4-8 for each network. Note that all partial derivatives in this section do not flow through the recurrent spikes $\mathbf{z}^{t-1}$, and mathematical notations are consistent with SI. A.

##### B.1 The network with LIF neurons and Delta synapses.

The network comprising LIF neurons interconnected via Delta synapses represents a foundational model in computational neuroscience and neuromorphic engineering. Its discretized dynamics can be represented as:

$$\begin{aligned}\bar{\mathbf{v}}^{t-1} &= \mathbf{v}^{t-1} + \mathbf{z}^{t-1}(V_{\text{reset}} - \mathbf{v}^{t-1}),\\ \mathbf{v}^t &= e^{-\Delta t/\tau}\bar{\mathbf{v}}^{t-1} + (1-e^{-\Delta t/\tau})(V_{\text{rest}} + \underbrace{\boldsymbol{\theta}^{\text{rec}} \circledast \mathbf{z}^{t-1} + \boldsymbol{\theta}^{\text{in}} \circledast \mathbf{z}^{\text{in},t}}_{\text{neuronal interaction}}),\end{aligned} \tag{S44}$$

where $\mathbf{z}^{\text{in},t} \in \mathbb{R}^I$ is the input spike at the time $t$, $\boldsymbol{\theta}^{\text{rec}} \in \mathbb{R}^{H\times H}$ is the recurrent weight, and $\boldsymbol{\theta}^{\text{in}} \in \mathbb{R}^{I\times H}$ is the input weight. The hidden state $\mathbf{h}^t$ in Eq. 1 corresponds to $\mathbf{v}^t$ in this network, $\mathbf{D}_{ii}^t = \partial \mathbf{v}_i^t/\partial \mathbf{v}_i^{t-1}$, and $\mathbf{D}_{f,ii}^t = \partial \mathbf{v}_i^t/\partial \mathbf{I}_i^t$.

##### B.2 The network with ALIF neurons and Delta synapses.

Its discretized dynamics is given by:

$$\begin{aligned}\bar{\mathbf{v}}^{t-1} &= \mathbf{v}^{t-1} + \mathbf{z}^{t-1}(V_{\text{reset}} - \mathbf{v}^{t-1}),\\ \mathbf{a}^t &= e^{-\Delta t/\tau_{\mathbf{a}}}\mathbf{a}^{t-1} + \mathbf{z}^{t-1}A,\\ \mathbf{v}^t &= e^{-\Delta t/\tau}\bar{\mathbf{v}}^{t-1} + (1-e^{-\Delta t/\tau})(V_{\text{rest}} + \mathbf{a}^t + \underbrace{\boldsymbol{\theta}^{\text{rec}} \circledast \mathbf{z}^{t-1} + \boldsymbol{\theta}^{\text{in}} \circledast \mathbf{z}^{\text{in},t}}_{\text{neuronal interaction}}),\end{aligned} \tag{S45}$$

where $\mathbf{a}^t \in \mathbb{R}^H$ is the internal adaption current,

$\mathbf{h}_i^t = \begin{pmatrix}\mathbf{v}_i^t\\ \mathbf{a}_i^t\end{pmatrix}$,

$\mathbf{D}_{ii}^t = \begin{pmatrix}\partial \mathbf{v}_i^t/\partial \mathbf{v}_i^{t-1} & \partial \mathbf{v}_i^t/\partial \mathbf{a}_i^{t-1}\\ \partial \mathbf{a}_i^t/\partial \mathbf{v}_i^{t-1} & \partial \mathbf{a}_i^t/\partial \mathbf{a}_i^{t-1}\end{pmatrix}$, and

$\mathbf{D}_{f,ii}^t = \begin{pmatrix}\partial \mathbf{v}_i^t/\partial \mathbf{I}_i^{t-1}\\ \partial \mathbf{a}_i^t/\partial \mathbf{I}_i^{t-1}\end{pmatrix}$.

##### B.3 The network with LIF neurons and current-based Exponential synapses.

The LIF neuron networks with current-based exponential synapses constitute another foundational model in computational neuroscience. Its discretized dynamics can be represented as:

$$\begin{aligned}\bar{\mathbf{v}}^{t-1} &= \mathbf{v}^{t-1} + \mathbf{z}^{t-1}(V_{\text{reset}} - \mathbf{v}^{t-1}),\\ \mathbf{g}^t &= e^{-\Delta t/\tau_{\mathbf{g}}}\mathbf{g}^{t-1} + \underbrace{\boldsymbol{\theta}^{\text{rec}} \circledast \mathbf{z}^{t-1} + \boldsymbol{\theta}^{\text{in}} \circledast \mathbf{z}^{\text{in},t}}_{\text{neuronal interaction}},\\ \mathbf{v}^t &= e^{-\Delta t/\tau}\bar{\mathbf{v}}^{t-1} + (1-e^{-\Delta t/\tau})(V_{\text{rest}} + \mathbf{g}^t),\end{aligned} \tag{S46}$$

where the synaptic current $\mathbf{g}^t \in \mathbb{R}^H$ is modeled using the exponential synapse with the CUBA output. In this network,

$\mathbf{h}_i^t = \begin{pmatrix}\mathbf{v}_i^t\\ \mathbf{g}_i^t\end{pmatrix}$,

$\mathbf{D}^t_{ii} = \begin{pmatrix} \partial\mathbf{v}^t_i/\partial\mathbf{v}^{t-1}_i & \partial\mathbf{v}^t_i/\partial\mathbf{g}^{t-1}_i \\ \partial\mathbf{g}^t_i/\partial\mathbf{v}^{t-1}_i & \partial\mathbf{g}^t_i/\partial\mathbf{g}^{t-1}_i \end{pmatrix}$, and

$\mathbf{D}^t_{f,ii} = \begin{pmatrix} \partial\mathbf{v}^t_i/\partial\mathbf{I}^{t-1}_i \\ \partial\mathbf{g}^t_i/\partial\mathbf{I}^{t-1}_i \end{pmatrix}$.

##### B.4 The network with ALIF neurons and current-based Exponential synapses.

The dynamics of the network can be discretized as:

$$\begin{aligned}
\bar{\mathbf{v}}^{t-1} &= \mathbf{v}^{t-1} + \mathbf{z}^{t-1}(V_{\text{reset}} - \mathbf{v}^{t-1}), \\
\mathbf{g}^t &= e^{-\Delta t/\tau_{\mathbf{g}}}\mathbf{g}^{t-1} + \underbrace{\boldsymbol{\theta}^{\text{rec}} \circledast \mathbf{z}^{t-1} + \boldsymbol{\theta}^{\text{in}} \circledast \mathbf{z}^{\text{in},t}}_{\text{neuronal interaction}}, \\
\mathbf{a}^t &= e^{-\Delta t/\tau_{\mathbf{a}}}\mathbf{a}^{t-1} + \mathbf{z}^{t-1}A, \\
\mathbf{v}^t &= e^{-\Delta t/\tau}\bar{\mathbf{v}}^{t-1} + (1 - e^{-\Delta t/\tau})(V_{\text{rest}} + \mathbf{a}^t + \mathbf{g}^t),
\end{aligned} \tag{S47}$$

where

$\mathbf{h}^t_i = \begin{pmatrix} \mathbf{v}^t_i \\ \mathbf{a}^t_i \\ \mathbf{g}^t_i \end{pmatrix}$,

$\mathbf{D}^t_{ii} = \begin{pmatrix} \partial\mathbf{v}^t_i/\partial\mathbf{v}^{t-1}_i & \partial\mathbf{v}^t_i/\partial\mathbf{a}^{t-1}_i & \partial\mathbf{v}^t_i/\partial\mathbf{g}^{t-1}_i \\ \partial\mathbf{a}^t_i/\partial\mathbf{v}^{t-1}_i & \partial\mathbf{a}^t_i/\partial\mathbf{a}^{t-1}_i & \partial\mathbf{a}^t_i/\partial\mathbf{g}^{t-1}_i \\ \partial\mathbf{g}^t_i/\partial\mathbf{v}^{t-1}_i & \partial\mathbf{g}^t_i/\partial\mathbf{a}^{t-1}_i & \partial\mathbf{g}^t_i/\partial\mathbf{g}^{t-1}_i \end{pmatrix}$, and

$\mathbf{D}^t_{f,ii} = \begin{pmatrix} \partial\mathbf{v}^t_i/\partial\mathbf{I}^{t-1}_i \\ \partial\mathbf{a}^t_i/\partial\mathbf{I}^{t-1}_i \\ \partial\mathbf{g}^t_i/\partial\mathbf{I}^{t-1}_i \end{pmatrix}$

##### B.5 The network with ALIF neurons and conductance-based Exponential synapses.

The dynamics of the network can be discretized as:

$$\begin{aligned}
\bar{\mathbf{v}}^{t-1} &= \mathbf{v}^{t-1} + \mathbf{z}^{t-1}(V_{\text{reset}} - \mathbf{v}^{t-1}), \\
\mathbf{g}^t &= e^{-\Delta t/\tau_{\mathbf{g}}}\mathbf{g}^{t-1} + \underbrace{\boldsymbol{\theta}^{\text{rec}} \circledast \mathbf{z}^{t-1} + \boldsymbol{\theta}^{\text{in}} \circledast \mathbf{z}^{\text{in},t}}_{\text{neuronal interaction}}, \\
\mathbf{a}^t &= e^{-\Delta t/\tau_{\mathbf{a}}}\mathbf{a}^{t-1} + \mathbf{z}^{t-1}A, \\
\mathbf{v}^t &= e^{-\Delta t(1+\mathbf{g}^t)/\tau}\bar{\mathbf{v}}^{t-1} + (1 - e^{-\Delta t(1+\mathbf{g}^t)/\tau})(V_{\text{rest}} + \mathbf{a}^t + E \circ \mathbf{g}^t),
\end{aligned} \tag{S48}$$

where

$\mathbf{h}^t_i = \begin{pmatrix} \mathbf{v}^t_i \\ \mathbf{a}^t_i \\ \mathbf{g}^t_i \end{pmatrix}$,

$\mathbf{D}^t_{ii} = \begin{pmatrix} \partial\mathbf{v}^t_i/\partial\mathbf{v}^{t-1}_i & \partial\mathbf{v}^t_i/\partial\mathbf{a}^{t-1}_i & \partial\mathbf{v}^t_i/\partial\mathbf{g}^{t-1}_i \\ \partial\mathbf{a}^t_i/\partial\mathbf{v}^{t-1}_i & \partial\mathbf{a}^t_i/\partial\mathbf{a}^{t-1}_i & \partial\mathbf{a}^t_i/\partial\mathbf{g}^{t-1}_i \\ \partial\mathbf{g}^t_i/\partial\mathbf{v}^{t-1}_i & \partial\mathbf{g}^t_i/\partial\mathbf{a}^{t-1}_i & \partial\mathbf{g}^t_i/\partial\mathbf{g}^{t-1}_i \end{pmatrix}$, and

$\mathbf{D}^t_{f,ii} = \begin{pmatrix} \partial\mathbf{v}^t_i/\partial\mathbf{I}^{t-1}_i \\ \partial\mathbf{a}^t_i/\partial\mathbf{I}^{t-1}_i \\ \partial\mathbf{g}^t_i/\partial\mathbf{I}^{t-1}_i \end{pmatrix}$.

##### B.6 The network with LIF neurons, STD, and current-based Exponential synapses.

The dynamics of the network can be discretized as:

$$\begin{aligned}
\bar{\mathbf{v}}^{t-1} &= \mathbf{v}^{t-1} + \mathbf{z}^{t-1}(V_{\text{reset}} - \mathbf{v}^{t-1}), \\
\mathbf{x}^t &= e^{-\Delta t/\tau_d}\mathbf{x}^{t-1} + (1 - e^{-\Delta t/\tau_d}) - U\mathbf{x}^{t-1}\mathbf{z}^{t-1}, \\
\mathbf{g}^t &= e^{-\Delta t/\tau_{\mathbf{g}}}\mathbf{g}^{t-1} + \underbrace{\boldsymbol{\theta}^{\text{rec}} \circledast (\mathbf{x}^t \circ \mathbf{z}^{t-1}) + \boldsymbol{\theta}^{\text{in}} \circledast \mathbf{x}\mathbf{z}^{\text{in},t}}_{\text{neuronal interaction}}, \\
\mathbf{v}^t &= e^{-\Delta t/\tau}\bar{\mathbf{v}}^{t-1} + (1 - e^{-\Delta t/\tau})(V_{\text{rest}} + \mathbf{g}^t),
\end{aligned} \tag{S49}$$

where $\mathbf{x}^t \in \mathbb{R}^H$ is the short-term depression variable for the recurrent spikes, and $\mathbf{xz}^{\text{in},t} \in \mathbb{R}^I$ is the short-term depression variable for the input spikes,

$$\mathbf{h}_i^t = \begin{pmatrix} \mathbf{v}_i^t \\ \mathbf{x}_i^t \\ \mathbf{g}_i^t \end{pmatrix},$$

$$\mathbf{D}_{ii}^t = \begin{pmatrix} \partial \mathbf{v}_i^t/\partial \mathbf{v}_i^{t-1} & \partial \mathbf{v}_i^t/\partial \mathbf{x}_i^{t-1} & \partial \mathbf{v}_i^t/\partial \mathbf{g}_i^{t-1} \\ \partial \mathbf{x}_i^t/\partial \mathbf{v}_i^{t-1} & \partial \mathbf{x}_i^t/\partial \mathbf{x}_i^{t-1} & \partial \mathbf{x}_i^t/\partial \mathbf{g}_i^{t-1} \\ \partial \mathbf{g}_i^t/\partial \mathbf{v}_i^{t-1} & \partial \mathbf{g}_i^t/\partial \mathbf{x}_i^{t-1} & \partial \mathbf{g}_i^t/\partial \mathbf{g}_i^{t-1} \end{pmatrix}, \text{ and}$$

$$\mathbf{D}_{f,ii}^t = \begin{pmatrix} \partial \mathbf{v}_i^t/\partial \mathbf{I}_i^{t-1} \\ \partial \mathbf{x}_i^t/\partial \mathbf{I}_i^{t-1} \\ \partial \mathbf{g}_i^t/\partial \mathbf{I}_i^{t-1} \end{pmatrix}$$

##### B.7 The network with ALIF neurons, STD, and current-based Exponential synapses.

The dynamics of the network can be discretized as:

$$\begin{aligned} \bar{\mathbf{v}}^{t-1} &= \mathbf{v}^{t-1} + \mathbf{z}^{t-1}(V_{\text{reset}} - \mathbf{v}^{t-1}), \\ \mathbf{x}^t &= e^{-\Delta t/\tau_d}\mathbf{x}^{t-1} + (1 - e^{-\Delta t/\tau_d}) - U\mathbf{x}^{t-1}\mathbf{z}^{t-1}, \\ \mathbf{g}^t &= e^{-\Delta t/\tau_{\mathbf{g}}}\mathbf{g}^{t-1} + \underbrace{\boldsymbol{\theta}^{\text{rec}} \circledast (\mathbf{x}^t \circ \mathbf{z}^{t-1}) + \boldsymbol{\theta}^{\text{in}} \circledast \mathbf{xz}^{\text{in},t}}_{\text{neuronal interaction}}, \\ \mathbf{a}^t &= e^{-\Delta t/\tau_{\mathbf{a}}}\mathbf{a}^{t-1} + \mathbf{z}^{t-1}A, \\ \mathbf{v}^t &= e^{-\Delta t/\tau}\bar{\mathbf{v}}^{t-1} + (1 - e^{-\Delta t/\tau})(V_{\text{rest}} + \mathbf{a}^t + \mathbf{g}^t), \end{aligned} \tag{S50}$$

where

$$\mathbf{h}_i^t = \begin{pmatrix} \mathbf{v}_i^t \\ \mathbf{x}_i^t, \\ \mathbf{a}_i^t, \\ \mathbf{g}_i^t \end{pmatrix},$$

$$\mathbf{D}_{ii}^t = \begin{pmatrix} \partial \mathbf{v}_i^t/\partial \mathbf{v}_i^{t-1} & \partial \mathbf{v}_i^t/\partial \mathbf{x}_i^{t-1} & \partial \mathbf{v}_i^t/\partial \mathbf{a}_i^{t-1} & \partial \mathbf{v}_i^t/\partial \mathbf{g}_i^{t-1} \\ \partial \mathbf{x}_i^t/\partial \mathbf{v}_i^{t-1} & \partial \mathbf{x}_i^t/\partial \mathbf{x}_i^{t-1} & \partial \mathbf{x}_i^t/\partial \mathbf{a}_i^{t-1} & \partial \mathbf{x}_i^t/\partial \mathbf{g}_i^{t-1} \\ \partial \mathbf{a}_i^t/\partial \mathbf{v}_i^{t-1} & \partial \mathbf{a}_i^t/\partial \mathbf{x}_i^{t-1} & \partial \mathbf{a}_i^t/\partial \mathbf{a}_i^{t-1} & \partial \mathbf{a}_i^t/\partial \mathbf{g}_i^{t-1} \\ \partial \mathbf{g}_i^t/\partial \mathbf{v}_i^{t-1} & \partial \mathbf{g}_i^t/\partial \mathbf{x}_i^{t-1} & \partial \mathbf{g}_i^t/\partial \mathbf{a}_i^{t-1} & \partial \mathbf{g}_i^t/\partial \mathbf{g}_i^{t-1} \end{pmatrix}, \text{ and}$$

$$\mathbf{D}_{f,ii}^t = \begin{pmatrix} \partial \mathbf{v}_i^t/\partial \mathbf{I}_i^{t-1} \\ \partial \mathbf{x}_i^t/\partial \mathbf{I}_i^{t-1} \\ \partial \mathbf{a}_i^t/\partial \mathbf{I}_i^{t-1} \\ \partial \mathbf{g}_i^t/\partial \mathbf{I}_i^{t-1} \end{pmatrix}.$$

##### B.8 The network with LIF neurons, STP, and current-based Exponential synapses.

The dynamics of the network can be discretized as:

$$\begin{aligned} \bar{\mathbf{v}}^{t-1} &= \mathbf{v}^{t-1} + \mathbf{z}^{t-1}(V_{\text{reset}} - \mathbf{v}^{t-1}), \\ \mathbf{u}^t &= e^{-\Delta t/\tau_f}\mathbf{u}^{t-1} + U(1 - \mathbf{u}^{t-1})\mathbf{z}^{t-1}, \\ \mathbf{x}^t &= e^{-\Delta t/\tau_d}\mathbf{x}^{t-1} + (1 - e^{-\Delta t/\tau_d}) - \mathbf{u}^t\mathbf{x}^{t-1}\mathbf{z}^{t-1}, \\ \mathbf{g}^t &= e^{-\Delta t/\tau_{\mathbf{g}}}\mathbf{g}^{t-1} + \underbrace{\boldsymbol{\theta}^{\text{rec}} \circledast (\mathbf{u}^t \circ \mathbf{x}^t \circ \mathbf{z}^{t-1}) + \boldsymbol{\theta}^{\text{in}} \circledast \mathbf{uxz}^{\text{in},t}}_{\text{neuronal interaction}}, \\ \mathbf{v}^t &= e^{-\Delta t/\tau}\bar{\mathbf{v}}^{t-1} + (1 - e^{-\Delta t/\tau})(V_{\text{rest}} + \mathbf{g}^t), \end{aligned} \tag{S51}$$

where $\mathbf{u}^t, \mathbf{x}^t \in \mathbb{R}^H$ are short-term plasticity variables for the recurrent spikes, and $\mathbf{uxz}^{\text{in},t} \in \mathbb{R}^I$ is the short-term plasticity variable for the input spikes,

$$\mathbf{h}_i^t = \begin{pmatrix} \mathbf{v}_i^t \\ \mathbf{u}_i^t \\ \mathbf{x}_i^t \\ \mathbf{g}_i^t \end{pmatrix},$$

$$\mathbf{D}_{ii}^t = \begin{pmatrix} \partial \mathbf{v}_i^t/\partial \mathbf{v}_i^{t-1} & \partial \mathbf{v}_i^t/\partial \mathbf{u}_i^{t-1} & \partial \mathbf{v}_i^t/\partial \mathbf{x}_i^{t-1} & \partial \mathbf{v}_i^t/\partial \mathbf{g}_i^{t-1} \\ \partial \mathbf{u}_i^t/\partial \mathbf{v}_i^{t-1} & \partial \mathbf{u}_i^t/\partial \mathbf{u}_i^{t-1} & \partial \mathbf{u}_i^t/\partial \mathbf{x}_i^{t-1} & \partial \mathbf{u}_i^t/\partial \mathbf{g}_i^{t-1} \\ \partial \mathbf{x}_i^t/\partial \mathbf{v}_i^{t-1} & \partial \mathbf{x}_i^t/\partial \mathbf{u}_i^{t-1} & \partial \mathbf{x}_i^t/\partial \mathbf{x}_i^{t-1} & \partial \mathbf{x}_i^t/\partial \mathbf{g}_i^{t-1} \\ \partial \mathbf{g}_i^t/\partial \mathbf{v}_i^{t-1} & \partial \mathbf{g}_i^t/\partial \mathbf{u}_i^{t-1} & \partial \mathbf{g}_i^t/\partial \mathbf{x}_i^{t-1} & \partial \mathbf{g}_i^t/\partial \mathbf{g}_i^{t-1} \end{pmatrix}, \text{ and}$$

$$\mathbf{D}_{f,ii}^t = \begin{pmatrix} \partial \mathbf{v}_i^t/\partial \mathbf{I}_i^{t-1} \\ \partial \mathbf{u}_i^t/\partial \mathbf{I}_i^{t-1} \\ \partial \mathbf{x}_i^t/\partial \mathbf{I}_i^{t-1} \\ \partial \mathbf{g}_i^t/\partial \mathbf{I}_i^{t-1} \end{pmatrix}.$$

##### B.9 The network with ALIF neurons, STP, and current-based Exponential synapses.

The dynamics of the network can be discretized as:

$$\begin{aligned} \bar{\mathbf{v}}^{t-1} &= \mathbf{v}^{t-1} + \mathbf{z}^{t-1}(V_{\text{reset}} - \mathbf{v}^{t-1}), \\ \mathbf{u}^t &= e^{-\Delta t/\tau_f}\mathbf{u}^{t-1} + U(1-\mathbf{u}^{t-1})\mathbf{z}^{t-1}, \\ \mathbf{x}^t &= e^{-\Delta t/\tau_d}\mathbf{x}^{t-1} + (1-e^{-\Delta t/\tau_d}) - \mathbf{u}^t\mathbf{x}^{t-1}\mathbf{z}^{t-1}, \\ \mathbf{g}^t &= e^{-\Delta t/\tau_{\mathbf{g}}}\mathbf{g}^{t-1} + \underbrace{\boldsymbol{\theta}^{\text{rec}} \circledast (\mathbf{u}^t \circ \mathbf{x}^t \circ \mathbf{z}^{t-1}) + \boldsymbol{\theta}^{\text{in}} \circledast \mathbf{u}\mathbf{x}\mathbf{z}^{\text{in},t}}_{\text{neuronal interaction}}, \\ \mathbf{a}^t &= e^{-\Delta t/\tau_{\mathbf{a}}}\mathbf{a}^{t-1} + \mathbf{z}^{t-1}A, \\ \mathbf{v}^t &= e^{-\Delta t/\tau}\bar{\mathbf{v}}^{t-1} + (1-e^{-\Delta t/\tau})(V_{\text{rest}} + \mathbf{a}^t + \mathbf{g}^t), \end{aligned} \tag{S52}$$

where

$$\mathbf{h}_i^t = \begin{pmatrix} \mathbf{v}_i^t \\ \mathbf{u}_i^t \\ \mathbf{x}_i^t \\ \mathbf{a}_i^t \\ \mathbf{g}_i^t \end{pmatrix},$$

$$\mathbf{D}_{ii}^t = \begin{pmatrix} \partial \mathbf{v}_i^t/\partial \mathbf{v}_i^{t-1} & \partial \mathbf{v}_i^t/\partial \mathbf{u}_i^{t-1} & \partial \mathbf{v}_i^t/\partial \mathbf{x}_i^{t-1} & \partial \mathbf{v}_i^t/\partial \mathbf{a}_i^{t-1} & \partial \mathbf{v}_i^t/\partial \mathbf{g}_i^{t-1} \\ \partial \mathbf{u}_i^t/\partial \mathbf{v}_i^{t-1} & \partial \mathbf{u}_i^t/\partial \mathbf{u}_i^{t-1} & \partial \mathbf{u}_i^t/\partial \mathbf{x}_i^{t-1} & \partial \mathbf{u}_i^t/\partial \mathbf{a}_i^{t-1} & \partial \mathbf{u}_i^t/\partial \mathbf{g}_i^{t-1} \\ \partial \mathbf{x}_i^t/\partial \mathbf{v}_i^{t-1} & \partial \mathbf{x}_i^t/\partial \mathbf{u}_i^{t-1} & \partial \mathbf{x}_i^t/\partial \mathbf{x}_i^{t-1} & \partial \mathbf{x}_i^t/\partial \mathbf{a}_i^{t-1} & \partial \mathbf{x}_i^t/\partial \mathbf{g}_i^{t-1} \\ \partial \mathbf{a}_i^t/\partial \mathbf{v}_i^{t-1} & \partial \mathbf{a}_i^t/\partial \mathbf{u}_i^{t-1} & \partial \mathbf{a}_i^t/\partial \mathbf{x}_i^{t-1} & \partial \mathbf{a}_i^t/\partial \mathbf{a}_i^{t-1} & \partial \mathbf{a}_i^t/\partial \mathbf{g}_i^{t-1} \\ \partial \mathbf{g}_i^t/\partial \mathbf{v}_i^{t-1} & \partial \mathbf{g}_i^t/\partial \mathbf{u}_i^{t-1} & \partial \mathbf{g}_i^t/\partial \mathbf{x}_i^{t-1} & \partial \mathbf{g}_i^t/\partial \mathbf{a}_i^{t-1} & \partial \mathbf{g}_i^t/\partial \mathbf{g}_i^{t-1} \end{pmatrix}, \text{ and}$$

$$\mathbf{D}_{f,ii}^t = \begin{pmatrix} \partial \mathbf{v}_i^t/\partial \mathbf{I}_i^{t-1} \\ \partial \mathbf{u}_i^t/\partial \mathbf{I}_i^{t-1} \\ \partial \mathbf{x}_i^t/\partial \mathbf{I}_i^{t-1} \\ \partial \mathbf{a}_i^t/\partial \mathbf{I}_i^{t-1} \\ \partial \mathbf{g}_i^t/\partial \mathbf{I}_i^{t-1} \end{pmatrix}.$$

##### B.10 The E/I network with GIF neurons and conductance-based Exponential synapses.

To implement the computational task of evidence accumulation (see Performance evaluations on brain simulation tasks using excitatory-inhibitory spiking networks), we design an excitatory-inhibitory (E-I) spiking neural network. Neurons in the network follow the generalized integrate-and-fire (GIF) dynamics, while synaptic interactions are modeled using exponential conductance-based synapses.

The GIF model, originally proposed by Mihalaş and Niebur [27], is an extension of the classical leaky integrate-and-fire (LIF) model. Unlike LIF, the GIF model incorporates a set of internal adaptation currents $I_j$, enabling the neuron to generate a rich repertoire of firing patterns through parameter tuning. The continuous-time dynamics of the model are given by:

$$\begin{aligned} \frac{\mathrm{d}I_j}{\mathrm{d}t} &= -k_j I_j, \\ \tau\frac{\mathrm{d}V}{\mathrm{d}t} &= -(V - V_{\text{rest}}) + R\sum_j I_j + RI, \\ \frac{\mathrm{d}V_{\text{th}}}{\mathrm{d}t} &= a(V - V_{\text{rest}}) - b(V_{\text{th}} - V_{\text{th},\infty}), \end{aligned} \tag{S53}$$

where $V$ is the membrane potential, $V_{\text{th}}$ is the dynamic firing threshold, $I$ is the synaptic input current, and $I_j$ are internal adaptation currents with decay rates $k_j$. When the membrane potential reaches the threshold ($V \geq V_{\text{th}}$), the neuron emits a spike, and the state is updated as follows:

$$\begin{aligned} I_j &\leftarrow R_j I_j + A_j, \\ V &\leftarrow V_{\text{reset}}, \\ V_{\text{th}} &\leftarrow \max(V_{\text{th,rest}}, V_{\text{th}}). \end{aligned} \tag{S54}$$

By appropriately configuring the parameters $k_j$, $R_j$, and $A_j$, the GIF model can reproduce a wide range of single-neuron spiking behaviors, including spike-frequency adaptation and bursting. In the original formulation [27], two adaptation currents ($I_1$, $I_2$) were used.

The discretized dynamics of our network implementation are described as follows:

$$\begin{aligned} \bar{\mathbf{v}}^{t-1} &= \mathbf{v}^{t-1} + \mathbf{z}^{t-1}(V_{\text{reset}} - \mathbf{v}^{t-1}), \\ \mathbf{I}_1^t &= e^{-\Delta t/\tau_1}\mathbf{I}_1^{t-1} + A_1 \cdot \mathbf{z}^{t-1}, \\ \mathbf{I}_2^t &= e^{-\Delta t/\tau_2}\mathbf{I}_2^{t-1} + A_2 \cdot \mathbf{z}^{t-1}, \\ \mathbf{V}_{\text{th}}^t &= e^{-\Delta tb}\mathbf{V}_{\text{th}}^{t-1} + (1 - e^{-\Delta tb})(a(V - V_{\text{rest}}) + V_{\text{th},\infty}), \\ \mathbf{g}_{\text{e}}^t &= e^{-\Delta t/\tau_{\text{e}}}\mathbf{g}_{\text{e}}^{t-1} + |\boldsymbol{\theta}_{\text{e}}^{\text{rec}}| \circledast \mathbf{z}_{\text{e}}^{t-1} + |\boldsymbol{\theta}^{\text{in}}| \circledast \mathbf{z}^{\text{in},t}, \\ \mathbf{g}_{\text{i}}^t &= e^{-\Delta t/\tau_{\text{i}}}\mathbf{g}_{\text{i}}^{t-1} - |\boldsymbol{\theta}_{\text{i}}^{\text{rec}}| \circledast \mathbf{z}_{\text{i}}^{t-1}, \\ \mathbf{d}^t &= \exp\left(-\frac{\Delta t(1 + \mathbf{g}_{\text{e}}^t + \mathbf{g}_{\text{i}}^t)}{\tau}\right), \\ \mathbf{v}^t &= \mathbf{d}^t \cdot \bar{\mathbf{v}}^{t-1} + (1 - \mathbf{d}^t) \cdot (V_{\text{rest}} + \mathbf{a}^t + E_{\text{e}} \circ \mathbf{g}_{\text{e}}^t + E_{\text{i}} \circ \mathbf{g}_{\text{i}}^t), \end{aligned} \tag{S55}$$

Here, $\tau_{\text{e}}$ and $\tau_{\text{i}}$ are the time constants of excitatory and inhibitory synapses, respectively; $E_{\text{e}}$ and $E_{\text{i}}$ are the synaptic reversal potentials; $\mathbf{g}_{\text{e}}^t$ and $\mathbf{g}_{\text{i}}^t \in \mathbb{R}^H$ denote the excitatory and inhibitory conductances; $\mathbf{z}_{\text{e}}^{t-1}$ and $\mathbf{z}_{\text{i}}^{t-1}$ are the spike trains from excitatory and inhibitory neurons; $\boldsymbol{\theta}_{\text{e}}^{\text{rec}}$ and $\boldsymbol{\theta}_{\text{i}}^{\text{rec}}$ are the recurrent weight matrices for E and I neurons; $\boldsymbol{\theta}^{\text{in}}$ encodes feedforward input weights; and $\mathbf{d}^t$ is a decay factor modulated by total conductance. The symbol $\circledast$ denotes convolution over spike trains, and $|\cdot|$ indicates element-wise absolute value, ensuring non-negativity of the weights.

In this network, the hidden state

$$\mathbf{h}_i^t = \begin{pmatrix} \mathbf{v}_i^t \\ \mathbf{I}_{1,i}^t \\ \mathbf{I}_{2,i}^t \\ \mathbf{V}_{\text{th},i}^t \\ \mathbf{g}_{\text{e},i}^t \\ \mathbf{g}_{\text{i},i}^t \end{pmatrix},$$

$$\mathbf{D}_{ii}^t = \begin{pmatrix} \partial\mathbf{v}_i^t/\partial\mathbf{v}_i^{t-1} & \partial\mathbf{v}_i^t/\partial\mathbf{I}_{1,i}^{t-1} & \partial\mathbf{v}_i^t/\partial\mathbf{I}_{2,i}^{t-1} & \partial\mathbf{v}_i^t/\partial\mathbf{V}_{\text{th},i}^{t-1} & \partial\mathbf{v}_i^t/\partial\mathbf{g}_{\text{e},i}^{t-1} & \partial\mathbf{v}_i^t/\partial\mathbf{g}_{\text{i},i}^{t-1} \\ \partial\mathbf{I}_{1,i}^t/\partial\mathbf{v}_i^{t-1} & \partial\mathbf{I}_{1,i}^t/\partial\mathbf{I}_{1,i}^{t-1} & \partial\mathbf{I}_{1,i}^t/\partial\mathbf{I}_{2,i}^{t-1} & \partial\mathbf{I}_{1,i}^t/\partial\mathbf{V}_{\text{th},i}^{t-1} & \partial\mathbf{I}_{1,i}^t/\partial\mathbf{g}_{\text{e},i}^{t-1} & \partial\mathbf{I}_{1,i}^t/\partial\mathbf{g}_{\text{i},i}^{t-1} \\ \partial\mathbf{I}_{2,i}^t/\partial\mathbf{v}_i^{t-1} & \partial\mathbf{I}_{2,i}^t/\partial\mathbf{I}_{1,i}^{t-1} & \partial\mathbf{I}_{2,i}^t/\partial\mathbf{I}_{2,i}^{t-1} & \partial\mathbf{I}_{2,i}^t/\partial\mathbf{V}_{\text{th},i}^{t-1} & \partial\mathbf{I}_{2,i}^t/\partial\mathbf{g}_{\text{e},i}^{t-1} & \partial\mathbf{I}_{2,i}^t/\partial\mathbf{g}_{\text{i},i}^{t-1} \\ \partial\mathbf{V}_{\text{th},i}^t/\partial\mathbf{v}_i^{t-1} & \partial\mathbf{V}_{\text{th},i}^t/\partial\mathbf{I}_{1,i}^{t-1} & \partial\mathbf{V}_{\text{th},i}^t/\partial\mathbf{I}_{2,i}^{t-1} & \partial\mathbf{V}_{\text{th},i}^t/\partial\mathbf{V}_{\text{th},i}^{t-1} & \partial\mathbf{V}_{\text{th},i}^t/\partial\mathbf{g}_{\text{e},i}^{t-1} & \partial\mathbf{V}_{\text{th},i}^t/\partial\mathbf{g}_{\text{i},i}^{t-1} \\ \partial\mathbf{g}_{\text{e},i}^t/\partial\mathbf{v}_i^{t-1} & \partial\mathbf{g}_{\text{e},i}^t/\partial\mathbf{I}_{1,i}^{t-1} & \partial\mathbf{g}_{\text{e},i}^t/\partial\mathbf{I}_{2,i}^{t-1} & \partial\mathbf{g}_{\text{e},i}^t/\partial\mathbf{V}_{\text{th},i}^{t-1} & \partial\mathbf{g}_{\text{e},i}^t/\partial\mathbf{g}_{\text{e},i}^{t-1} & \partial\mathbf{g}_{\text{e},i}^t/\partial\mathbf{g}_{\text{i},i}^{t-1} \\ \partial\mathbf{g}_{\text{i},i}^t/\partial\mathbf{v}_i^{t-1} & \partial\mathbf{g}_{\text{i},i}^t/\partial\mathbf{I}_{1,i}^{t-1} & \partial\mathbf{g}_{\text{i},i}^t/\partial\mathbf{I}_{2,i}^{t-1} & \partial\mathbf{g}_{\text{i},i}^t/\partial\mathbf{V}_{\text{th},i}^{t-1} & \partial\mathbf{g}_{\text{i},i}^t/\partial\mathbf{g}_{\text{e},i}^{t-1} & \partial\mathbf{g}_{\text{i},i}^t/\partial\mathbf{g}_{\text{i},i}^{t-1} \end{pmatrix}, \text{ and}$$

$$\mathbf{D}_{f,ii}^t = \begin{pmatrix} \partial\mathbf{v}_i^t/\partial\mathbf{I}_i^{t-1} \\ \partial\mathbf{I}_{1,i}^t/\partial\mathbf{I}_i^{t-1} \\ \partial\mathbf{I}_{2,i}^t/\partial\mathbf{I}_i^{t-1} \\ \partial\mathbf{I}_{\text{th},i}^t/\partial\mathbf{I}_i^{t-1} \\ \partial\mathbf{g}_{\text{e},i}^t/\partial\mathbf{I}_i^{t-1} \\ \partial\mathbf{g}_{\text{i},i}^t/\partial\mathbf{I}_i^{t-1} \end{pmatrix}.$$

##### B.11 The event-based gated recurrent unit.

EGRU [28] extends the GRU [29] by introducing two internal gates—an update gate $u$ and a reset gate $r$—that govern the evolution of its hidden state $y$, and a state variable $z$ that blends the current input $x$ with the gated memory. At time step $t$, the layer's dynamics are described by

$$\begin{aligned} u^{\langle t\rangle} &= \sigma\big(W_u\,[\,x^{\langle t\rangle},\, y^{\langle t-1\rangle}\,] + b_u\big),\\ r^{\langle t\rangle} &= \sigma\big(W_r\,[\,x^{\langle t\rangle},\, y^{\langle t-1\rangle}\,] + b_r\big),\\ z^{\langle t\rangle} &= g\big(W_z\,[\,x^{\langle t\rangle},\, r^{\langle t\rangle} \odot y^{\langle t-1\rangle}\,] + b_z\big),\\ y^{\langle t\rangle} &= u^{\langle t\rangle} \odot z^{\langle t\rangle} \;+\; \big(1 - u^{\langle t\rangle}\big) \odot y^{\langle t-1\rangle}, \end{aligned} \tag{S56}$$

where each $W$ and $b$ are the layer's weights and biases, $\sigma(\cdot)$ is the sigmoid activation, $g(\cdot)$ a pointwise nonlinearity (often $\tanh$), $\odot$ denotes element-wise multiplication, and $[\cdot,\cdot]$ concatenates two vectors.

To make the hidden state event-driven, EGRU replaces the continuous output $y_i^{\langle t\rangle}$ with a thresholded, spike-like variable. An auxiliary state $c_i^{\langle t\rangle}$ accumulates the gated updates, and whenever it exceeds a neuron-specific threshold $\vartheta_i$, an "event" is emitted and the state is reset:

$$\begin{aligned} y_i^{\langle t\rangle} &= c_i^{\langle t\rangle}\, H\big(c_i^{\langle t\rangle} - \vartheta_i\big),\\ c_i^{\langle t\rangle} &= u_i^{\langle t\rangle}\, z_i^{\langle t\rangle} \;+\; \big(1 - u_i^{\langle t\rangle}\big)\, c_i^{\langle t-1\rangle} \;-\; y_i^{\langle t-1\rangle}, \end{aligned} \tag{S57}$$

where $H(\cdot)$ is the Heaviside step function. In this way, each hidden unit only produces a nonzero output when its membrane potential $c_i$ crosses the threshold, then immediately clears, yielding a sparse, event-based sequence of activations.

#### C SNN inherent properties

This section offers a formal and detailed description of the intrinsic properties of SNNs. While some content overlaps with the section BrainScale achieves linear-memory online learning algorithm, this segment provides a more comprehensive explanation and employs rigorous mathematical formalism to elucidate the concepts.

##### C.1 Property 1: Intrinsic neuronal dynamics dominate the hidden Jacobian

A typical SNN model exhibits two types of recurrent connections. The first type comprises $\mathbf{D}^t$ recurrent spiking connections that propagate action potentials between neurons (see $\circledast$ in Eq. 1 and examples in SI. B). The second type $\mathbf{J}^t$ consists of intrinsic neuronal dynamics, such as leaky terms in neuronal and synaptic processes (see $f$ Eq. 1 and examples in SI. B). As a result, the hidden Jacobian is composed of two parts: $\frac{\partial \mathbf{h}^t}{\partial \mathbf{h}^{t-1}} = \mathbf{D}^t + \mathbf{J}^t$, where $\mathbf{D}^t \in \mathbb{R}^{H \times H}$ is a diagonal matrix[1] resulted from the element-wise intrinsic dynamics, and $\mathbf{J}^t$ is the square matrix derived from the recurrent spiking connections.

A defining characteristic of SNNs is their sparse activation patterns, with neurons firing at relatively low frequencies. Empirical studies have shown that biological neurons typically emit between 0.2 and 5 spikes per second [30], with only 1-2% of neurons in a cortical network active at any given moment [31, 32]. This fact leads to the hypothesis that $\mathbf{J}^t$ exerts limited influence and can be ignored in the hidden Jacobian, since $\mathbf{J}^t$ is highly sparse in both space and time. Earlier practices, such as E-prop [33] and RFLO [34], have also demonstrated that neglecting the gradient flow through recurrent spiking connections in specific neuronal dynamics does not significantly hinder learning performance. To further validate this hypothesis, we conducted empirical evaluations across various SNN architectures with different neuronal and synaptic dynamics (SI. B). Our findings consistently indicated that the hidden Jacobian of most SNN models is predominantly shaped by the diagonal Jacobian $\mathbf{D}^t$ induced by intrinsic dynamics. The diagonal approximation demonstrates a remarkably high cosine similarity with the full hidden Jacobian (see Fig. 2B). Notably, under biologically plausible firing regimes (fraction of spiking neurons < 10%), the cosine similarity exceeds 99%.

**Proposition C.1.** *In sparsely activated SNN models, the hidden Jacobian $\frac{\partial \mathbf{h}^t}{\partial \mathbf{h}^{t-1}}$ is dominated by the diagonal Jacobian $\mathbf{D}^t$ induced by intrinsic neural dynamics. This can be expressed as:*

$$\frac{\partial \mathbf{h}^t}{\partial \mathbf{h}^{t-1}} \approx \mathbf{D}^t. \tag{S58}$$

##### C.2 Property 2: Physical connections yield decomposable hidden-to-weight Jacobian

Biological SNNs are characterized by physically constrained connections, reflecting the actual synaptic formations between pre- and postsynaptic neurons. Specifically, in `AlignPre` and `AlignPost` abstractions, this structural property can be mathematically expressed as a parameterized transformation from inputs to hidden vectors, followed by the element-wise activation applied to the resulting hidden vectors (see Eq. 1). Such mathematical operation leads to an important implication: the hidden-to-weight Jacobian $\frac{\partial \mathbf{h}^t}{\partial \boldsymbol{\theta}^t}$, which quantifies the sensitivity of the network's output to changes in synaptic weights, can be decomposed into a Kronecker product of a vector and a diagonal matrix (see Theorem C.2 and its proof in SI. D; this observation has also been illustrated in [35] for rate-based recurrent neural networks).

**Theorem C.2.** *Given Eq. 1, let $\mathbf{I}^t = \boldsymbol{\theta} \circledast \mathbf{x}^t$, then the gradient of $\mathbf{h}^t$ with respect to $\boldsymbol{\theta}$ at time $t$ is*

$$\frac{\partial \mathbf{h}^t}{\partial \boldsymbol{\theta}^t} = \mathbf{D}_f^t \otimes \mathbf{x}^t, \tag{S59}$$

*where $\mathbf{D}_f^t$ is a diagonal matrix[2] with $\frac{\partial \mathbf{h}_i^t}{\partial \mathbf{I}_i^t}$ as its diagonal entry $\mathbf{D}_{f,ii}^t$, and $\otimes$ is the Kronecker product.*

[1] $\mathbf{D}^t$ is actually a block diagonal matrix of dimensions $\mathbb{R}^{Hd \times Hd}$, consisting of $H$-by-$H$ blocks. Each diagonal block, $\mathbf{D}_{ii}^t$, is a $d$-by-$d$ square matrix, where $d$ is the number of state variables aligned to neuron $i$. For examples, refer to SI. B. Without loss of generality, all subsequent derivations assume $d = 1$, making $\mathbf{D}^t$ a diagonal matrix of dimensions $\mathbb{R}^{H \times H}$.

[2] $\mathbf{D}_f^t$ is actually a block diagonal matrix of dimensions $\mathbb{R}^{Hd \times H}$, consisting of $H$-by-$H$ blocks. Each diagonal block, $\mathbf{D}_{f,ii}^t$, is a $d$-by-1 matrix. For example, refer to SI. B. With loss of generality, all subsequent derivations assume $d = 1$, making $\mathbf{D}_f^t$ a diagonal matrix of dimensions $\mathbb{R}^{H \times H}$.

##### C.3 Property 3: Sign-preserving neuronal outputs induce low-rank structure

Another unique characteristic of SNN models is that the outputs of each neuron, corresponding to inputs $\mathbf{x}^t$, maintain a consistent sign across all time steps. For `AlignPost` projections, $\mathbf{x}$ denotes the binary spike vector (see Supplementary Eqs. S37-S38); while for `AlignPre` projections, $\mathbf{x}^t$ denotes the synaptic conductance (see Supplementary Eqs. S35-S36). In both cases, the elements in $\mathbf{x}^t$ are sign-preserving and consistently positive across all time steps. Additionally, for the monotonous activation function $f$, its derivative $\mathbf{D}_f^t$ also maintains a consistent sign throughout all times. This distinctive property leads to an important mathematical implication: the summation of multiple rank-one matrices can be approximated by the rank-one outer product of the summed vectors. Theorem C.3 formalizes this concept, with its theoretical proof presented in SI. E and empirical validation on various SNN dynamics using real-world neuromorphic datasets provided in SI. F.

**Theorem C.3.** *Let $\mathbf{a}_i \in \mathbb{R}^m$ and $\mathbf{b}_i \in \mathbb{R}^n$ be vectors where all elements in $\mathbf{a}_i$ and $\mathbf{b}_i$ are of the same sign (i.e., they are all positive or negative). Then, the cosine similarity between $\sum_{i=1}^r \mathbf{a}_i \otimes \mathbf{b}_i$ and $\left(\frac{1}{r}\sum_{i=1}^r \mathbf{a}_i\right) \otimes \sum_{i=1}^r \mathbf{b}_i$ approaches 1 as $r \to \infty$:*

$$\lim_{r\to\infty} \cos\left(\sum_{i=1}^r \mathbf{a}_i \otimes \mathbf{b}_i, \left(\frac{1}{r}\sum_{i=1}^r \mathbf{a}_i\right) \otimes \sum_{i=1}^r \mathbf{b}_i\right) \to 1. \tag{S60}$$

#### D Proof of Theorem C.2

*Proof.* Given Eq. 1, using the chain rule, we can write the gradient as:

$$\frac{\partial \mathbf{h}}{\partial \boldsymbol{\theta}} = \frac{\partial \mathbf{h}}{\partial \hat{\mathbf{h}}} \frac{\partial \hat{\mathbf{h}}}{\partial \boldsymbol{\theta}}$$

where:

- $\mathbf{h} = f(\hat{\mathbf{h}}) \in \mathbb{R}^H$ by definition of the mapping
- $\mathbf{I} = \boldsymbol{\theta} \circledast \mathbf{x} \in \mathbb{R}^H$ is the pre-activation

Calculating each component:

- $\frac{\partial \mathbf{h}_i}{\partial \mathbf{I}_i} = f'(\mathbf{I}_i)$, $\frac{\partial \mathbf{h}_i}{\partial \mathbf{I}_j} = 0$ $(i \neq j)$, since $f$ is an element-wise activation function. Therefore, $\frac{\partial \mathbf{h}}{\partial \mathbf{I}} = \mathbf{D}_f \in \mathbb{R}^{H \times H}$, where $\mathbf{D}_f$ is a diagonal matrix, and $D_{f,ii} = f'(\mathbf{I}_i)$ by the derivative of $f$.
- $\frac{\partial \mathbf{I}_j}{\partial \boldsymbol{\theta}_{ij}} = \mathbf{x}_i$. In the vector form, $\frac{\partial \mathbf{I}}{\partial \boldsymbol{\theta}_{ij}} = \mathbf{x}$ by the definition of matrix multiplication.

Substituting back into the chain rule:

$$\frac{\partial \mathbf{h}_j}{\partial \boldsymbol{\theta}_{ij}} = f'(\mathbf{I}_j) \frac{\partial \mathbf{I}_j}{\partial \boldsymbol{\theta}_{ij}} = f'(\mathbf{I}_j)\mathbf{x}_i$$

In the vector form, we got

$$\frac{\partial \mathbf{h}}{\partial \boldsymbol{\theta}} = \mathbf{D}_f \otimes \mathbf{x}$$

where $\otimes$ is the outer product.

□

#### E Proof of Theorem C.3

**Theorem E.1.** *Let $\mathbf{a}_i \in \mathbb{R}^m$ and $\mathbf{b}_i \in \mathbb{R}^n$ be vectors. Elements in $\mathbf{a}_i$ are sampled from the distribution with mean $\mathbf{a}$ and variance $\sigma_{\mathbf{a}}^2$, i.e., $\mathbf{a}_i = \mathbf{a} + \sigma_{\mathbf{a}}\boldsymbol{\xi}_i$, $\langle \boldsymbol{\xi}_i \rangle = \mathbf{0}$, $\langle \boldsymbol{\xi}_i^T \boldsymbol{\xi}_j \rangle = m\delta_{i,j}$. Elements in $\mathbf{b}_i$ are sampled from the distribution with mean $\mathbf{b}$ and variance $\sigma_{\mathbf{b}}^2$, i.e., $\mathbf{b}_i = \mathbf{b} + \sigma_{\mathbf{b}}\boldsymbol{\xi}_i$, $\langle \boldsymbol{\xi}_i \rangle = \mathbf{0}$, $\langle \boldsymbol{\xi}_i^T \boldsymbol{\xi}_j \rangle = n\delta_{i,j}$. Then, as $r \to \infty$, the cosine similarity between $\frac{1}{r}\sum_{i=1}^r \mathbf{a}_i \otimes \mathbf{b}_i$ and $\left(\frac{1}{r}\sum_{i=1}^r \mathbf{a}_i\right) \otimes \left(\frac{1}{r}\sum_{i=1}^r \mathbf{b}_i\right)$ has the form of:*

$$\begin{aligned}
&\lim_{r\to\infty} \cos\left(\frac{1}{r}\sum_{i=1}^r \mathbf{a}_i \otimes \mathbf{b}_i, \left(\frac{1}{r}\sum_{i=1}^r \mathbf{a}_i\right) \otimes \left(\frac{1}{r}\sum_{i=1}^r \mathbf{b}_i\right)\right) \\
&\approx \sqrt{1 - \frac{1}{r r_{\mathbf{a}} r_{\mathbf{b}} + r_{\mathbf{a}} + r_{\mathbf{b}} + 1}}, \\
&\to 1,
\end{aligned} \tag{S61}$$

*when $r_{\mathbf{a}} = \frac{\mathbf{a}^T\mathbf{a}}{m\sigma_{\mathbf{a}}^2} \neq 0$ and $r_{\mathbf{b}} = \frac{\mathbf{b}^T\mathbf{b}}{n\sigma_{\mathbf{b}}^2} \neq 0$.*

*Proof.* Let $\mathbf{A} = \frac{1}{r}\sum_{i=1}^r \mathbf{a}_i$, and $\mathbf{B} = \frac{1}{r}\sum_{i=1}^r \mathbf{b}_i$.

Then, we have:

$$\begin{aligned}
\mathbb{E}\left[\mathbf{a}_i^T\mathbf{A}\right] &= \mathbb{E}\left[\mathbf{A}^T\mathbf{A}\right] = \mathbf{a}^T\mathbf{a} + \frac{m\sigma_{\mathbf{a}}^2}{r}, \\
\mathbb{E}\left[\mathbf{b}_i^T\mathbf{B}\right] &= \mathbb{E}\left[\mathbf{B}^T\mathbf{B}\right] = \mathbf{b}^T\mathbf{b} + \frac{n\sigma_{\mathbf{b}}^2}{r} \\
\mathbb{E}\left[\sum_{i,j} \mathbf{a}_i^T\mathbf{a}_j\mathbf{b}_i^T\mathbf{b}_j\right] &= (r^2 - r)(\mathbf{a}^T\mathbf{a})(\mathbf{b}^T\mathbf{b}) + r(\mathbf{a}^T\mathbf{a} + m\sigma_{\mathbf{a}}^2)(\mathbf{b}^T\mathbf{b} + n\sigma_{\mathbf{b}}^2).
\end{aligned}$$

The cosine similarity between two vectors $\mathbf{u}$ and $\mathbf{v}$ is defined as:

$$\cos(\mathbf{u}, \mathbf{v}) = \frac{\mathbf{u}^T\mathbf{v}}{|\mathbf{u}||\mathbf{v}|},$$

where $|\cdot|$ denotes the L2 norm.

Then, consider the cosine similarity between $\frac{1}{r}\sum_{i=1}^r \mathbf{a}_i \otimes \mathbf{b}_i$ and $\mathbf{A} \otimes \mathbf{B}$:

$$\begin{aligned}
\cos\left(\frac{1}{r}\sum_{i=1}^r \mathbf{a}_i \otimes \mathbf{b}_i, \mathbf{A} \otimes \mathbf{B}\right) &= \frac{\frac{1}{r}\sum_i^r (\mathbf{a}_i^T\mathbf{A})(\mathbf{b}_i^T\mathbf{B})}{|\frac{1}{r}\sum_i^r \mathbf{a}_i \otimes \mathbf{b}_i||\mathbf{A} \otimes \mathbf{B}|}, \\
&= \frac{\sum_i^r (\mathbf{a}_i^T\mathbf{A})(\mathbf{b}_i^T\mathbf{B})}{\sqrt{\left(\sum_{i,j} (\mathbf{a}_i^T\mathbf{a}_j)(\mathbf{b}_i^T\mathbf{b}_j)\right)(\mathbf{A}^T\mathbf{A})(\mathbf{B}^T\mathbf{B})}}, \\
&\approx \frac{r(\mathbf{a}^T\mathbf{a} + \frac{m\sigma_{\mathbf{a}}^2}{r})(\mathbf{b}^T\mathbf{b} + \frac{n\sigma_{\mathbf{b}}^2}{r})}{\sqrt{r^2(\mathbf{a}^T\mathbf{a})(\mathbf{b}^T\mathbf{b}) + r(n\sigma_{\mathbf{b}}^2\mathbf{a}^T\mathbf{a} + m\sigma_{\mathbf{a}}^2\mathbf{b}^T\mathbf{b} + mn\sigma_{\mathbf{a}}^2\sigma_{\mathbf{b}}^2)}}, \\
&= \sqrt{\frac{(\mathbf{a}^T\mathbf{a} + \frac{m\sigma_{\mathbf{a}}^2}{r})(\mathbf{b}^T\mathbf{b} + \frac{n\sigma_{\mathbf{b}}^2}{r})}{(\mathbf{a}^T\mathbf{a})(\mathbf{b}^T\mathbf{b}) + \frac{1}{r}(n\sigma_{\mathbf{b}}^2\mathbf{a}^T\mathbf{a} + m\sigma_{\mathbf{a}}^2\mathbf{b}^T\mathbf{b} + mn\sigma_{\mathbf{a}}^2\sigma_{\mathbf{b}}^2)}}, \\
&= \sqrt{1 + \frac{(\frac{1}{r^2} - \frac{1}{r})mn\sigma_{\mathbf{a}}^2\sigma_{\mathbf{b}}^2}{(\mathbf{a}^T\mathbf{a})(\mathbf{b}^T\mathbf{b}) + \frac{1}{r}(n\sigma_{\mathbf{b}}^2\mathbf{a}^T\mathbf{a} + m\sigma_{\mathbf{a}}^2\mathbf{b}^T\mathbf{b} + mn\sigma_{\mathbf{a}}^2\sigma_{\mathbf{b}}^2)}}, \\
&= \sqrt{1 + \frac{(\frac{1}{r^2} - \frac{1}{r})}{r_{\mathbf{a}}r_{\mathbf{b}} + \frac{1}{r}(r_{\mathbf{a}} + r_{\mathbf{b}} + 1)}}.
\end{aligned} \tag{S62}$$

When $r \to \infty$, $\frac{1}{r^2} \to 0$, and Eq. S62 can be approximated by:

$$\cos\left(\frac{1}{r}\sum_{i=1}^{r}\mathbf{a}_i \otimes \mathbf{b}_i, \mathbf{A} \otimes \mathbf{B}\right) \approx \sqrt{1 - \frac{1}{r r_{\mathbf{a}} r_{\mathbf{b}} + r_{\mathbf{a}} + r_{\mathbf{b}} + 1}}. \tag{S63}$$

When $r_{\mathbf{a}} \neq 0$ and $r_{\mathbf{b}} \neq 0$,

$$\lim_{r\to\infty} \cos\left(\frac{1}{r}\sum_{i=1}^{r}\mathbf{a}_i \otimes \mathbf{b}_i, \mathbf{A} \otimes \mathbf{B}\right) = 1. \tag{S64}$$

□

Given the above Theorem E.1, now let us give a formal proof of Theorem C.3.

*Proof of Theorem C.3.* Since $\mathbf{a}_i \in \mathbb{R}^m$ and $\mathbf{b}_i \in \mathbb{R}^n$ be vectors where all elements in $\mathbf{a}_i$ and $\mathbf{b}_i$ are of the same sign (i.e., they are all positive or negative), we have:

$$\mathbb{E}[\mathbf{a}_i] \neq 0,$$
$$\mathbb{E}[\mathbf{b}_i] \neq 0.$$

Then we have:

$$\frac{\mathbf{a}^T\mathbf{a}}{m\sigma_{\mathbf{a}}^2} \neq 0,$$
$$\frac{\mathbf{b}^T\mathbf{b}}{n\sigma_{\mathbf{b}}^2} \neq 0.$$

According to Theorem E.1, we have:

$$\lim_{r\to\infty} \cos\left(\sum_{i=1}^{r}\mathbf{a}_i \otimes \mathbf{b}_i, \left(\frac{1}{r}\sum_{i=1}^{r}\mathbf{a}_i\right) \otimes \sum_{i=1}^{r}\mathbf{b}_i\right) \to 1.$$

□

The cosine similarity is close to 1 means that the resulting matrices are pointing in almost the same direction. In the context of Jacobian matrices, this implies that the rate of change of one set of variables with respect to another set of variables is highly similar between the two matrices.

### F Experimental validations of Theorem C.3

To verify the theoretical analysis above, we conducted two types of numerical experiments.

#### F.1 Evaluations on the random dataset

First, we performed experiments using randomly generated datasets that mimic the conditions a real synaptic projection might process. Specifically, we assumed that presynaptic neurons generated spikes according to a binomial distribution with probability $p$, and the derivative of the activation function $f'$ followed a normal distribution $\mathcal{N}(\mu_f, \sigma_f^2)$. We fixed the $\mu_f = 1.0$, $m = 10$, and $n = 10$. Then, we systematically varied $\sigma_f$ and $p$, generated corresponding $r$ random samples, and computed the cosine similarity between $\sum_{i=1}^{r} \mathbf{x}_i \otimes \mathbf{D}_f^i$ and $\left(\frac{1}{r}\sum_{i=1}^{r} \mathbf{x}_i\right) \otimes \sum_{i=1}^{r} \mathbf{D}_f^i$ (see Theorem C.3) using the generated samples. For each parameter pair, we used $r = 10,000$. The resulting phase portrait is visualized in Fig. S1A. We also computed the theoretical cosine similarity using Eq. S63. These theoretical results were plotted in Fig. S1B. As we can see, the theoretical and experimental results matched very well.

#### F.2 Evaluations on the neuromorphic dataset

Next, we calculated the cosine similarity between $\sum_{i=1}^{r} \mathbf{x}_i \otimes \mathbf{D}_f^i$ and $\left(\frac{1}{r}\sum_{i=1}^{r} \mathbf{x}_i\right) \otimes \sum_{i=1}^{r} \mathbf{D}_f^i$ using diverse SNN dynamics to process the IBM DVS Gestures dataset [36]. We examined how the cosine similarity changed as the state evolved forward over time. For each experiment, we computed 500 data samples and reported the mean and 95% confidence interval of cosine similarities across these samples. Experimental results are plotted in Fig. S1C-J. We observed that, over the entire sequence, the rank-one approximation of most eligibility traces maintained a very high cosine similarity (>80%). Moreover, there were two types of eligibility traces. In the first type, the cosine similarity gradually decayed over time. They correspond to the eligibility traces of fast decaying variables, such as the synaptic conductance $\mathbf{g}$ and the membrane potential $\mathbf{v}$ in models presented in SI. B (please also refer to the lines of "$\mathbf{x} \otimes f_{\mathbf{g}}$" and "$\mathbf{x} \otimes f_{\mathbf{v}}$" in Fig. S1C-J). In the second type, the cosine similarity started very low, then rapidly increased to approach 1.0 as time progressed, and remained high as the network states evolved. These eligibility traces track the neuronal and synaptic variables with slow times constants, such as the slow adaption current $\mathbf{a}$ and the short-term depression variable $\mathbf{x}$ in models presented in SI. B (please also refer to the lines of "$\mathbf{x} \otimes f_{\mathbf{x}}$" and "$\mathbf{x} \otimes f_{\mathbf{a}}$" in Fig. S1E-J). Overall, our empirical experiment demonstrated that the rank-one approximation can effectively capture the essential features of eligibility traces of most variables over long time sequences, and even achieve a perfect reproduction of eligibility traces for those variables with slow dynamics.

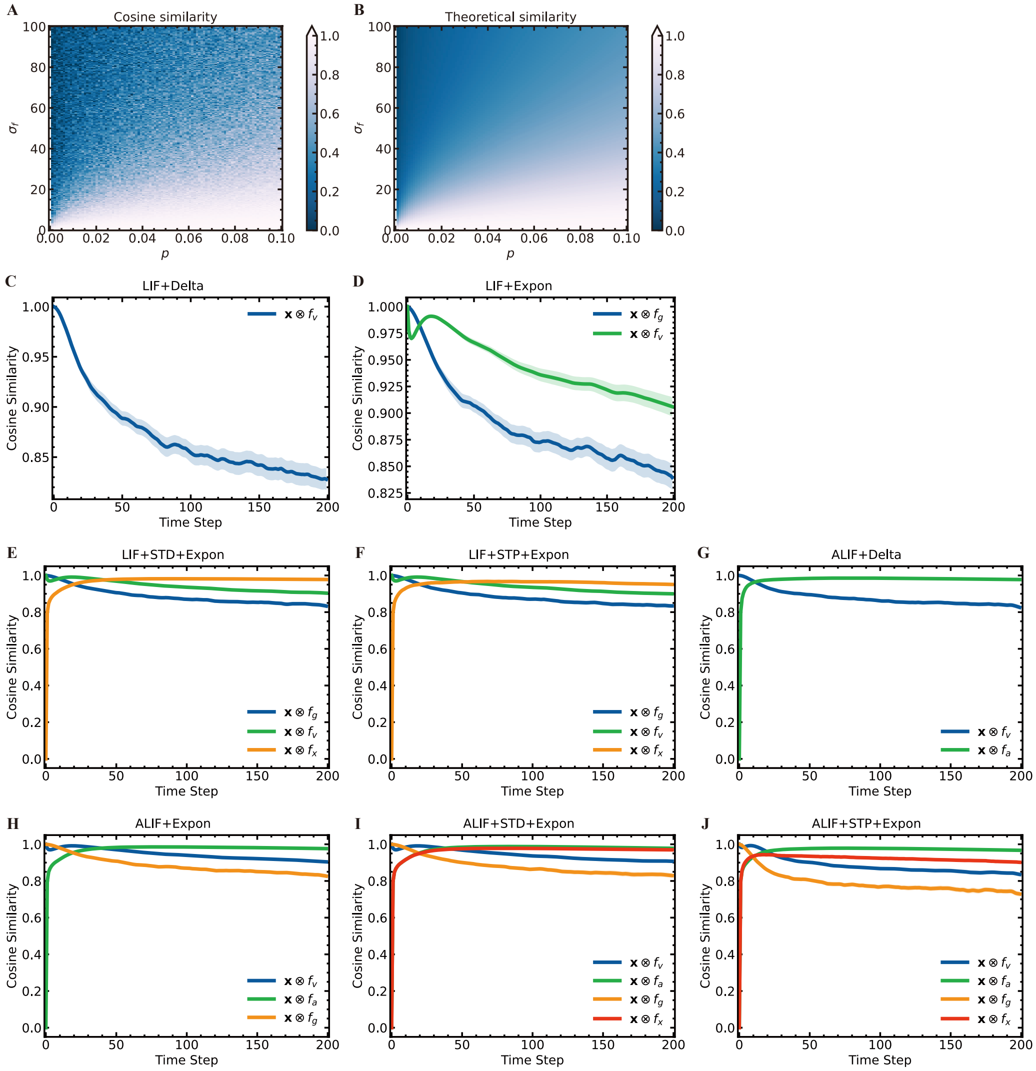

Figure S1: **Empirical Validation of Theorem E.1 and Theorem C.3**. (A) Phase portrait of cosine similarity as a function of presynaptic spiking probability $p$ and the standard deviation of postsynaptic activation derivatives $\sigma_f$. The standard deviation $\sigma_f$ varies widely from 1 to 100. The spiking probability $p$ is varied between 0 and 0.1. Note: Higher spiking probabilities are omitted due to consistently high cosine similarity. (B) Theoretical cosine similarity calculated using Eq. S63, using the same parameter range as in (A). (C-J) Empirical evaluations of cosine similarity across various SNN dynamics while processing the IBM DVS Gestures dataset [36]. The x-axis represents the time step, and the y-axis denotes the cosine similarity. Each panel (C-J) corresponds to a different SNN architecture: (C) `LIF+Delta` dynamics (Eq. S44); (D) `LIF+Expon` dynamics (Eq. S46); (E) `LIF+STD+Expon` dynamics (Eq. S49); (F) `LIF+STP+Expon` dynamics (Eq. S51); (G) `ALIF+Delta` dynamics (Eq. S45); (H) `ALIF+Expon` dynamics (Eq. S47); (I) `ALIF+STD+Expon` dynamics (Eq. S50); (J) `ALIF+STP+Expon` dynamics (Eq. S52). Each line within a panel represents the cosine similarity between exact and approximated eligibility traces for the corresponding hidden states. For example, $\mathbf{x} \otimes f_{\mathbf{v}}$ in (C) shows the cosine similarity of the eligibility trace tracked by hidden state $\mathbf{v}$ in the `LIF+Delta` network.

#### G Derivation of online learning algorithms

In this section, we provide the details of the algorithmic derivations underpinning our online learning algorithms.

##### G.1 Theoretical basics of RTRL

Let us consider biological SNN models whose dynamics are governed by Eq. 1. At each time step $t \in \mathcal{T}$, we assume that the state is transformed into an output $\mathbf{y}^t = f_o(\mathbf{h}^t, \boldsymbol{\phi})$, while the network receives a loss $\mathcal{L}^t(\mathbf{y}^t, \hat{\mathbf{y}}^t)$. The objective is to optimize the total loss $\mathcal{L} = \sum_t \mathcal{L}^t$ with respect to the parameters $\boldsymbol{\theta}$, utilizing the gradient

$$\nabla_{\boldsymbol{\theta}} \mathcal{L} = \sum_{t \in \mathcal{T}} \frac{\partial \mathcal{L}^t}{\partial \boldsymbol{\theta}} = \sum_{t \in \mathcal{T}} \sum_{k=1}^{t} \frac{\partial \mathcal{L}^t}{\partial \mathbf{h}^t} \frac{\partial \mathbf{h}^t}{\partial \mathbf{h}^k} \frac{\partial \mathbf{h}^k}{\partial \boldsymbol{\theta}^k}, \tag{S65}$$

where $\boldsymbol{\theta}^k$ represents the parameters utilized at time $t$, $\frac{\partial \mathcal{L}^t}{\partial \mathbf{h}^t} \frac{\partial \mathbf{h}^t}{\partial \mathbf{h}^k} \frac{\partial \mathbf{h}^k}{\partial \boldsymbol{\theta}^k}$ measures how $\boldsymbol{\theta}$ at step $k$ affects the loss at step $t \geq k$, and the factor $\frac{\partial \mathbf{h}^t}{\partial \mathbf{h}^k}$ transports the error through time from step $t$ to step $k$. When $k \ll t$, the transport captures a long-term dependency; otherwise, a short-term dependency.

To compute Eq. S65, two methods are commonly used, that is BPTT [37–39] and RTRL [40–42], which iteratively perform the error propagation on the unfolded computation graph through time backwardly and forwardly, respectively (also refer to Fig. S2).

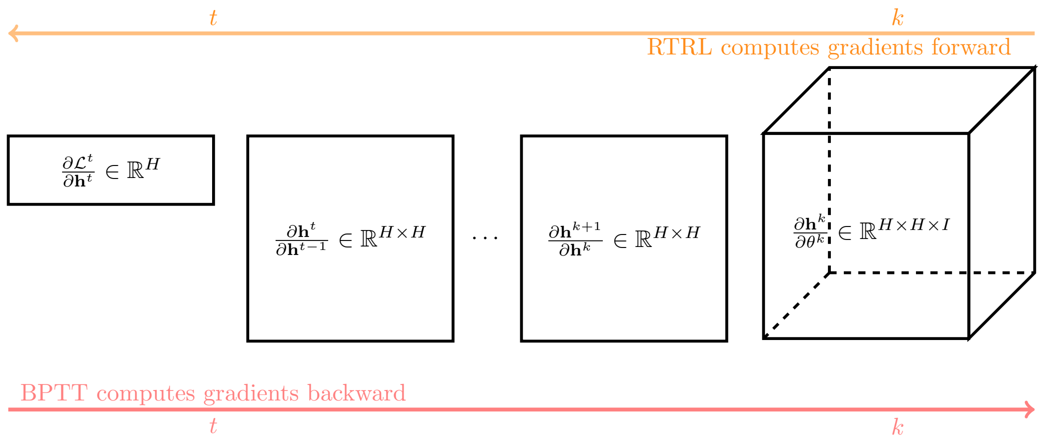

Figure S2: **Comparison of Gradient Computation Methods: BPTT vs. RTRL**. This figure illustrates the differences between BPTT and RTRL in computing gradients $\frac{\partial \mathcal{L}^t}{\partial \boldsymbol{\theta}^k}$ for $t \geq k$. **BPTT**: Gradients are computed in a backward manner. The hidden states $\mathbf{h}$ are stored from time steps $k$ to $t$. The gradient $\frac{\partial \mathcal{L}^t}{\partial \boldsymbol{\theta}^k}$ is then propagated backward through time using the chain rule $\frac{\partial \mathcal{L}^t}{\partial \boldsymbol{\theta}^k} = \frac{\partial \mathcal{L}^t}{\partial \mathbf{h}^t} \frac{\partial \mathbf{h}^t}{\partial \mathbf{h}^{t-1}} \cdots \frac{\partial \mathbf{h}^k}{\partial \boldsymbol{\theta}^k}$. **RTRL**: Gradients are computed in a forward manner. At time step $k$, the influence Jacobian $\boldsymbol{\epsilon}^k = \frac{\partial \mathbf{h}^k}{\partial \boldsymbol{\theta}^k}$ is calculated. At the next time step $k+1$, the influence Jacobian is updated to $\boldsymbol{\epsilon}^{k+1} = \frac{\partial \mathbf{h}^{k+1}}{\partial \mathbf{h}^k} \boldsymbol{\epsilon}^k$. This process continues forward in time, with $\boldsymbol{\epsilon}^{k+2} = \frac{\partial \mathbf{h}^{k+2}}{\partial \mathbf{h}^{k+1}} \boldsymbol{\epsilon}^{k+1}$, and so forth.

For the BPTT, the gradient is solved iteratively, spanning from the final time step to the first one:

$$\nabla_{\boldsymbol{\theta}} \mathcal{L} = \sum_{t \in \mathcal{T}} \frac{\partial \mathcal{L}}{\partial \mathbf{h}^t} \frac{\partial \mathbf{h}^t}{\partial \boldsymbol{\theta}^t} = \sum_{t \in \mathcal{T}} \Phi^t \frac{\partial \mathbf{h}^t}{\partial \boldsymbol{\theta}^t}, \tag{S66}$$

$$\Phi^t = \Phi^{t+1} \frac{\partial \mathbf{h}^{t+1}}{\partial \mathbf{h}^t} + \frac{\partial \mathcal{L}^t}{\partial \mathbf{h}^t}, \tag{S67}$$

where $\Phi \in \mathbb{R}^H$ defines the partial gradients of the loss with respect to the hidden state. Here, we only use the batch size $B = 1$.

For the RTRL, the gradient is updated in the forward direction:

$$\nabla_{\boldsymbol{\theta}}\mathcal{L} = \sum_{t\in\mathcal{T}} \frac{\partial \mathcal{L}^t}{\partial \mathbf{h}^t}\frac{\partial \mathbf{h}^t}{\partial \boldsymbol{\theta}} = \sum_{t\in\mathcal{T}} \frac{\partial \mathcal{L}^t}{\partial \mathbf{h}^t}\boldsymbol{\epsilon}^t, \tag{S68}$$

$$\boldsymbol{\epsilon}^t = \frac{\partial \mathbf{h}^t}{\partial \mathbf{h}^{t-1}}\boldsymbol{\epsilon}^{t-1} + \frac{\partial \mathbf{h}^t}{\partial \boldsymbol{\theta}^t}, \tag{S69}$$

where the influence matrix $\boldsymbol{\epsilon} \in \mathbb{R}^{H\times H\times I}$ calculates the effect of the hidden units on the parameters.

To better illustrate the difference between BPTT and RTRL, let's examine the gradients of the loss at time $t$ with respect to the parameter at time $k$, denoted as $\frac{\partial \mathcal{L}^t}{\partial \boldsymbol{\theta}^k}$, where $t \geq k$. Fig. S2 demonstrates that BPTT's backward computation of gradients is significantly more efficient than RTRL's forward computation. This is because the product of Jacobians from the end to the beginning of the composed function only requires $\mathcal{O}(H)$ memory storage. In contrast, RTRL's memory complexity is $\mathcal{O}(H \times H \times I)$ as it computes Jacobian products forwardly. To reduce the intrinsic complexity of RTRL, we should take full consideration of SNN properties.

##### G.2 Derivation of the D-RTRL algorithm from RTRL

In the following sections, we will concentrate on the scenario where the hidden state dimension $d = 1$ to develop the D-RTRL algorithm. This simplification enables us to clearly demonstrate the core principles of the algorithm without the additional complexity introduced by multi-dimensional hidden states. For learning rules applicable when ($d > 1$), please refer to SI. G.4.

As stated in the above sectons, the raw RTRL has a high memory complexity. By substituting Eq. S58 and Eq. S59 into Eq. S69, we can approximate $\boldsymbol{\epsilon}^t$ as:

$$\boldsymbol{\epsilon}^t \approx \underbrace{\mathbf{D}^t}_{\approx \partial \mathbf{h}^t/\partial \mathbf{h}^{t-1}} \boldsymbol{\epsilon}^{t-1} + \underbrace{\mathbf{D}^t_f \otimes \mathbf{x}^t}_{=\partial \mathbf{h}^t/\partial \boldsymbol{\theta}^t}. \tag{S70}$$

Eq. S70 results in a very sparse matrix with $\mathcal{O}(H \times H \times I)$ memory complexity. For compact storage, the above equation is equivalent to the following form:

$$\boldsymbol{\epsilon}^t \approx \mathbf{D}^t \boldsymbol{\epsilon}^{t-1} + \mathrm{diag}(\mathbf{D}^t_f) \otimes \mathbf{x}^t, \tag{S71}$$

where $\mathrm{diag}(\mathbf{D}^t_f) \in \mathbb{R}^H$ is the vector storing the diagonal elements of $\mathbf{D}^t_f$. This results in $\boldsymbol{\epsilon}^t$ having the shape of $\mathbb{R}^{H\times I}$.

In practice, $\mathbf{D}^t$ is stored as a vector $\mathrm{diag}(\mathbf{D}^t) \in \mathbb{R}^{H\times 1}$. Consequently, Eq. S71 can be reformulated using element-wise multiplication. Combining with Eq. S68, we can express the learning rule of the D-RTRL algorithm as:

$$\begin{aligned} \nabla_{\boldsymbol{\theta}}\mathcal{L} &\approx \sum_{t\in\mathcal{T}} \frac{\partial \mathcal{L}^t}{\partial \mathbf{h}^t} \circ \boldsymbol{\epsilon}^t, \\ \boldsymbol{\epsilon}^t &\approx \mathrm{diag}(\mathbf{D}^t) \circ \boldsymbol{\epsilon}^{t-1} + \mathrm{diag}(\mathbf{D}^t_f) \otimes \mathbf{x}^t. \end{aligned} \tag{S72}$$

where $\frac{\partial \mathcal{L}^t}{\partial \mathbf{h}^t} \in \mathbb{R}^{H\times 1}$ is a column vector.

D-RTRL generally exhibits a memory complexity of $\mathcal{O}(BP)$, where $B$ represents the batch size and $P$ denotes the number of parameters. This efficient scaling is primarily due to the nature of the eligibility trace $\boldsymbol{\epsilon}^t$, which only needs to store the historical information for each active synaptic connection. The memory efficiency of D-RTRL becomes particularly advantageous when dealing with sparse neural networks, a common characteristic in biologically inspired models. In sparse networks, many synaptic connections have zero weights. D-RTRL capitalizes on this sparsity by eliminating the need to store eligibility traces for these zero-weight connections. This optimization can lead to significant memory savings, especially in large-scale networks where sparsity is pronounced. In convolutional layers, D-RTRL exhibits greater memory efficiency since the parameter size is typically smaller than the number of presynaptic and postsynaptic neurons for small-sized convolutional kernels.

Moreover, the $\mathcal{O}(BP)$ complexity of D-RTRL represents a substantial improvement over the $\mathcal{O}(BH^2)$ complexity of previous online algorithms such as E-prop [33] and OSTL [43], where $H$ is the number of hidden units. This reduction in complexity enables D-RTRL to scale more effectively to larger networks and longer temporal sequences, making it a promising approach for training complex spiking networks.

##### G.3 Derivation of the ES-D-RTRL algorithm from D-RTRL

In this section, we derive the learning rule for the ES-D-RTRL algorithm by focusing on the scenario where the hidden state dimension $d = 1$. The derived learning rule can naturally extend to cases where $d > 1$. For more details on the generalization to multi-dimensional hidden states, please refer to SI. G.4.

Given the learning rule of the D-RTRL algorithm in Eq. S72, let's consider the influence of $\boldsymbol{\theta}$ at step $k$ to the loss $\mathcal{L}$ at step $t$ ($t \geq k$). It can be approximated by:

$$\frac{\partial \mathcal{L}^t}{\partial \boldsymbol{\theta}^k} \approx \frac{\partial \mathcal{L}^t}{\partial \mathbf{h}^t} \underbrace{\left( \prod_{i=k+1}^{t} \mathbf{D}^i \right)}_{\approx \partial \mathbf{h}^t / \partial \mathbf{h}^k} \underbrace{\left( \mathbf{D}_f^k \otimes \mathbf{x}^k \right)}_{= \partial \mathbf{h}^k / \partial \boldsymbol{\theta}^k} . \tag{S73}$$

Therefore, at the time $t$, the parameter gradient accumulated up to time $t$ has the form of:

$$\nabla_{\boldsymbol{\theta}} \mathcal{L}^t = \sum_{k=1}^{t} \frac{\partial \mathcal{L}^t}{\partial \boldsymbol{\theta}^k} \approx \frac{\partial \mathcal{L}^t}{\partial \mathbf{h}^t} \sum_{k=1}^{t} \left( \prod_{i=k+1}^{t} \mathbf{D}^i \right) \left( \mathbf{D}_f^k \otimes \mathbf{x}^k \right) . \tag{S74}$$

Here, we assume that a learning target is provided at each time step $k \in [1, \cdots, t]$. Moreover, $\frac{\partial \mathcal{L}^t}{\partial \mathbf{h}^t} \in \mathbb{R}^H$ is defined as a row vector in Supplementary Eqs. S73 and S74.

Since $\mathbf{D}^i$ and $\mathbf{D}_f$ are diagonal matrices, we can only keep their diagonal elements as vectors, and their matrix multiplication operations are equivalent to the element-wise Hadamard products of diagonal vectors:

$$\nabla_{\boldsymbol{\theta}} \mathcal{L}^t \approx \frac{\partial \mathcal{L}^t}{\partial \mathbf{h}^t} \circ \left[ \sum_{k=1}^{t} \left( \underbrace{\mathrm{diag}(\mathbf{D}^t) \circ \cdots \circ \mathrm{diag}(\mathbf{D}^{k+1}) \circ \mathrm{diag}(\mathbf{D}_f^k)}_{\text{postsynaptic activity}} \right) \otimes \underbrace{\mathbf{x}^k}_{\text{presynaptic activity}} \right] . \tag{S75}$$

From this equation onward, $\frac{\partial \mathcal{L}^t}{\partial \mathbf{h}^t} \in \mathbb{R}^{H \times 1}$ is defined as a column vector.

Eq. S75 results in significant memory overhead because the summation requires storing t pairs of vectors representing pre- and postsynaptic activity. This memory requirement is equivalent to BPTT, with complexity scaling linearly as time progresses.

We found a solution to this issue by fully leveraging the unique properties of SNNs. The key insight lies in the nature of neuronal output of SNNs: (1) Binary events: neurons in SNNs output binary signals (spikes); (2) Non-negativity: these spike events are always non-negative. These properties are detailed in SI. C, and the foundation of our solution is provided by Theorem C.3. This theorem allows us to approximate Eq. S75 by

$$\nabla_{\boldsymbol{\theta}} \mathcal{L}^t \approx \frac{\partial \mathcal{L}^t}{\partial \mathbf{h}^t} \circ \left( \frac{1}{t} \sum_{k=1}^{t} \mathbf{b}_k^t \otimes \sum_{k=1}^{t} \mathbf{x}^k \right) , \tag{S76}$$

where $\mathbf{b}_k^t = \mathrm{diag}(\mathbf{D}^t) \circ \cdots \circ \mathrm{diag}(\mathbf{D}^{k+1}) \circ \mathrm{diag}(\mathbf{D}_f^k)$.

To avoid the exploding problem, the eligibility trace should prioritize recent activities rather than retain a complete historical record over a very long sequence. We implemented this by modifying Eq. S76 using two key techniques: exponential smoothing for $\mathbf{b}^t$ and a low-pass filter for $\mathbf{x}$. The former computes a running average of $\mathbf{b}^t$ online, while the latter calculates a running summation of $\mathbf{x}$. These adaptations form the foundation of our ES-D-RTRL algorithm, whose learning rule can be expressed as follows:

$$\begin{aligned} \nabla_{\boldsymbol{\theta}} \mathcal{L} &\approx \sum_{t=\mathcal{T}} \frac{\partial \mathcal{L}^t}{\partial \mathbf{h}^t} \circ \left( \boldsymbol{\epsilon}_{\mathbf{f}}^t \otimes \boldsymbol{\epsilon}_{\mathbf{x}}^t \right) , \\ \boldsymbol{\epsilon}_{\mathbf{x}}^t &= \alpha \boldsymbol{\epsilon}_{\mathbf{x}}^{t-1} + \mathbf{x}^t, \\ \boldsymbol{\epsilon}_{\mathbf{f}}^t &= \alpha\, \mathrm{diag}(\mathbf{D}^t) \circ \boldsymbol{\epsilon}_{\mathbf{f}}^{t-1} + (1-\alpha)\mathrm{diag}(\mathbf{D}_f^t), \end{aligned} \tag{S77}$$

where $\alpha$ $(0 < \alpha < 1)$ is the smoothing factor, corresponding to the time constant $\tau = -\Delta t / \ln(\alpha)$. The only

##### G.4 Generalization to hidden states with multiple variables ($d > 1$)

It is important to note that the derivations presented in SI. G.2 and G.3 are based on the simplifying assumption that $d = 1$, where $d$ represents the number of variables in each neuron's hidden states. However, these derivations can be generalized to cases where $d > 1$ without loss of generality.

When $d > 1$, the following adjustments to the matrix dimensions and structures are necessary:

- $\mathbf{h}^t$: The hidden state becomes a block matrix with dimensions $\mathbb{R}^{Hd\times 1}$, where $H$ is the number of hidden units. It contains $H$ blocks with each block is a matrix of shape $\mathbb{R}^{d\times 1}$. The block $\mathbf{h}_i$ belongs to the state variables of neuron $i$ (see examples in SI. B).
- $\mathbf{D}^t$: This matrix becomes a block diagonal matrix with dimensions $\mathbb{R}^{Hd\times Hd}$. It consists of $H$ blocks along the diagonal, each block being a $d \times d$ matrix. Each diagonal block $\mathbf{D}^t_{ii}$ belongs to the Jacobian matrix between $d$ state variables of neuron $i$ (see examples in SI. B).
- $\mathbf{D}^t_f$: This matrix transforms into a block diagonal matrix of shape $\mathbb{R}^{Hd\times H}$. It comprises $H$ blocks along the diagonal, where each block is a matrix of shape $\mathbb{R}^{d\times 1}$. Each diagonal block $\mathbf{D}_{f,ii} \in \mathbb{R}^{b\times 1}$ belongs to the hidden-to-weight Jacobian $\partial \mathbf{h}^t_i / \partial \mathbf{I}^t_i$ of neuron $i$ (see examples in SI. B).
- $\frac{\partial \mathcal{L}^t}{\partial \mathbf{h}^t}$: The gradient of the loss with respect to the hidden state becomes a block matrix of shape $\mathbb{R}^{Hd\times 1}$. It consists of $H$ blocks, each with dimensions $\mathbb{R}^{d\times 1}$.

As we can see, in the case of multi-dimensional hidden states ($d > 1$), the matrices in Supplementary Eqs. S72 and S77 transform into block matrices. Each element of these block matrices is no longer a scalar, but rather a sub-matrix or vector. However, the transition from scalar ($d = 1$) to multi-dimensional ($d > 1$) hidden states maintains the overall online learning framework as shown in Supplementary Eqs. S72 and S77. The only difference is on the $\circ$ operations in the aforementioned equations.

For $d = 1$, $\circ$ represents the ***element-wise scalar multiplication***. This is because $\mathbf{h}^t_i$, $\mathbf{D}^t_{ii}$, $\mathbf{D}^t_{f,ii}$, and $\frac{\partial \mathcal{L}^t}{\partial \mathbf{h}^t_i}$ are all scalar values.

For $d > 1$, $\circ$ evolves into a ***block-wise multiplication*** operation. In this context, the multiplication is performed on corresponding blocks, with each block undergoing $d$-dimensional matrix or inner multiplication.

To elucidate this distinction, let us examine the learning rule for a single weight $\boldsymbol{\theta}_{ji}$ connecting neurons $i$ and $j$. For the D-RTRL algorithm, the block-wise implementation of Eq. S72 becomes:

$$\begin{aligned} \nabla_{\boldsymbol{\theta}_{ji}} \mathcal{L} &\approx \sum_{t \in \mathcal{T}} \left\langle \frac{\partial \mathcal{L}^t}{\partial \mathbf{h}^t_j} \,,\, \boldsymbol{\epsilon}^t_{ji} \right\rangle, \\ \boldsymbol{\epsilon}^t_{ji} &\approx \mathbf{D}^t_{jj} \boldsymbol{\epsilon}^{t-1}_{ji} + \mathbf{D}^t_{f,jj} \cdot {\mathbf{x}_i}^t, \end{aligned} \tag{S78}$$

where ${\mathbf{x}_i}^t \in \mathbb{R}$ is a scalar value, $\boldsymbol{\epsilon}^t \in \mathbb{R}^{Hd\times H}$ now becomes a block matrix with $H$-by-$H$ blocks and each block $\boldsymbol{\epsilon}^t_{ji}$ is a vector with the shape of $\mathbb{R}^{d\times 1}$, and $\langle \cdot \,,\, \cdot \rangle : \mathbb{R}^{d\times 1} \times \mathbb{R}^{d\times 1} \to \mathbb{R}$ is the inner product between two vectors.

Similarly, for the ES-D-RTRL algorithm, the learning rule in Eq. S77 is implemented block-wise as:

$$\begin{aligned} \nabla_{\boldsymbol{\theta}_{ji}} \mathcal{L} &\approx \sum_{t = \mathcal{T}} \left\langle \frac{\partial \mathcal{L}^t}{\partial \mathbf{h}^t_j} \,,\, \boldsymbol{\epsilon}^t_{\mathbf{f},j} \right\rangle \cdot \boldsymbol{\epsilon}^t_{\mathbf{x},i}, \\ \boldsymbol{\epsilon}^t_{\mathbf{f},j} &= \alpha \mathbf{D}^t_{jj} \boldsymbol{\epsilon}^{t-1}_{\mathbf{f},j} + (1-\alpha) \mathbf{D}^t_{f,jj}, \end{aligned} \tag{S79}$$

where $\boldsymbol{\epsilon}^t_{\mathbf{f}} \in \mathbb{R}^{Hd\times 1}$ now becomes a block matrix with each block $\boldsymbol{\epsilon}^t_{\mathbf{f},j}$ a vector with the shape of $\mathbb{R}^{d\times 1}$, $\boldsymbol{\epsilon}^t_{\mathbf{x},i} \in \mathbb{R}$ is a scalar value that tracking the eligibility trace of presynaptic neuron $i$.

These formulations highlight the critical differences in computational operations between scalar and multi-dimensional hidden states. The transition from element-wise to block-wise operations not only affects the mathematical representation but also has implications for computational efficiency and implementation strategies in practice. Furthermore, this generalization to multi-dimensional hidden states opens up new possibilities for capturing more complex temporal dependencies and representations in SNNs, potentially leading to enhanced performance in various sequence modeling tasks.

##### G.5 Learning signals in deep recurrent layers

In a network with $L$ stacked recurrent layers, where $\mathbf{h}^t = [\mathbf{h}_1^t, \cdots, \mathbf{h}_L^t]$ represents the hidden states at time $t$, the computation of exact eligibility traces is inherently very complex [42]. This complexity arises from the interdependencies between layers and the propagation of gradients through time. To maintain the simplicity and efficiency of the algorithms discussed in Section , we introduce an approximation that simplifies these calculations.

Our approach involves ignoring the influence of hidden states in deeper layers $(\mathbf{h}_{l+1}, \cdots, \mathbf{h}_L)$ on the parameters of the $l$-th layer, denoted as $\boldsymbol{\theta}_l$. Instead, we focus on maintaining the eligibility trace $\boldsymbol{\epsilon}_l^t$ with respect to its own hidden state $\mathbf{h}_l$ (Fig. 2D). This simplification allows for more tractable computations while still capturing the essential dynamics of the recurrent network.

When a top-down learning signal reaches layer $l$ at time $t \in \mathcal{T}$, we can compute its parameter gradients using the following equation:

$$\nabla_{\boldsymbol{\theta}_l} \mathcal{L} = \sum_{t \in \mathcal{T}} \frac{\partial \mathcal{L}^t}{\partial \mathbf{h}_l^t} \circ \boldsymbol{\epsilon}_l^t. \tag{S80}$$

##### G.6 Theoretical analysis of memory efficiency and computational speed

The summary of memory and time complexity of the evaluated algorithms is presented in Fig. 4I.

BPTT has a memory complexity of $\mathcal{O}(TBN)$, where $T$ is the number of time steps unrolled during training, $B$ is the batch size, and $N$ is the hidden state size. Generally, the memory complexity of RTRL is cubic in the number of hidden units at a single time step, making it impractical for large networks. However, when $T > N^2$, RTRL shows a memory advantage over BPTT. E-prop [33], an approximation of RTRL, provides a biologically plausible learning rule for spiking networks but has a memory complexity of $\mathcal{O}(BN^2)$ for maintaining the eligibility trace. OSTL [43] shares the same memory complexity as E-prop. Both E-prop and OSTL are designed for densely connected recurrent layers. In contrast, our D-RTRL has a memory complexity proportional to the parameter size $P$, achieving $\mathcal{O}(BP)$ complexity. For densely connected layers, D-RTRL matches the memory complexity of E-prop and OSTL at $\mathcal{O}(BN^2)$. For sparsely connected layers, D-RTRL is more memory-efficient than E-prop and OSTL, as it only stores the eligibility trace at non-zero weight positions. In convolutional layers, D-RTRL exhibits greater memory efficiency since the parameter size is typically smaller than the number of presynaptic and postsynaptic neurons for small-sized convolutional kernels. ES-D-RTRL shows a memory complexity of $\mathcal{O}(BN)$, scaling linearly with the number of neurons in the network because it only needs to store two traces of presynaptic and postsynaptic activities (refer to Eq. 7-8). For densely connected layers, ES-D-RTRL is more memory-efficient than D-RTRL, as the number of pre-synaptic and post-synaptic neurons is usually smaller than the number of connections between them. However, for convolutional layers with small kernels, D-RTRL may be more memory-efficient than ES-D-RTRL.

The computational complexity of BPTT is relatively low because the backward gradient computation at each time step involves a batched VJP with a complexity of $\mathcal{O}(BN^2)$. Raw RTRL has a time complexity of $\mathcal{O}(BN^4)$ per step, making it nearly impossible to use for training networks with $N > 1000$. Both OSTL and E-prop have a time complexity of $\mathcal{O}(BN^2)$. The computational complexity of D-RTRL depends on the size of $\boldsymbol{\theta}$. For densely connected layers, it is $\mathcal{O}(BN^2)$; for sparsely connected layers, it is $\mathcal{O}(pBN^2)$, where $p$ is the connection probability; for convolutional layers, the time complexity depends on the size of the convolutional kernels. It is important to note that while BPTT and D-RTRL both have a theoretical complexity of $\mathcal{O}(BN^2)$ for dense connection layers, they differ in practice. In BPTT, it is a matrix-matrix multiplication, a computation-bound operation, while in D-RTRL, it is a batched matrix-vector element-wise multiplication (refer to Fig. 2B and Eq. 4), a memory-bound operation. The ES-D-RTRL algorithm has a computational complexity of $\mathcal{O}(BN)$ for eligibility trace omputations, which are element-wise operations among presynaptic and postsynaptic neuronal activities (refer to Fig. 2C and Eq. 8).

#### H Approximation accuracy on weight gradients

We investigated the factors that affect the performance of the hidden Jacobian and weight gradient estimations in our online learning algorithms.

One prominent factor might be the network size, which determines the scalability of our proposed learning algorithms for large-scale SNN networks. Our evaluations demonstrated that the size of recurrent units did not significantly influence the Jacobian and gradient approximation, regardless of the evaluated dataset or network dynamics (Fig. S9). This result indicates that our online learning rules can be effectively applied to larger SNN models. Remarkably, for the ES-D-RTRL algorithm, we even observed a slight increase in gradient cosine similarity as the network size increased.

Another factor we explored was the firing rate, which is determined by the feedforward and recurrent connections. By increasing the feedforward connection strength, we observed an increase in firing rate accompanied by a slight decrease in cosine similarity for both Jacobian and gradient estimation (Fig. S10). However, for the ES-D-RTRL algorithm, the gradient cosine similarity consistently increased, indicating a preference for higher firing rates. Notably, the strength of recurrent connections had a more pronounced effect than feedforward connections. Increasing the strength of recurrent connections resulted in substantially lower cosine similarity (Fig. S11). This effect can be attributed to the increase in both firing frequency and the size of non-zero elements in the non-zero columns of the Jacobian matrix caused by the recurrent spiking connections.

Finally, we examined the depth of recurrent layers on weight gradients. We assessed the layer-wise cosine similarity between online gradient estimates and offline BPTT gradients using the SHD dataset [44]. Specifically, we evaluated a five-layer SNN network with 200 hidden units per layer, considering a variety of neuronal and synaptic dynamics. Consistently, we observed a decrease in cosine similarity as the layer depth increased (see Fig. S12). Lower layers exhibited lower cosine similarity to their exact gradients, corresponding to the fact that more recurrence information of hidden states in top layers is ignored.

### I Experimental details for cognitive tasks

This section provides an overview of the experimental methodology employed in our study for training spiking networks on cognitive tasks.

#### I.1 Surrogate gradient function

Translating membrane potential fluctuations into discrete spike events, as described by Eq. S10, relies on a Heaviside step function with its transition point at the neuron's firing threshold $v_{\text{th}}$. However, this function presents challenges for gradient-based learning algorithms due to its discontinuity at the threshold and lack of meaningful gradients elsewhere. To overcome these limitations and enable effective training of spiking networks, we employed a surrogate gradient approach used in previous works [45].

During the forward pass, we use the Heaviside function to generate the spike:

$$\mathbf{z}^t = \text{spike}(x) = \mathcal{H}(\mathbf{v}^t - v_{\text{th}}),$$

where $x$ is used to represent $\mathbf{v}^t - v_{\text{th}}$.

During the backward pass, we approximated the derivative of the Heaviside step function with the ReLU function:

$$\text{spike}'(x) = \text{ReLU}(\alpha * (\text{width} - |x|))$$

where $\mathrm{width} = 1.0$, and $\alpha = 0.3$. $\alpha$ is the parameter that controls the altitude of the gradient, and $\mathrm{width}$ is the parameter that controls the width of the gradient. The derivative of the spike function, $\text{spike}'(x)$, is a triangular function centered at $x = 0$. It reaches its maximum value of $\alpha * \mathrm{width}$ at $x = 0$ and decreases linearly to zero at $x = \pm\text{width}$. When $|x| > \text{width}$, $\text{spike}'(x) = 0$.

#### I.2 Weight initialization

For the networks described in Supplementary Eqs. S44-S52, we employed a specific weight initialization strategy for input, recurrent, and readout connections. The weights were sampled from a scaled Gaussian distribution, defined as: $W_{ji} \sim \sqrt{\frac{s}{N_{\text{in}}}}\mathcal{N}(0,1)$, where $W_{ji}$ represents the weight from neuron $i$ to neuron $j$, $N_{\text{in}}$ denotes the number of afferent (input-providing) neurons, $\mathcal{N}(0,1)$ is the standard normal distribution with mean 0 and variance 1, $s$ is a scaling factor that allows for control over the initial magnitude of the weights. This initialization schema helps maintain consistent variance of activations and gradients across layers, potentially aiding in training stability and convergence.

For the network with E/I separation, as described in Eq. S55, we modified the initialization procedure slightly. While still using the scaled Gaussian distribution, we applied an absolute value operation to ensure all weights were non-negative: $W_{ji} \sim \sqrt{\frac{s}{N_{\text{in}}}}|\mathcal{N}(0,1)|$. This modification ensures that excitatory remain excitatory and inhibitory connections remain inhibitory, adhering to Dale's principle often observed in biological networks. The absolute value operation maintains the scale and distribution shape while restricting weights to positive values.

#### I.3 Delayed match-to-sample task

We applied our learning algorithms to train a recurrent spiking network (Eq. S47) on a delayed match-to-sample (DMTS) task [46, 47] to test the long-term dependency learning capability of our learning algorithms (see Algorithmic evaluations on approximation, long-term dependency, memory, and speed). This task comprises four phases: fixation, sample, delay, and test (Fig. S3). The network includes 100 motion neurons, each covering a 360-degree range of directions. During the fixation phase, the network dynamics are initialized, and motion neurons fire at the baseline rate due to the absence of stimuli. In the sample and test phases, one of eight motion directions is presented to the motion neurons and lasts 500 ms. The delay phase, lasting over 1000 ms, occurs between the sample and test phases, during which motion neurons revert to the baseline firing rate. This task is challenging since it requires the network to maintain the sample direction information over a long delay period and then make a decision based on the test stimulus. The decision is made during the test

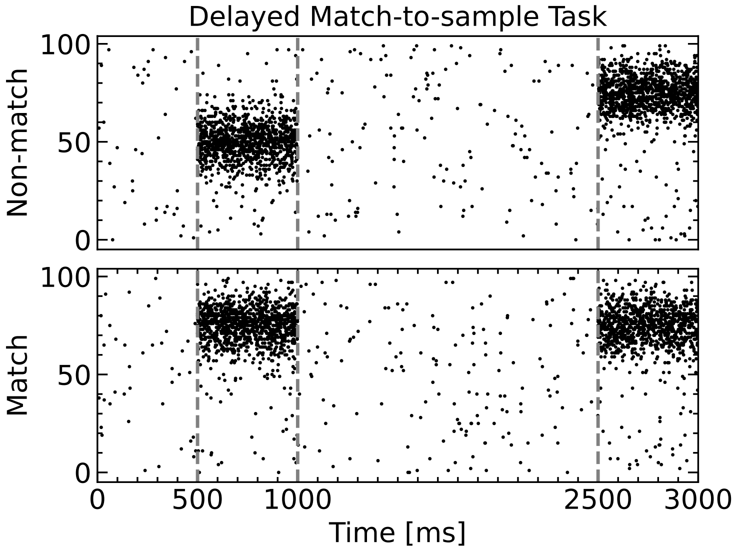

Figure S3: **Spiking Dynamics of Input Neurons During the DMTS Task: Two Examples.** The top panel illustrates the non-match case, while the bottom panel shows the match case. The task consists of four sequential phases: fixation (500 ms), sample (500 ms), delay (1000 ms), and test (500 ms).

phase, where a leaky readout layer processes network activity to determine whether the sequentially presented directions match or not.

In the DMTS task, the input to the network is provided by 100 motion-direction-tuned neurons. These neurons are designed to simulate the response properties of direction-selective cells found in visual cortical areas such as the middle temporal area. The tuning of these motion direction-selective neurons follows a von Mises distribution, which is a circular analog of the Gaussian distribution appropriate for directional data. The firing rate activity $u_i^t$ of input neuron $i$ during the stimulus presentation period is given by:

$$u_i^t = A \exp\left(\kappa \cos\left(\theta - \theta_{\mathrm{pref}}^i\right)\right) \tag{S81}$$

where $\theta$ is the direction of the presented stimulus, $\theta_{\mathrm{pref}}^i$ is the preferred direction of input neuron $i$, $\kappa$ is the concentration parameter, which is set to 2.0 in this study, and $A$ is the maximum firing rate, which is set to 40 Hz.

The von Mises distribution provides a peak response when the stimulus direction $\theta$ matches the neuron's preferred direction $\theta_{\mathrm{pref}}^i$, with a smooth fall-off for other directions. The concentration parameter $\kappa$ controls the width of this tuning curve, with higher values resulting in sharper tuning.

In addition to their direction-tuned responses during stimulus presentation, these input neurons maintain a constant background firing rate $r_{\mathrm{bg}}$ during all periods of the task, including inter-stimulus intervals. This background rate is set to 1 Hz, simulating the spontaneous activity observed in cortical neurons even in the absence of specific stimuli.

Finally, we generated the spike train of motion direction-tuned neurons using a Poisson process with the above-calculated firing rate $u_i^t + r_{\mathrm{bg}}$ during sample and test periods. Two examples were shown in Fig. S3.

##### I.4 Evidence accumulation task

The evidence accumulation paradigm is a fundamental tool in neuroscience and cognitive psychology for exploring the mechanisms of decision-making. Our study utilizes a task design similar to the one described by Morcos et al. [48], where a virtual-reality T-maze assesses the decision-making capabilities of a head-restrained mouse. As the mouse navigates the maze, it encounters visual cues on both the left and right sides, as illustrated in Fig. 5A. Upon reaching the T-junction, the mouse must choose a direction based on the majority of cues received from one side, regardless of the sequence or the position of the last cue. The complexity of the task lies in the requirement for the mouse to count and compare cues from each side independently and retain this information until the decision point, well before any reward is offered.

To simulate this task, we employed a population consisting of 100 neurons segmented into four groups. The first and second groups, each comprising 25 neurons, encode the cues from the left and right sides, respectively. Another 25 neurons process the recall cue during a subsequent recall period, and the final group of 25 neurons generates constant background noise at a rate of 10 Hz. The cues and the recall stimuli are modeled with Poisson noise at a firing rate of 40 Hz, with each stimulus lasting 150 ms and interspersed with 50 ms intervals of silence. The recall stimulus follows a delay of 1000 ms, during which background noise persists. The operational details of this neural representation are further depicted in Fig. 5A.

##### I.5 The leaky rate readout

The networks used in Fig. 4E–H and Fig. 5 incorporate a leaky readout mechanism, consisting of dedicated output neurons corresponding to each task-specific label or behavior. These output neurons receive linear projections from the recurrent layer, which outputs normalized membrane potentials. In this setup, decoding is performed based on membrane potentials rather than neuronal spikes.

The following differential equation governs the temporal evolution of the readout neurons:

$$\tau_{\text{out}} \frac{d\mathbf{y}}{dt} = -\mathbf{y} + W^{\text{out}} \mathbf{r}^{\text{rec}} + b^{\text{out}}. \tag{S82}$$

In this equation, $\mathbf{y}$ represents the activity of output neurons, $\tau_{\text{out}}$ denotes the output neurons' time constant, $W^{\text{out}}$ signifies the synaptic weight matrix connecting recurrent and output neurons, $b^{\text{out}}$ is the bias term, and $\mathbf{r}^{\text{rec}}$ concatenates both membrane potentials and spike activities of the recurrent neurons. For computational implementations, a discrete-time approximation of this dynamics is employed:

$$\mathbf{y}^t = e^{-\Delta t/\tau_{\text{out}}} \mathbf{y}^{t-1} + (1 - e^{-\Delta t/\tau_{\text{out}}})(W^{\text{out}} \mathbf{r}^{\text{rec},t-1} + b^{\text{out}}). \tag{S83}$$

### J Resting-state neural activity of the *Drosophila* brain

In this section, we detail the resting-state calcium imaging data acquired from the *Drosophila* brain and outline the methodology for converting these calcium signals into deconvolved neuronal firing rates.

#### J.1 Resting-state whole-brain *in vivo* imaging data

Our whole-brain *in vivo* imaging data derive from experimental studies by Mann et al. (2017) [49] and Turner et al. (2021) [50]. They recorded resting-state neural activity in *Drosophila* using whole-brain calcium imaging of adult flies expressing the calcium-sensitive fluorescent protein GCaMP6s and membrane-tagged tdTomato. The central brain was exposed and imaged *in vivo* using resonant scanning two-photon microscopy at 3 $\mu$m isotropic resolution with 1.2 Hz temporal sampling for approximately 17 minutes per recording. They aligned functional data to a standard brain atlas using CMTK computational tools and defined regions of interest (ROIs) based on anatomical boundaries from established atlases (Ito or Branson). From each ROI, they extracted fluorescence time series, applied high-pass filtering to remove slow drift, and converted signals to $\Delta$F/F measurements. They quantified resting-state functional connectivity by calculating Pearson correlation coefficients between ROI time series, then applied Fisher Z-transformation for statistical analysis.

The exemplary traces of recording $\Delta$F/F can be obtained in Fig. 6A.

#### J.2 Deconvolution of calcium imaging data into neuropil firing rate

Challenges exist in obtaining $\Delta$F/F signals directly from simulated network models. To address this, we utilize a deconvolution-based approach to infer neuronal firing rates from calcium imaging data—specifically $\Delta$F/F signals—via methods described in prior work [51, 52]. This framework establishes a common metric for comparing experimental calcium imaging data with outputs from model simulations, enabling direct quantitative analysis across empirical and computational domains.

We formulate the problem within a linear convolution framework and employ regularized non-negative optimization to recover the underlying spike events, which are then converted to firing rates. The method accounts for the exponential decay characteristics of calcium transients and incorporates sparsity constraints to address the ill-posed nature of the deconvolution problem.

**Mathematical model** The relationship between neural spikes and the observed calcium fluorescence signal can be modeled as a linear convolution system. Let $s(t)$ represent the spike train (where each spike is a Dirac delta function), and $c(t)$ represent the calcium concentration signal. The dynamics can be described by:

$$c(t) = (h * s)(t) + \eta(t)$$

where $h(t)$ is the calcium impulse response function (typically modeled as an exponential decay), $*$ denotes convolution, and $\eta(t)$ represents noise. The impulse response function is often modeled as:

$$h(t) = Ae^{-t/\tau} \cdot H(t)$$

where $A$ is the amplitude, $\tau$ is the decay time constant, and $H(t)$ is the Heaviside step function.

In discrete time, with sampling interval $\Delta t$, this becomes:

$$c[n] = \sum_{k=0}^{n} h[n-k] \cdot s[k] + \eta[n]$$

With an exponential decay, the discrete impulse response becomes:

$$h[n] = Ae^{-n\Delta t/\tau} \quad \text{for } n \geq 0$$

For computational purposes, we reformulate the discrete convolution as a matrix operation. Let $\mathbf{c} = [c[0], c[1], \ldots, c[N-1]]^T$ be the vector of calcium measurements, and $\mathbf{s} = [s[0], s[1], \ldots, s[N-1]]^T$ be the vector of spike events to be estimated. The linear convolution can be expressed as:

$$\mathbf{c} = \mathbf{H}\mathbf{s} + \boldsymbol{\eta}$$

where $\mathbf{H}$ is an $N \times N$ lower triangular Toeplitz matrix with elements:

$$H_{ij} = \begin{cases} Ae^{-(i-j)\Delta t/\tau} & \text{if } i \geq j \\ 0 & \text{otherwise} \end{cases}$$

**Deconvolution approach** The deconvolution problem aims to recover the spike train $\mathbf{s}$ given the observed calcium signal $\mathbf{c}$ and knowledge of the calcium dynamics (matrix $\mathbf{H}$). This is an inverse problem that can be formulated as an optimization problem:

$$\min_{\mathbf{s}} \|\mathbf{H}\mathbf{s} - \mathbf{c}\|_2^2 \quad \text{subject to } \mathbf{s} \geq 0$$

However, this problem is often ill-posed due to noise and the low-pass filtering nature of calcium dynamics, making multiple spike patterns potentially consistent with the observed calcium signal.

To address the ill-posedness, we introduce regularization terms that incorporate prior knowledge about neural spike trains, particularly their sparsity. The L1-norm is commonly used to promote sparsity, leading to the following regularized optimization problem:

$$\min_{\mathbf{s}} \|\mathbf{H}\mathbf{s} - \mathbf{c}\|_2^2 + \lambda\|\mathbf{s}\|_1 \quad \text{subject to } \mathbf{s} \geq 0$$

where $\lambda > 0$ is the regularization parameter controlling the trade-off between the data fidelity term and the sparsity constraint.

The constrained optimization problem above can be solved using various numerical optimization algorithms. In our implementation, we use the L-BFGS-B algorithm [53], which is a limited-memory version of the Broyden–Fletcher–Goldfarb–Shanno algorithm with box constraints (to enforce non-negativity).

The objective function to be minimized is:

$$f(\mathbf{s}) = \|\mathbf{H}\mathbf{s} - \mathbf{c}\|_2^2 + \lambda\|\mathbf{s}\|_1 = \sum_{i=0}^{N-1} \left( \sum_{j=0}^{i} H_{ij} s_j - c_i \right)^2 + \lambda \sum_{i=0}^{N-1} |s_i|$$

With the non-negativity constraint $s_i \geq 0$ for all $i$, the absolute value in the L1-norm can be simplified to $|s_i| = s_i$.

Once the spike train $\mathbf{s}$ is estimated, it represents the magnitude of spike events at each time point. To convert this to a firing rate in Hz (spikes per second), we multiply by the sampling rate:

$$r[n] = s[n] \cdot f_s$$

where $r[n]$ is the firing rate at time point $n$, and $f_s = 1/\Delta t$ is the sampling frequency in Hz.

**Practical considerations** Prior to deconvolution, it is often beneficial to preprocess the calcium signal to reduce noise [54]. In our implementation, we apply Savitzky-Golay filtering, which performs polynomial regression on a moving window to smooth the data while preserving the shape of the signal.

Several parameters influence the performance of the deconvolution:

1. Decay time constant $\tau$: This should be set based on the calcium indicator used and the specific cell types under study. Typical values range from 0.5 to 2 seconds.

2. Regularization parameter $\lambda$: This controls the sparsity of the recovered spike train. Higher values lead to sparser solutions but may miss small events.
3. Sampling rate $f_s$: This determines the temporal resolution of the recovered spike train and should match the acquisition rate of the calcium imaging data.

### K Whole-brain *Drosophila* connectome-constrained model

In this section, we describe the whole-brain *Drosophila* connectome-constrained spiking network model.

#### K.1 Network architecture

The whole-brain spiking network contains more than 125,000 neurons and 50 million synaptic connections. We break down the network into several interconnected components (Fig. S4) that work together to generate and propagate spike events over time.

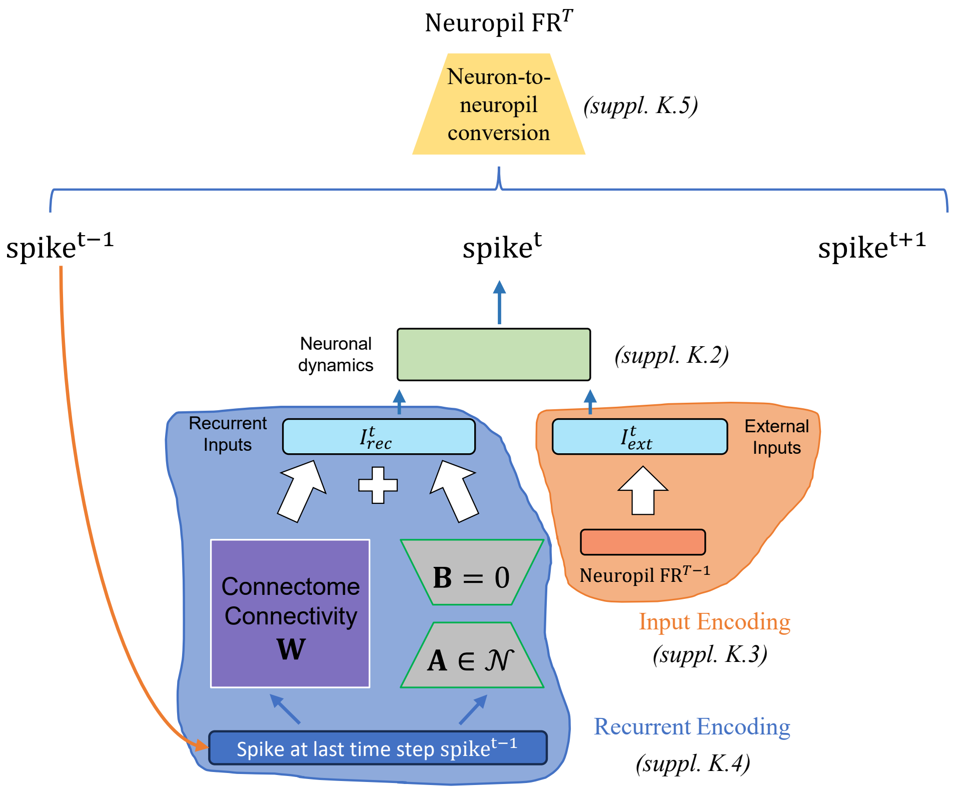

Figure S4: **Schematic diagram of the whole-brain *Drosophila* spiking network architecture**. The network processes information across discrete time steps, generating sequential spike outputs (spike$^{t-1}$, spike$^t$, spike$^{t+1}$) through an integrated system of external input processing and recurrent connectivity. External stimuli enter through the input encoding module (orange, detailed in SI. K.3), which converts continuous inputs ($I^t_{\text{ext}}$) into spike-compatible representations using previous neuropil firing rates (NeuropilFR$^{T-1}$). The recurrent processing core (blue) integrates both external inputs and internal feedback signals ($I^t_{\text{rec}}$) through the connectome connectivity matrix ($W$) and neuronal dynamics processor (green, SI. K.4). The system operates under mathematical constraints where bias terms B = 0 and activation parameters $A \in \mathcal{N}$. Temporal continuity is maintained through a recurrent encoding mechanism (SI. J.4) that processes spikes from the previous time step (spike$^{t-1}$) to generate current recurrent inputs. The final output stage employs a neuron-to-neuropil conversion process (yellow, supplemental J.5) that transforms individual spike events into population-level neuropil firing rates (NeuropilFR$^T$). Orange arrows indicate the recurrent connections that enable the network to maintain temporal memory and perform complex spatiotemporal computations. This architecture represents a biologically-inspired computational framework that integrates discrete spike-based processing with continuous population dynamics.

To fully understand the training process, it's essential to recognize that it operates on two distinct time scales. The first corresponds to the neuropil firing rate, sampled at 1.2 Hz—meaning each data point represents approximately 833 ms of biological time (see SI. J.1). We denote the neuropil firing rate at the $T$-th time step as $\mathrm{Neuropil\ FR}^T$.

In contrast, the simulated spiking neural network operates at a much finer temporal resolution, with a simulation time step of 0.2 ms. Thus, each single time step in the neuropil data spans $833/0.2 = 4165$ simulation steps. We denote the fine-grained simulation time as $t$, distinguishing it from the coarser neuropil time index $T$.

Within each simulation step $t$, we model the processing of both recurrent and external inputs in the spiking network.

The blue section illustrates the generation of recurrent input at time $t$, denoted by $I^t_{\text{rec}}$. This includes spikes produced at the previous simulation step $t-1$. Recurrent connectivity in the network comprises two components: (1) A structural connectivity matrix $W$, derived from the FlyWire connectome [55], representing canonical anatomical wiring in the template brain. (2) A low-rank adjustment component that accounts for biological variability and incomplete or noisy measurements in the connectome. This term captures weak or unobserved connections that are not directly evident in the dataset but may influence network dynamics. For details, refer to SI. K.4.

The brown section shows how the network encodes the previous neuropil firing rate, $\mathrm{Neuropil\ FR}^{T-1}$, into an external input at time $t$, denoted by $I^t_{\text{ext}}$. This transformation is performed by a gated recurrent unit (GRU) network [29]. See SI. K.3 for further information.

After completing all 4165 simulation steps corresponding to one neuropil time step, we compute the firing rate of each neuron in the whole-brain *Drosophila* network. To compare this output with the experimentally observed neuropil firing rate (which is convolved with calcium indicator dynamics), we apply a transformation algorithm to convert neuron-level firing rates into neuropil-level firing rates. Details of this procedure are provided in SI. K.5.

##### K.2 Network dynamics

The network dynamics builds upon the leaky integrate-and-fire framework developed by Shiu et al. (2024) [56]. The model incorporates leaky integrate-and-fire neurons coupled through exponential synapses, providing a biologically plausible representation of neural dynamics.

The continuous-time dynamics of the system are governed by the following differential equations:

$$T_{\text{mbr}} \frac{\mathrm{d}v_i}{\mathrm{d}t} = g_i - (v_i - V_{\text{resting}}) + I_{\text{ext}}, \tag{S84}$$

$$\frac{\mathrm{d}g_i}{\mathrm{d}t} = -\frac{g_i}{\tau} + I_{\text{rec}}, \tag{S85}$$

where $v_i$ represents the membrane potential of neuron $i$, and $g_i$ denotes the synaptic conductance resulting from aggregate presynaptic input to neuron $i$. $I_{\text{rec}}$ is the recurrent input current.

When the membrane potential reaches threshold ($v_i[n+1] \geq V_{\text{threshold}}$), the neuron fires a spike and the membrane potential is reset to $V_{\text{reset}}$, followed by a refractory period $T_{\text{refractory}}$ during which further spiking is suppressed.

The biophysical parameters are set to physiologically realistic values.

Membrane properties:

- Resting potential: $V_{\text{resting}} = -52$ mV
- Reset potential: $V_{\text{reset}} = -52$ mV
- Spike threshold: $V_{\text{threshold}} = -45$ mV
- Membrane resistance: $R_{\text{mbr}} = 10\mathrm{K}\Omega \cdot \mathrm{cm}^2$
- Membrane capacitance: $C_{\text{mbr}} = 2\mu\mathrm{F} \cdot \mathrm{cm}^{-2}$
- Membrane time constant: $T_{\text{mbr}} = C_{\text{mbr}} \times R_{\text{mbr}} = 20$ ms

Synaptic properties:

- Synaptic decay time constant: $\tau = 5$ ms

- Synaptic delay: $T_{\text{delay}} = 1.8$ ms
- Baseline synaptic weight: $W_{\text{syn}} = 0.275$ mV

Temporal properties:

- Refractory period: $T_{\text{refractory}} = 2.2$ ms

##### K.3 Input encoder

The input encoder transforms neuropil firing rates from the previous time step into background input for the current time step. Specifically, the encoder takes the neuropil firing rate at time step $T - 1$ (denoted as Neuropil FR$^{T-1}$) and generates the external input $I_{\text{ext}}^{T}$ for each neuron at time step $T$. All firing rates are measured in Hz.

During the $T$-th neuropil time step, neuron $i$ receives Poisson spike noise based on the transformed firing rate $\text{FR}_{\text{ext},i}^{T}$. To capture temporal dependencies between different time steps, we employ a Gated Recurrent Unit (GRU) [29] followed by a linear transformation as the encoder for the *Drosophila* whole brain model (Fig. S5). The transformation is given by:

$$\text{FR}_{\text{ext}}^{T} = \text{GRU}(\text{Neuropil FR}^{T-1}) \tag{S86}$$

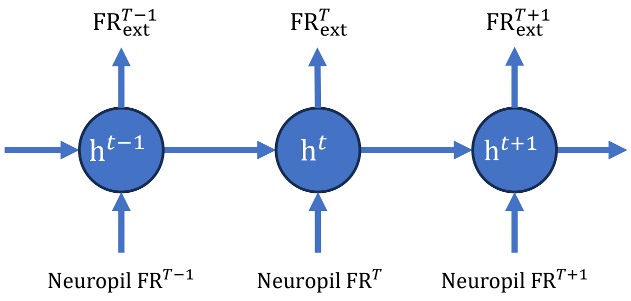

Figure S5: The input encoder using GRU+Linear network.

The `GRU+Linear` encoder processes sequential neuropil firing rates through a recurrent architecture where hidden states $h^{t-1}$, $h^{t}$, and $h^{t+1}$ maintain temporal information across time steps, each hidden state receives neuropil firing rates as input (Neuropil FR$^{T-1}$, Neuropil FR$^{T}$, Neuropil FR$^{T+1}$), the `GRU` gates regulate information flow and memory retention, the `Linear` output layers transform hidden states to external firing rates (FRext$^{T-1}$, FRext$^{T}$, $\text{FR}_{\text{ext}}^{T+1}$).

Then, the neuropil-time-scale firing rate is converted into Poisson noise, which is subsequently added to the membrane potential. Specifically, the external input current is modeled as:

$$I_{\text{ext}}(t) = \sum_{j} \delta(t - t_j) w_i, \tag{S87}$$

where $\delta(t - t_j)$ represents Poisson spike times and $w_i$ denotes the synaptic weight for neuron $i$. Note that the neuropil firing rate is operated at the neuropil time scale (see SI. K.1).

During training, Neuropil FR$^{T}$ corresponds to the experimentally recorded neuropil firing rate ExpFR$^{T}$; during inference, it refers to the simulated neuropil firing rate at the final time step, SimFR$^{T}$.

##### K.4 Recurrent encoder

The recurrent encoder converts neural spikes from the final time step into recurrent currents that influence subsequent neural dynamics. This recurrent connectivity architecture comprises two distinct components, as detailed in SI. K.1: first, anatomically-grounded connections derived from the template connectome [55], and second, learnable low-rank synaptic weights that capture functional connectivity patterns not present in the anatomical structure.

The total recurrent input to neuron $i$ at time $t$ combines both connectivity types:

$$I_{\mathrm{rec},i}(t) = \sum_j \delta(t - t_j^{\mathrm{spk}}) w_{j,i} + \sum_k \sum_j \delta(t - t_j^{\mathrm{spk}}) A_{k,i} B_{k,j} \tag{S88}$$

In the first term, the anatomical connection weight $w_{j,i}$ from presynaptic neuron $j$ to postsynaptic neuron $i$ is computed by scaling the Flywire connectome strength [55] with a sign factor ($+1$ for excitatory, $-1$ for inhibitory synapses) and a baseline synaptic weight parameter $W_{\mathrm{syn}}$.

The second term represents low-rank connectivity through factorized matrices $A$ and $B$, where the rank is set to 20. Unlike the fixed anatomical weights, these low-rank components are learned from scratch during training, enabling the model to capture functional connectivity patterns that emerge from task demands.

##### K.5 Computing neuropil firing rates from simulated single neuron activity

The overall firing rate of each *Drosophila* neuropil is estimated as a weighted average of its constituent neurons' firing rates. Because both neuronal distribution and synaptic connectivity vary across different regions, a simple arithmetic mean may misrepresent the true activity. Instead, each neuron's contribution is weighted by the number of synaptic spines it projects into the neuropil—information obtained from the Flywire connectome data [55]. This approach is formalized as:

$$\mathrm{SimFR}_i = \frac{\sum_j r_j \, w_j}{\sum_j w_j} \tag{S89}$$

where:

- $i$ denotes the $i$-th neuropil,
- $r_j$ is the firing rate of the $j$-th neuron,
- $w_j$ represents the weight, defined as normalized synaptic spines this neuron projects to the neuropil.

By emphasizing neurons with higher connectivity or located in denser regions, this method provides a more biologically accurate representation of the neuropil's collective activity.

### L Robust automatic online learning through the Jaxpr compilation

The backpropagation algorithm, a landmark in machine learning, has gained prominence due to its computational efficiency and seamless integration with modern automatic differentiation frameworks like PyTorch [57] and TensorFlow [58]. These frameworks simplify the implementation of backpropagation, allowing researchers and developers to focus on innovating and optimizing model architectures without delving into underlying details. Automatic differentiation systems efficiently and accurately perform gradient computations, further enhancing the algorithm's utility.

Inspired by the success of backpropagation, we aim to develop a general, scalable programming interface for online learning algorithms in SNNs. This interface allows to lower technical barriers by abstracting complex learning mechanisms, such as the automatic derivation and generation of eligibility traces and learning signals. Users need only define the network's dynamics and architecture, while the interface manages the subsequent online learning processes, including gradient calculations and parameter updates. Our design philosophy promotes the widespread adoption of SNN online learning algorithms across diverse model dynamics and architectures.

However, the development of such a framework presents three core challenges: (1) deriving eligibility traces, (2) generating learning signals, and (3) computing parameter gradients. To address these challenges, we leverage the Jaxpr language, an intermediate representation in JAX [59], which facilitates the inspection of computation graphs using abstract symbols. Without Jaxpr, analyzing the mathematical equations of user-defined models would be difficult. By abstracting user-defined Python code into manipulable symbols, we can derive and generate the necessary components for our online learning algorithms. Ultimately, this allows us to regenerate the Jaxpr code, encompassing both hidden state evolution and the online learning algorithms. We refer to this process as *online learning compilation*.

#### L.1 Deriving eligibility traces

For the automatic derivation of *eligibility traces*, our compilation consists of three steps.

The first step involves analyzing the hidden state group $\mathbf{h}_l$ that belong to a single recurrent layer. We aim to understand which state variables compose the hidden state $\mathbf{h}_l = [\mathbf{v}_l^1, \cdots, \mathbf{v}_l^d]$ (where $d$ is the number of variables in the hidden states), and how these state variables interact and transition into each other. This analysis is essential for computing the block diagonal matrix $\mathbf{D}_l^t = \begin{pmatrix} \partial \mathbf{v}_l^1/\partial \mathbf{v}_l^1 & \cdots & \partial \mathbf{v}_l^1/\partial \mathbf{v}_l^d \\ \vdots & \ddots & \vdots \\ \partial \mathbf{v}_l^d/\partial \mathbf{v}_l^1 & \cdots & \partial \mathbf{v}_l^d/\partial \mathbf{v}_l^d \end{pmatrix}$. This step yields multiple hidden state groups $[\mathbf{h}_1, \cdots \mathbf{h}_l, \cdots, \mathbf{h}_L]$, and their corresponding block diagonal matrices $\mathbf{D}_l^t$.

The second step involves analyzing how each weight parameter is connected to hidden state groups, requiring us to compute the weight-to-hidden relationships $\frac{\partial \mathbf{h}^t}{\partial \theta^t}$. By systematically enumerating the computation graph, we inspect which state variable $\mathbf{v}_l^j$ each parameter $\boldsymbol{\theta}_k$ is connected to, where $l$ is the index of the hidden state group, $j$ is the index of the variable in the hidden group $\mathbf{h}_l$, and $k$ is the index of the weight parameter. We then establish the relationship between the parameter $\boldsymbol{\theta}_k$ and the hidden state group $\mathbf{h}_l$. These dependencies are crucial for computing the block diagonal matrix $\mathbf{D}_f^t = \begin{pmatrix} \partial \mathbf{v}_l^1/\partial \boldsymbol{\theta} k \\ \vdots \\ \partial \mathbf{v}_l^d/\partial \boldsymbol{\theta}_k \end{pmatrix}$.

The third step involves generating the Jaxpr representation of eligibility traces based on the analysis results from the previous two steps. This process is crucial for translating our analytical insights into executable code. We leverage the symbolic Jaxpr language to generate mathematical operations as presented in Eqs. 4-8.

#### L.2 Generating learning signals

To generate the *learning signals* essential for Eq. 2, we currently employ the standard backpropagation algorithm through deep layers to maximize learning performance. This approach has proven effective in various deep-learning applications, allowing for the efficient propagation of error signals from the readout layer back through the hidden states.

However, implementing backpropagation through multiple hidden layers presents a significant challenge in the JAX framework. Specifically, it necessitates computing derivatives of gradients with respect to intermediate variables $[\mathbf{h}_1, \cdots \mathbf{h}_l, \cdots, \mathbf{h}_L]$. This operation is particularly difficult in JAX due to its inability to save gradients of intermediate variables. While JAX excels at gradient computation for explicit function parameters, hidden states computed within a function cannot be directly passed as function parameters, creating a bottleneck in the backpropagation process.

To overcome this limitation, we employ a perturbation function. Our approach involves adding a zero perturbation $\delta_{\mathbf{h}}$ to the computed hidden states, resulting in $\mathbf{h}^t + \delta_{\mathbf{h}}$. By treating this zero perturbation as a function parameter, we can effectively compute the required derivatives. This method capitalizes on the fact that the derivatives of the zero perturbation $\frac{\partial \mathcal{L}^t}{\partial \delta_{\mathbf{h}^t}}$ are mathematically equivalent to the derivatives of the corresponding hidden states $\frac{\partial \mathcal{L}^t}{\partial \mathbf{h}^t}$.

The practical implementation of this technique involves several key steps. We begin by rewriting the model's computation graph, incorporating a perturbation term into each hidden state: $\mathbf{h}^t \leftarrow \mathbf{h}^t + \delta_{\mathbf{h}^t}$. This modification allows us to introduce the perturbation without altering the network results. Then, utilizing the `jax.vjp` (vector-Jacobian product) interface provided by JAX, we compute the derivatives of the zero perturbations $\frac{\partial \mathcal{L}^t}{\partial \delta_{\mathbf{h}^t}}$. Finally, we extract this computation as a Jaxpr representation, allowing it to be leveraged in subsequent steps.

##### L.3 Computing parameter gradients

In the final stage of our algorithm, we integrate the eligibility traces and learning signals computed in previous steps to calculate parameter gradients. To ensure compatibility with standard automatic differentiation workflows, we use JAX's `jax.custom_vjp` to define a custom vector-Jacobian product (VJP) rule. This allows weight gradients at time $t$ to be accessed via standard interfaces such as `jax.grad`, while internally applying our online learning rules. This design enables seamless integration of our online gradient computation into JAX's differentiation framework.

Our implementation revolves around the definition of two critical functions:

1. Forward Pass Function: This function encapsulates the forward propagation of the network and the computation of eligibility traces by merging the Jaxpr representation of the two steps.
2. Backward Pass Function: This function is responsible for: a) Computing the learning signals $\frac{\partial \mathcal{L}^t}{\partial \mathbf{h}^t}$ based on the provided learning targets. b) Calculating the parameter gradients by combining the eligibility traces delivered from the forward pass and learning signals.

The integration of these two functions within the `jax.custom_vjp` interface allows us to take online gradient computation through the established JAX's autograd routines, such as `jax.grad` or `jax.vjp` functions. See Listing S4 for an example of BrainScale's online gradient computation.

Overall, the out-of-the-box interface for SNN online learning in BrainScale is remarkably simple, requiring no more than two lines of code. For example:

```
# define online computation using the ES-D-RTRL algorithm
model = brainscale.ES_D_RTRL(net, decay_or_rank=0.98)

# define online computation using the D-RTRL algorithm
model = brainscale.D_RTRL(net)

# compile the online learning graph using the input data
model.compile_graph(inputs)
```

Listing S1: The pseudo code to compile online learning implementations of the user-defined network `net`.

##### L.4 Optimizations during online learning compilation

During online learning compilation, we perform comprehensive optimizations to maximize computational performance and minimize memory overhead. These optimizations are crucial for scaling online learning algorithms to large-scale SNN architectures while maintaining real-time performance constraints. Our optimization strategy operates at multiple levels: computational graph analysis, memory management, and algorithmic efficiency. Currently, we implement the following key optimizations.

**Reusing spike eligibility traces.** In realistic neural circuit modeling, there are scenarios where the same presynaptic population projects to multiple post-synaptic populations through different synaptic connections. This architectural pattern is ubiquitous in biological neural networks, where a single neuron population often broadcasts its output to multiple downstream targets with varying connection weights and dynamics.

This structural redundancy presents a significant optimization opportunity. When the same presynaptic spike train $\mathbf{x}^t$ is used across multiple synaptic projections, the corresponding eligibility trace $\boldsymbol{\epsilon}_{\mathbf{x}}^t$ computed according to Eq. 7 remains identical regardless of the target population. Rather than recomputing these traces for each projection, our compilation framework automatically identifies shared presynaptic populations during graph analysis and caches their eligibility traces for reuse.

The implementation involves maintaining a registry of computed eligibility traces indexed by presynaptic population identifiers. During forward propagation, the system first checks whether the eligibility trace for a given presynaptic population has already been computed in the current time step. If so, the cached trace is retrieved; otherwise, it is computed once and stored for subsequent reuse. This optimization significantly reduces both computational overhead and memory allocation, particularly in densely connected networks where fan-out connections are common.

**Vectorizing hidden group Jacobian computations.** The computation of hidden-to-hidden Jacobians $\mathbf{D}_l^t = \frac{\partial \mathbf{h}_l^t}{\partial \mathbf{h}_l^{t-1}}$ and weight-to-hidden derivatives $\mathbf{D}_f^t = \frac{\partial \mathbf{h}_l^t}{\partial \boldsymbol{\theta}_k}$ can be computationally intensive. Naive implementations would compute these derivatives element-wise, resulting in numerous separate automatic differentiation calls. Our compilation framework addresses this inefficiency through comprehensive vectorization of Jacobian computations.

During graph analysis, we identify hidden state groups $\mathbf{h}_l$ that share similar computational patterns and can be processed in batches. The vectorization strategy operates on two levels:

Intra-group vectorization: For hidden states within the same group $\mathbf{h}_l = [\mathbf{v}_l^1, \cdots, \mathbf{v}_l^d]$, we compute the entire block diagonal matrix $\mathbf{D}_l^t$ in a single vectorized operation rather than computing individual partial derivatives $\partial \mathbf{v}_l^i / \partial \mathbf{v}_l^j$ separately. This is achieved by constructing the Jacobian computation as a batched matrix operation, leveraging JAX's efficient vectorized automatic differentiation primitives.

Inter-group vectorization: When multiple hidden groups share identical dynamics (common in layered architectures), we further batch Jacobian computations across groups. This involves reshaping the state tensors to enable simultaneous processing of multiple groups through broadcast operations.

The technical implementation utilizes JAX's `vmap` functionality to automatically vectorize over batch dimensions, combined with careful tensor reshaping to align dimensions for efficient computation. We also employ JAX's `scan` primitive for temporal dependencies, ensuring that recurrent computations maintain their sequential nature while benefiting from vectorization.

**Compilation-time optimizations.** Our online learning compilation framework also utilizes several general compile-time optimizations that improve the generated code efficiency:

- Dead code elimination: We analyze the computation graph to identify and remove unused calculations, particularly beneficial when users define complex network dynamics but only utilize specific learning algorithms.
- Operation fusion: Sequential operations that can be combined into single kernel calls are automatically fused during compilation, reducing memory bandwidth requirements and improving cache locality.

- Constant folding: Network parameters and hyperparameters that remain constant during learning are precomputed and embedded directly in the generated code, eliminating redundant calculations.

### M Comparison with other online learning algorithms

Online learning algorithms for spiking neural networks represent a rapidly evolving field that combines biologically inspired computation with practical machine learning applications. The field encompasses both supervised and unsupervised learning approaches, each addressing different aspects of temporal information processing and adaptation.

#### M.1 Unsupervised learning mechanisms

Unsupervised online learning algorithms for SNNs leverage fundamental biological principles to enable self-organization and feature extraction without labeled data.

Spike-timing dependent plasticity (STDP) [60] represents the cornerstone of unsupervised SNN learning. The mathematical formulation follows $\Delta w = A^+ \exp(-\Delta t/\tau^+)$ for $\Delta t > 0$ (LTP) and $\Delta w = -A^- \exp(\Delta t/\tau^-)$ for $\Delta t < 0$ (LTD), where the temporal relationship between pre- and postsynaptic spikes determines synaptic strength changes.

Triplet STDP [61] provides more biological accuracy by incorporating multiple spike interactions through the formulation $\Delta w = A_2^+ r_1(t) + A_3^+ r_2(t)$, where $r_1(t)$ and $r_2(t)$ are spike traces with different time constants. This mechanism enables temporal selectivity, competitive dynamics, and automatic feature extraction from spatiotemporal patterns.

Homeostatic plasticity [62] maintains network stability through multiple mechanisms including synaptic scaling, intrinsic plasticity, and metaplasticity. The mathematical foundation involves $\tau_{\text{scale}} \mathrm{d}w_i/\mathrm{d}t = \alpha(\rho_{\text{target}} - \rho_{\text{post}})w_i$ for synaptic scaling and $\tau_{\text{ip}} \mathrm{d}\theta/\mathrm{d}t = \mu(\rho_{\text{target}} - \rho_{\text{post}})$ for intrinsic plasticity.

Competitive learning [63] implements winner-takes-all mechanisms through lateral inhibition and synaptic competition. The implementation requires three essential components: Hebbian learning (through STDP), synaptic competition (synaptic forgetting), and neural competition (lateral inhibition). The mathematical formulation involves $\Delta w_{ij} = \eta^+ \times \text{pre\_trace}_j \times \text{post\_spike}_i$ for winning neurons and $\Delta w_{ij} = -\eta^- \times \text{pre\_trace}_j \times \text{lateral\_inhibition}$ for losing neurons. This creates specialized feature detectors and prevents multiple neurons from learning identical patterns.

Although shallow SNNs trained with STDP have demonstrated success in unsupervised visual classification tasks [64], plasticity-based learning algorithms remain inefficient for goal-directed learning. First, unsupervised rules like STDP are well-suited for capturing low-level statistical features—such as edges, orientations, and other input regularities—but lack mechanisms for optimizing high-level objectives like classification accuracy. Second, the performance of STDP is highly sensitive to hyperparameters: learning rate, temporal window ($\Delta t$), and synaptic bounds all require careful tuning. A temporal window that is too broad can cause neurons to fire synchronously in response to all inputs, while a window that is too narrow may fail to encode meaningful temporal correlations. Third, in multi-layer SNNs, STDP faces a fundamental credit assignment problem: synaptic updates in deeper layers cannot be directly linked to task-relevant errors. As the network depth increases beyond three layers, the spiking activity of output neurons often becomes decoupled from class-relevant input features, undermining the effectiveness of error-driven learning. As a result, plasticity-based training methods struggle to support efficient learning in complex tasks.

Typically, these unsupervised learning rules can be efficiently implemented in modern brain simulators, such as Brian2 [65], NEST [66], and BrainPy [26].

#### M.2 Early gradient-based online learning approaches

Supervised Hebbian learning (SHL) [67], ReSuMe [68], Tempotron [68], and Chronotron [69] represent early efforts to enable online supervised learning in spiking neural networks, each with distinct advantages and limitations.

The SHL framework [67] pioneered the integration of traditional Hebbian plasticity with external teaching signals, where an instructional input forces target neurons to spike at designated times while suppressing activity during non-target periods. This approach maintains biological plausibility through local synaptic modifications but remains constrained to single-layer architectures.

Building upon STDP principles, ReSuMe [68] introduced a more sophisticated learning mechanism where instructional signals modulate synaptic strength without directly altering membrane dynamics. While originally designed with online learning capabilities in mind, practical implementations typically employ fixed network topologies that lack structural adaptability to novel input patterns. The algorithm demonstrates particular strength in learning precise spatiotemporal spike patterns for applications such as neuroprosthetic control.

The Tempotron algorithm [70] represents a departure toward gradient-based optimization, employing traditional machine learning principles adapted for spiking neurons. This method trains LIF neurons to perform binary classification by learning to either generate action potentials or remain silent in response to input patterns. Despite its computational efficiency, the Tempotron's restriction to two-class problems significantly limits its practical applicability.

Addressing the Tempotron's limitations, the Chronotron [69] introduced dual learning modes: an offline E-learning variant that optimizes spike timing precision using distance metrics between target and actual spike trains, and an online I-learning approach that enables real-time adaptation through membrane potential tracking. This algorithm extends beyond binary classification to support multi-spike, multi-class learning scenarios, though at increased computational cost. Under specific parametric conditions, mathematical analysis reveals that Chronotron learning dynamics can reduce to Tempotron-equivalent behavior, establishing a theoretical connection between these approaches.

SHL is conceptually simple and biologically inspired, leveraging local activity correlations, but it lacks precise control over spike timing and struggles with complex temporal tasks. ReSuMe improves upon this by incorporating both target and actual spike timings through an STDP-like rule, enabling precise temporal learning in an online fashion; however, it is sensitive to spike jitter and assumes known desired spike trains. Tempotron offers robust online learning for binary spike/no-spike classification based on peak membrane potential, with efficient single-trial updates, but it is limited to single-spike decisions and does not scale well to complex temporal patterns. Chronotron, in contrast, is designed for precise multi-spike temporal pattern learning, optimizing spike timing globally; while powerful, it is more computationally demanding and typically requires access to the full output spike train, making it less suitable for strict online settings. Overall, these algorithms highlight the trade-off between biological plausibility, temporal precision, computational efficiency, and network architectures in online SNN learning.

##### M.3 Modern gradient-based online learning methods

Recent advances in SNN online learning have drawn inspiration from modern machine learning concepts developed for RNN training. A particularly promising alternative to BPTT is RTRL, which computes full gradients in a forward manner. However, RTRL's practical application is severely limited by its prohibitive $\mathcal{O}(N^3)$ memory complexity and $\mathcal{O}(N^4)$ computational complexity.

| Algorithm | Supported Models | Memory Complexity | Time Complexity |
|---|---|---|---|
| RTRL | General SNNs | $O(N^3)$ | $O(N^4)$ |
| OSTL | Not discussed | $O(N^2)$ | $O(N^2)$ |
| e-prop | Not discussed | $O(N^2)$ | $O(N^2)$ |
| OTTT | LIF | $O(N)$ | $O(N^2)$ |
| OTPE | LIF | $O(N)$ | $O(N^2)$ |
| NDOT | LIF | $O(N)$ | $O(N^2)$ |
| S-TLLR | LIF | $O(N)$ | $O(N^2)$ |
| **ES-D-RTRL (ours)** | AlignPre & AlignPost SNNs | $O(N)$ | $O(N^2)$ |

Table S1: Algorithm comparison of representative gradient-based online learning algorithms.

To address these computational bottlenecks, several approximation methods have been developed for RNN training. UORO [71] provides a rank-one approximation to RTRL but suffers from excessive noise in gradient estimates. More sophisticated approaches like KF-RTRL [35] and OK [72] offer low-rank approximations that reduce memory complexity to $\mathcal{O}(N^2)$, though they remain computationally expensive. SnAp [73] exploits parameter sparsity to reduce both computational and storage costs, but is limited to sparse RNN architectures and produces biased gradient computations.

Despite these advances in RNN training, the aforementioned methods have not been successfully adapted to SNN training due to their inherent limitations and the unique challenges posed by spiking dynamics. Instead, researchers have developed specialized approximations tailored specifically for spiking networks, which can be broadly categorized into two main approaches based on the complexity of the underlying neuron models.

**Approximations for general SNN models.** For general SNN architectures, two notable approaches have emerged with $\mathcal{O}(N^2)$ complexity. OSTL [43] simplifies the computation by assuming that all nonzero elements in the hidden state Jacobian lie along the diagonal, effectively treating neurons as independent units. In contrast, e-prop [33] takes a different approach by completely ignoring Jacobian information derived from recurrent connections, focusing instead on feedforward gradient propagation. However, neither algorithm explicitly characterizes the range of SNN architectures to which they can be applied. In principle, they can be applied to general SNN models defined in `AlignPre` and `AlignPost` abstractions.

**Approximations for simple neuron models.** For simplified neuron models such as the LIF neuron, researchers have achieved even greater computational efficiency. Methods including OTTT [74], OTPE [75], NDOT [76], and S-TLLR [77] have successfully derived online learning algorithms with $\mathcal{O}(N)$ memory complexity by exploiting the specific mathematical properties of these simplified neuron dynamics or ignoring some parts in gradient computations.

##### M.4 BrainScale: linear-memory complexity online learning algorithm for general SNNs

In general, *the field still lacks an online learning algorithm that can support the training of large-scale general SNN models with linear memory complexity*. Our ES-D-RTRL algorithm exactly fulfills this requirement. It shows several advantages over existing gradient-based, performance-driven online learning methods:

1. *Linear-memory complexity*: ES-D-RTRL reduces the memory complexity of online gradient computation from $\mathcal{O}(N^2)$ or higher to $\mathcal{O}(N)$, without relying on strong assumptions such as model sparsity or simplified neuron dynamics.

2. *Theory-guaranteed convergence and unbiasedness*: Unlike heuristic or biased approximations, ES-D-RTRL is derived from a principled estimator that guarantees unbiased gradient estimates under mild conditions.

3. *Compatibility with general neuron models*: ES-D-RTRL supports a wide range of spiking neuron dynamics, including models with complex neuronal, synaptic, or adaptive currents.

Together, these properties make ES-D-RTRL a strong candidate for scalable, gradient-based online learning in general-purpose spiking neural networks.

#### N Comparison with other frameworks

Currently, the fields of spiking neural network simulation, training, and neuromorphic computing have witnessed the emergence of numerous modeling frameworks, providing diverse technological pathways for brain simulation and brain-inspired computing.

| Category | Framework | Available Models | Hardware Support | Offline Gradients | Online Gradients | Plasticity Rules |
|---|---|---|---|---|---|---|
| Brain Simulation | NEURON | Multi - compartment neurons | CPU | ✗ | ✗ | ✗ |
| | NEST | LIF, EIF, etc. (point neurons) | CPU, MPI | ✗ | ✗ | STDP |
| | Brian/Brian2 | Custom ODE-based neurons/synapses | CPU, GPU | ✗ | ✗ | STDP |
| | Nengo | NEF-based, large-scale | CPU | ✗ | ✗ | NEF rules |
| Brain-inspired Computing | snnTorch | LIF | CPU, GPU | BPTT, ANN2SNN | ✗ | ✗ |
| | Norse | LIF, ALIF | CPU, GPU | BPTT | ✗ | ✗ |
| | Sinabs | LIF | CPU, GPU, Loihi | BPTT | ✗ | ✗ |
| | SpikingJelly | LIF | CPU, GPU | BPTT | ✗ | ✗ |
| | jaxsnn | LIF | BrainScaleS-2, CPU, GPU, TPU | EventProp | ✗ | ✗ |
| | SNNAX | LIF, ALIF | CPU, GPU, TPU | BPTT | ✗ | ✗ |
| | Spyx | LIF, ALIF | CPU, GPU, TPU | BPTT | ✗ | ✗ |
| | Lava-DL | LIF | Loihi | BPTT | DECOLLE | ✗ |
| Hybrid | BrainCog | Diverse neuron models | CPU, GPU | BPTT, ANN2SNN | ✗ | STDP |
| | **BrainPy (ours)** | Diverse neuron models | CPU, GPU, TPU | BPTT | ✗ | STDP |
| | **BrainScale (ours)** | Diverse neuron models | CPU, GPU, TPU | ✗ | ES-D-RTRL, D-RTRL, (designed for extensibility) | ✗ |

Table S2: Training functionality comparison of representative SNN frameworks.

##### N.1 SNN simulation and training

**Biologically plausible simulation frameworks.** NEURON [78] serves as a foundational tool in computational neuroscience, specializing in complex biological neuron modeling based on cellular morphology. It enables precise simulation for multi-compartment models and detailed ion channel dynamics. In contrast, NEST (Neural Simulation Tool) [66] is tailored for large-scale network simulation using point neuron models, capable of simulating networks with millions of neurons. Brian [79] and Brian2 [65] provide a flexible and efficient SNN modeling environment within the Python ecosystem. Their differential equation–based paradigm allows intuitive specification of diverse neuron and synapse models.

**Theoretically driven cognitive modeling frameworks.** Nengo [80] is built upon the neural engineering framework (NEF) and semantic pointer architecture (SPA) [81]. Nengo has enabled the construction of large-scale cognitive models, Spaun [82]. Spaun integrates 2.5 million spiking neurons and demonstrates complex cognitive behaviors such as image recognition, working memory, and question answering through multi-region coordination.

**Deep learning–inspired SNN frameworks.** With the rapid advancement of deep learning, several hybrid frameworks have emerged that combine deep neural networks with spiking mechanisms. Frameworks such as snnTorch [83], Norse [84], Sinabs [85], and SpikingJelly [86]—built on PyTorch [57]—support efficient training of deep SNNs via surrogate gradient methods and ANN-to-SNN conversion. Some of them [85, 83, 86], also show compatibility with neuromorphic deployment. These frameworks have shown significant performance improvements in applications like speech recognition, computer vision, and reinforcement learning. However, their design is primarily task-performance driven, with limited support for biological spiking architecture.

**Emerging frameworks in the JAX ecosystem.** Recent years have seen a surge of interest in JAX-based SNN frameworks, owing to JAX's powerful compilation optimizations and cross-platform hardware acceleration. jaxsnn [87] is an event-driven, machine-learning-inspired SNN training framework optimized for the BrainScaleS-2 neuromorphic backend. It integrates the EventProp algorithm [88] with a time-to-first-spike gradient strategy [89] for training spiking neural networks. The EventProp algorithm is a type of adjoint method that requires two passes: a forward pass for state transition and a backward pass for gradient computation. It is not online learning, but the memory only needs to store the state of the spiking moment (rather than the entire trajectory). SNNAX [90], built on JAX and Equinox, is a lightweight SNN framework combining PyTorch-style intuitive APIs with JAX-level performance. It leverages JAX's functional transformation features, including automatic differentiation, just-in-time (JIT) compilation, and vectorized operations, enabling efficient parallel computation on GPUs and TPUs. Spyx [91], developed with DeepMind's Haiku library, achieves near–custom CUDA kernel performance while maintaining PyTorch-like flexibility. Through GPU memory preloading and JIT optimization, Spyx achieves exceptional hardware efficiency in SNN computation.

**Integrated brain simulation and brain-inspired AI frameworks.** As seen above, frameworks differ markedly in their design philosophies, balancing biological plausibility and AI performance. Traditional tools like Nengo [80] and Brian/Brian2 [79, 65] excel at replicating experimental neuroscience phenomena but lag in performance on complex AI tasks. Conversely, performance-oriented frameworks like snnTorch [83] and SpikingJelly [86] achieve state-of-the-art results in AI benchmarks but offer limited biological interpretability.

BrainPy [26, 1] and BrainCog [92] stand out by striking a unique balance, aiming to unify accurate brain simulation with high-performance brain-inspired computing. Both platforms support biologically plausible modeling (e.g., diverse neuron models and learning rules) while achieving competitive results in real-world AI tasks. BrainPy [26], built on JAX, provides a unified programming interface for multiscale brain modeling—from single neurons to large-scale brain networks. It integrates model simulation, gradient-based training, and dynamical system analysis, featuring an extensive library of neuron models and plasticity mechanisms. BrainCog [92] systematizes composable components such as various biological neurons, encoding strategies, and learning rules. Technically, it offers comprehensive supervised and unsupervised training pipelines and plasticity rules. It also incorporates advanced techniques such as surrogate-gradient-based backpropagation and ANN-SNN conversion algorithms.

**Hardware support and deployment capabilities.** Traditional PyTorch-based frameworks mainly rely on GPU acceleration, whereas JAX-based systems natively support TPU and offer superior compilation optimization. Frameworks like jaxsnn [87] and Sinabs [85] extend their support to dedicated neuromorphic hardware, paving the way for co-design between software and hardware.

Of particular note is Lava-DL, which provides deep integration with Intel's Loihi architecture. It establishes a complete end-to-end pipeline from software training to neuromorphic hardware deployment. By enabling SLAYER-trained models [93] to be directly mapped onto Loihi chips, it realizes true end-to-end neuromorphic computation.

##### N.2 SNN online learning

As discussed above, most existing programming frameworks rely on BPTT-based offline learning algorithms for training spiking neural networks. However, this approach imposes significant limitations on the scalability of model training due to the high memory requirements of BPTT. In contrast, online learning—which maintains training memory only at a single time step—offers a promising solution for scalable SNN training.

**STDP-based online learning mechanisms.** Spike-timing-dependent plasticity (STDP), a Hebbian-inspired unsupervised learning rule, adjusts synaptic strengths based on the precise timing of pre- and post-synaptic spikes. Many modern frameworks implement STDP and its extended variants.

Brian/Brian2 [79, 65] excels in implementing biologically plausible STDP. Experiments demonstrate that small SNNs implemented in Brian2 can achieve an unsupervised visual classification task [64].

BrainCog [92] and BrainPy [26] provides a comprehensive suite of supervised and unsupervised learning methods, including STDP and other local plasticity rules.

Although Nengo [80] is grounded in NEF theory, it supports diverse learning rules, including the prescribed error sensitivity rule for online supervised learning. This method is especially effective for adaptive control systems, dynamically optimizing functional approximation via error minimization.

Nonetheless, STDP-based algorithms remain limited in their capacity to support efficient learning in large-scale, complex cognitive tasks.

**Hardware-coupled online learning.** Lava-DL offers deep integration with Intel's Loihi neuromorphic chip, supporting multiple operational modes including offline training, online learning, and efficient inference. The framework implements hardware-based online weight updates using the DECOLLE algorithm [94], though this capability is currently limited to LIF neurons.

Similarly, jaxsnn provides native support for the BrainScaleS-2 platform and enables on-chip learning through the EventProp algorithm. Despite this capability, the approach has notable limitations: first, EventProp is not a true online learning method and therefore cannot support real-time processing of streaming input data; second, like LAVA-DL, it supports only a restricted set of neuron models, primarily LIF neurons.

##### N.3 BrainScale: gradient-based online learning compilation for general SNNs

To date, the field still lacks a general-purpose software system tailored for high-performance online learning. BrainScale distinguishes itself through the following key innovations:

1. *Seamless integration with BrainPy*: BrainScale extends the capabilities of our BrainPy programming framework [26], enabling brain simulations augmented with powerful, gradient-based online learning support.

2. *Linear-memory ES-D-RTRL algorithm*: It implements the ES-D-RTRL algorithm, which approximates gradients while delivering strong training performance. Crucially, its linear-memory complexity eliminates the memory bottlenecks that have long hindered scalable online SNN training.

3. *Broad model compatibility*: The algorithm supports all SNN architectures defined using BrainPy's `AlignPre` and `AlignPost` abstractions—from simple LIF neurons to complex GIF models, and from basic exponential synapses to advanced short-term plasticity (STP) mechanisms.

4. *Automated online learning compilation*: BrainScale includes a compiler that automatically translates arbitrary user-defined models into efficient online learning code, leveraging IR analysis and code generation.

5. *Extensibility*: Importantly, this compilation capacity is general and can be easily extended to other online learning algorithms, such as eprop [33].

In summary, BrainScale goes beyond BrainPy's existing capabilities in simulation, offline training, and dynamical system analysis. It brings fully automated, scalable, and high-performance online learning to general SNNs. This system offers the same programming flexibility and comparable training performance as BPTT-based frameworks.

### O Environment settings

All evaluations and benchmarks in this study were conducted in a Python 3.12 environment on a system running the Ubuntu 24.04 LTS edition with CPU, GPU, and TPU devices.

The CPU experiments were run on an Intel Xeon W-2255, a 10-core/20-thread Cascade Lake processor with a base clock of 3.7 GHz and a turbo boost of 4.5 GHz. The CPU features 24.75 MB of L3 cache and supports up to 512 GB of six-channel DDR4-2933 ECC memory.

The GPU experiments used an NVIDIA RTX A6000, a professional Ampere GPU with 10,752 CUDA cores, 336 tensor cores, and 48 GB of GDDR6 memory. The card delivers 1 TB/s memory bandwidth, draws 300W power, and connects via PCIe 4.0x16, making it ideal for parallel computing and AI workloads.

The TPU experiments leveraged the Kaggle-free TPU v3-8 cloud instance. Specifically, the v3-8 instance gives 8 TPU v3 cores, each providing 128 GB/s of bandwidth to high-performance HBM memory.

### P Code examples of BrainScale programming interface

To demonstrate the versatility and practical utility of the BrainScale framework, we present three comprehensive case studies that illustrate model customization and gradient computation capabilities.

First, we implement a dense connection layer (Multi-Layer Perceptron) utilizing the `ETraceParam` concept, demonstrating parameter optimization in traditional neural architectures (Listing S2).

```python
import brainstate
import brainscale

class Linear(brainstate.nn.Module):
    def __init__(
        self,
        n_in: int,
        n_out: int,
        w_init=brainstate.init.KaimingNormal(),
        b_init=brainstate.init.ZeroInit()
    ):
        super().__init__()
        self.in_size = (n_in,)
        self.out_size = (n_out,)

        # parameters
        param = dict(weight=w_init([n_in, n_out]), bias=b_init([n_out]))
        # operations
        self.weight_op = brainscale.ETraceParam(param, brainscale.MatMulOp())

    def update(self, x):
        # call the model by ".execute" the weight_op
        return self.weight_op.execute(x)
```

Listing S2: Defining a dense connection layer in BrainScale.

Second, we define a Leaky Integrate-and-Fire (LIF) neuron model through the `ETraceState` concept, showcasing the framework's support for biological spiking neurons (Listing S3).

```python
import brainscale
import brainstate
import jax

class LIF(brainstate.nn.Neuron):
    def __init__(
        self,
        in_size,
        tau: float = 5.,
        V_th: float = 1.,
        V_reset: float = 0.,
        V_rest: float = 0.,
        spk_fun=brainstate.surrogate.ReluGrad(),
        spk_reset='soft'
    ):
        super().__init__(in_size, spk_fun=spk_fun, spk_reset=spk_reset)

        # parameters
        self.tau = tau
        self.V_th = V_th
        self.V_rest = V_rest
        self.V_reset = V_reset

    def dv(self, v, t, x):
        # "sum_current_inputs()" sums up all incoming currents
        x = self.sum_current_inputs(x, v)
```

```python
        # the differential equation
        return (-v + self.V_rest + x) / self.tau

    def init_state(self, batch_size: int = None, **kwargs):
        # initialize the membrane potential
        bs = () if batch_size is None else (batch_size,)
        V = jax.numpy.full(bs + self.varshape, self.V_rest)
        self.V = brainscale.ETraceState(V)

    def get_spike(self, V=None):
        V = self.V.value if V is None else V
        # scale the membrane potential
        v_scaled = (V - self.V_th) / self.V_th
        # generate spike using the surrogate gradient function
        return self.spk_fun(v_scaled)

    def update(self, x=0.):
        # the last spike and membrane potential
        last_v = self.V.value
        lst_spk = self.get_spike(last_v)
        if self.spk_reset == 'soft':
            V_th = self.V_th
        else:
            V_th = jax.lax.stop_gradient(last_v)
        V = last_v - (V_th - self.V_reset) * lst_spk
        # the current membrane potential
        V = brainstate.nn.exp_euler_step(self.dv, V, None, x)
        V = self.sum_delta_inputs(V)
        self.V.value = V
        # the current spike
        return self.get_spike(V)
```

Listing S3: Defining a LIF neuron model in BrainScale.

Third, we integrate these components to construct a spiking network and compute gradients for network parameters based on specified learning objectives, illustrating end-to-end differentiable training (Listing S4).

```python
import brainscale
import brainstate
import braintools
import jax
import jax.numpy as jnp

class LIF_Delta_Net(brainstate.nn.Module):
    def __init__(
        self,
        n_in: int,
        n_rec: int,
        n_out: int,
        tau_mem: float = 5.,
        V_th: float = 1.,
    ):
        super().__init__()
        self.neu = LIF(n_rec, tau=tau_mem, V_th=V_th)
        self.syn = brainstate.nn.DeltaProj(comm=Linear(n_in + n_rec, n_rec), post=self.neu)
        self.out = brainscale.nn.LeakyRateReadout(n_rec, n_out, tau=5.0)

    def update(self, i, spk):
        with brainstate.environ.context(i=i, t=i * brainstate.environ.get_dt()):
            spk = jnp.concat([spk, self.neu.get_spike()], axis=-1)
```

```
            self.syn(spk)
            return self.out(self.neu())

brainstate.environ.set(dt=1.0)

# define the one-batch inputs and targets
n_seq = 512
inputs = brainstate.random.rand(n_seq, 10) < 0.1
targets = brainstate.random.randint(0, 2)

# instantiate a spiking network
net = LIF_Delta_Net(10, 100, 2)
brainstate.nn.init_all_states(net)

# online learning algorithm
method = 'es-diag'  # or 'diag', 'hybrid'
if method == 'es-diag':
    model = brainscale.ES_D_RTRL(net, decay_or_rank=0.98)
elif method == 'diag':
    model = brainscale.D_RTRL(net)
elif method == 'hybrid':
    model = brainscale.HybridDimVjpAlgorithm(net, decay_or_rank=0.98)
else:
    raise ValueError(f'Unknown online learning methods: {method}.')

# compile the eligibility trace graph using the one-step input
model.compile_graph(0, inputs[0])

# retrieve parameters that need to compute gradients
weights = net.states().subset(brainstate.ParamState)

# define loss function
def loss_fn(i, x):
    out = model(i, x)
    return jnp.mean(braintools.metric.
    softmax_cross_entropy_with_integer_labels(out, targets))

# gradient computation using traditional autograd interface
def step_grad(last_grads, ix):
    # gradients computed at the current step
    grads = brainstate.transform.grad(loss_fn, weights)(ix[0], ix[1])
    # accumulate gradients: prev + current
    new_grads = jax.tree.map(jax.numpy.add, last_grads, grads)
    return new_grads, None

# loop over all sequences
indices = jax.numpy.arange(n_seq)
init_grads = jax.tree.map(jax.numpy.zeros_like, weights.to_dict_values
    ())
grads = brainstate.transform.scan(step_grad, init_grads, (indices,
    inputs))
```

Listing S4: Defining a network in BrainScale in which LIF neurons (defined in Listing S3) are interconnected with Delta synapses (see the linear transformation in Listing S2).

These examples collectively demonstrate BrainScale's unified programming interface, architectural flexibility, and extensibility, enabling researchers and practitioners to seamlessly design, implement, and train diverse spiking neural models within a single, cohesive framework.

### Q Supplementary figures

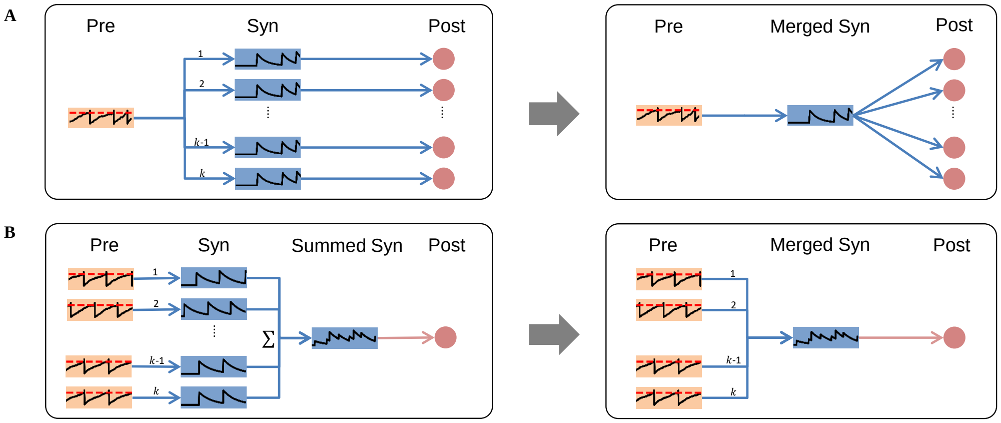

Figure S6: `AlignPre` **and** `AlignPost` **SNN abstractions**. (A) `AlignPre` abstraction models presynaptic neuron dynamics and is compatible with any synapse model. (B) `AlignPost` abstraction models postsynaptic neuron dynamics and is specifically designed for exponential-family synapse models. Orange represents the presynaptic neuron, blue represents synaptic dynamics, and red represents the postsynaptic neuron.

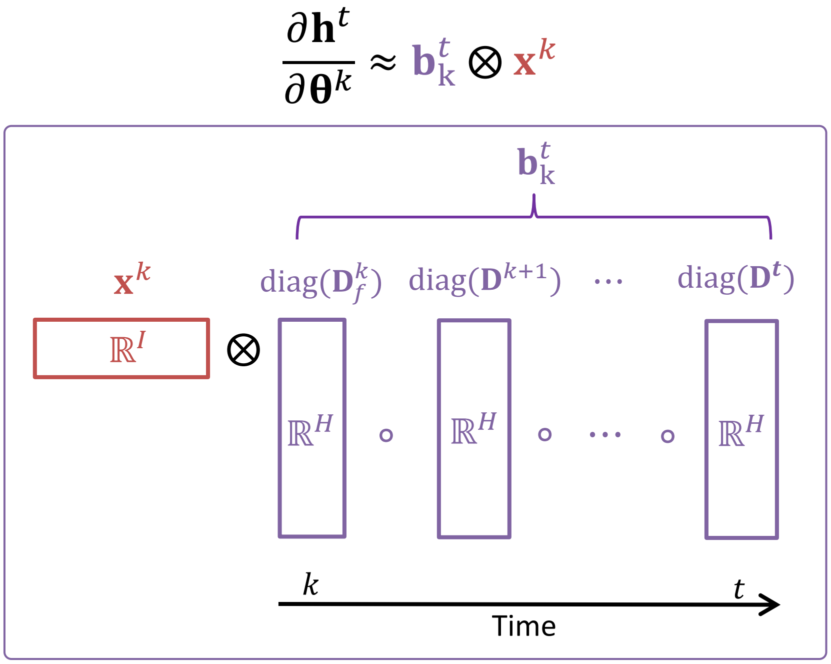

Figure S7: **Computation of the Eligibility trace $\frac{\partial \mathbf{h}^t}{\partial \boldsymbol{\theta}^k}$ in D-RTRL and ES-D-RTRL**. See the main text in Section .

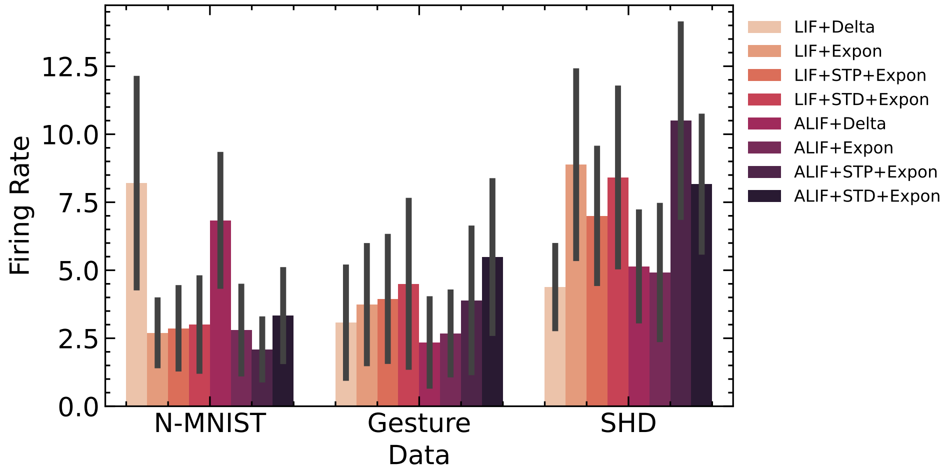

Figure S8: **Firing Rate Statistics During Jacobian and Gradient Evaluation**. This figure presents the firing rate statistics observed during the evaluation of Jacobian and gradient computations in Fig. 4A-D across various network dynamics and neuromorphic datasets.

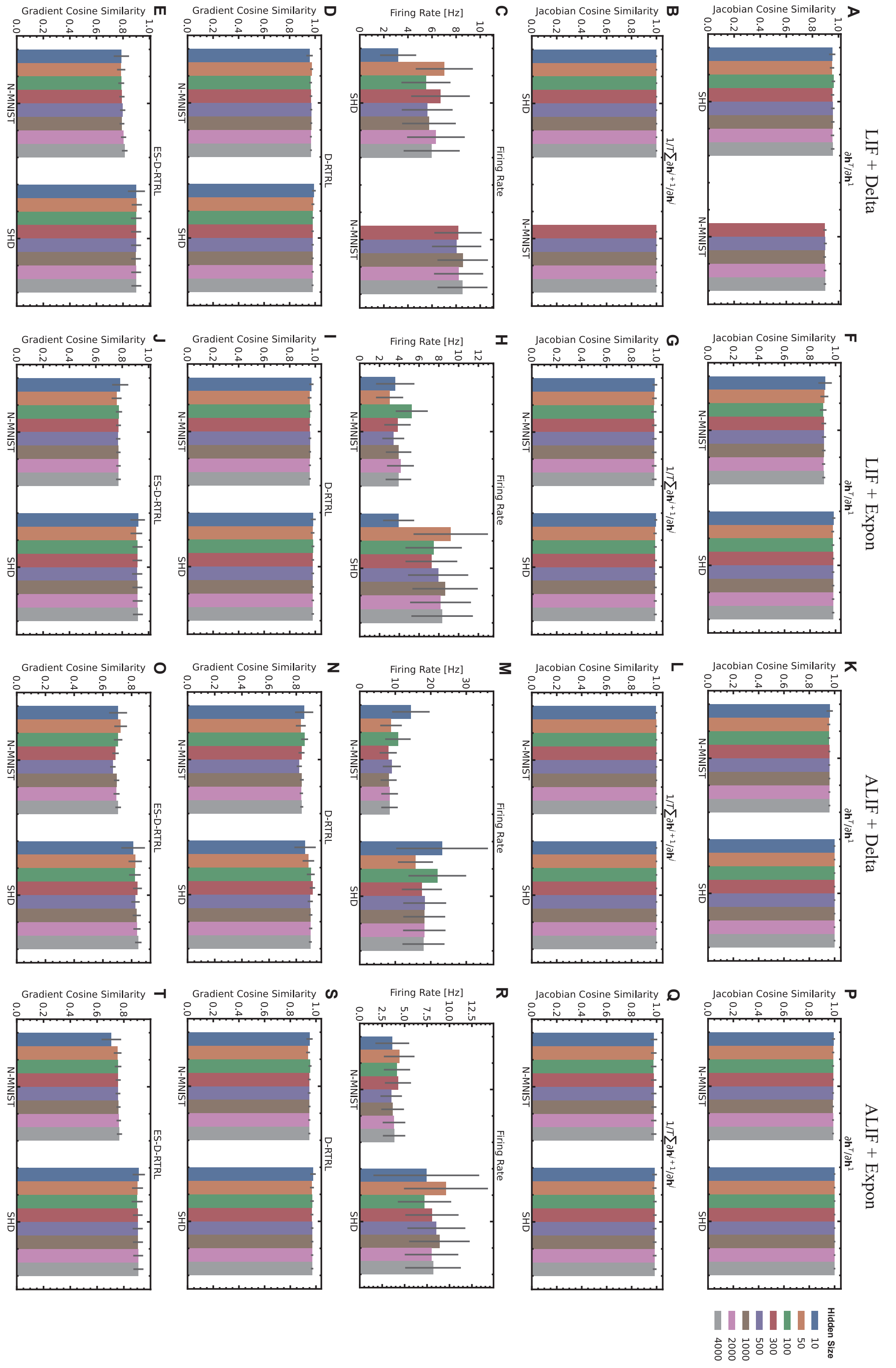

Figure S9: **Impact of Network Size on Cosine Similarity of Hidden Jacobian and Weight Gradients in Recurrent SNNs.**

Figure S9: This figure presents a comprehensive analysis of how network size affects the cosine similarity of hidden Jacobian and weight gradients in various SNN configurations. Experiments were conducted using single recurrent layer architectures across different network sizes, utilizing the SHD [44] and N-MNIST [95] datasets. Each data point in the box plots represents the average of repeated experiments conducted on each dataset composed of 500 sampled data points, with 95% confidence intervals shown. The figure is organized into four columns and five rows, exploring different neuron dynamics and synapse models. *First Column (A-E)*: Evaluations by using LIF neuron dynamics connected with Delta synapse models. *Second Column (F-J)*: Evaluations by using LIF neuron dynamics connected with Exponential synapse models. *Third Column (K-O)*: Evaluations by using ALIF neuron dynamics connected with Delta synapse models. *Fourth Column (P-T)*: Evaluations by using ALIF neuron dynamics connected with Exponential synapse models. *First Row (A, F, K, P)*: The cosine similarity of long-term hidden Jacobian $\frac{\partial \mathbf{h}^T}{\partial \mathbf{h}^1}$. *Second Row (B, G, L, Q)*: The average cosine similarity of one-step hidden Jacobian $1/T \sum_{j=1}^{T} \frac{\partial \mathbf{h}^j}{\partial \mathbf{h}^{j-1}}$. *Third Row (C, H, M, R)*: The firing rate of the simulated single-layer SNN network. *Fourth Row (D, I, N, S)*: The cosine similarity of the weight gradients between BPTT and D-RTRL. *Fifth Row (E, J, O, T)*: The cosine similarity of the weight gradients between BPTT and ES-D-RTRL.

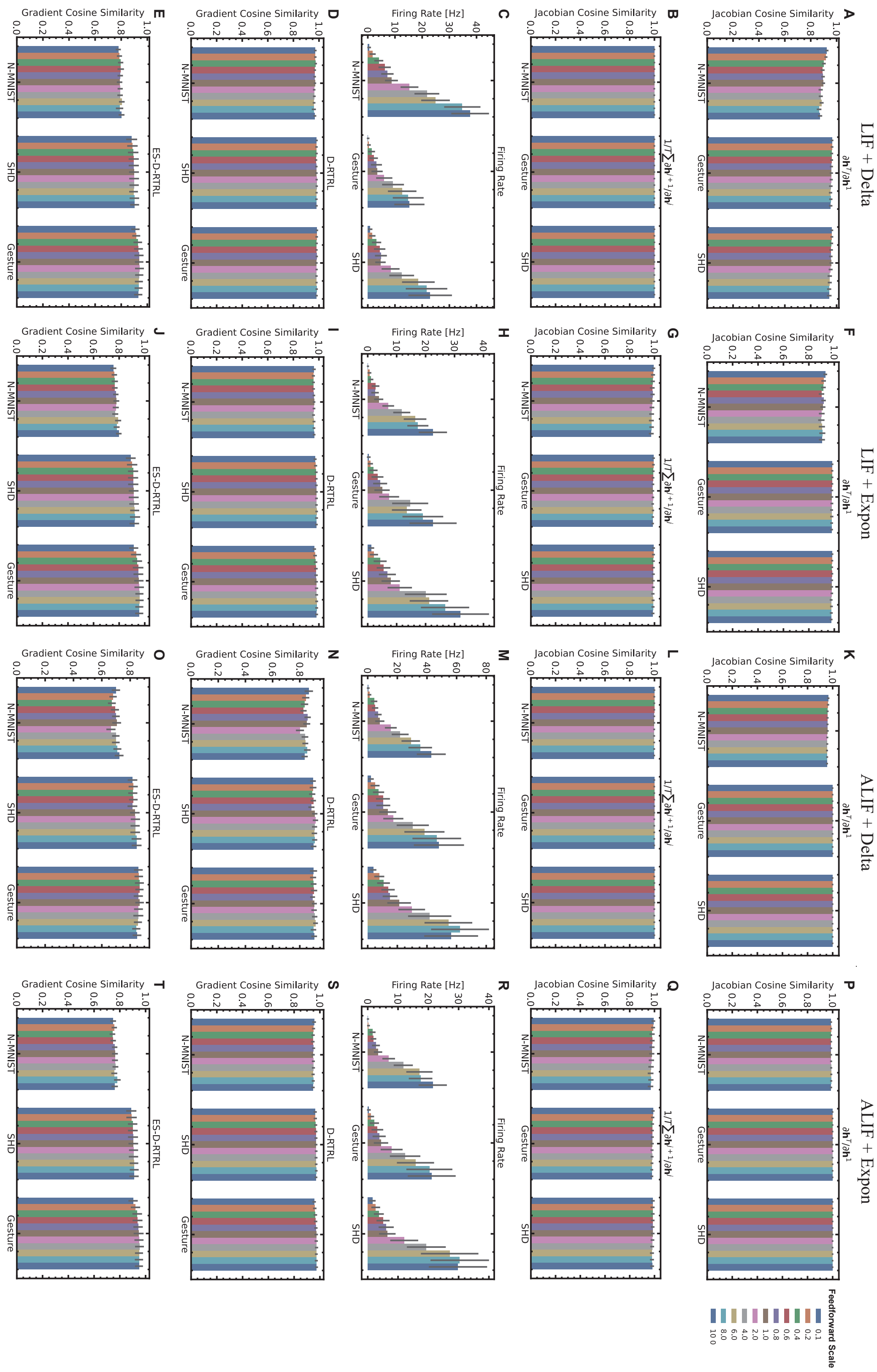

Figure S10: **Impact of Feedforward Connection Size on Cosine Similarity of Hidden Jacobian and Weight Gradients in Recurrent SNNs.**

Figure S10: This figure presents a comprehensive analysis of how feedforward connection size affects the cosine similarity of hidden Jacobian and weight gradients in various SNN configurations. Experiments were conducted using single recurrent layer architectures across different feedforward connection scales, utilizing the N-MNIST [95], Gesture [36], and SHD [44] datasets. Each data point in the box plots represents the average of repeated experiments conducted on each dataset composed of 500 sampled data points, with 95% confidence intervals shown. The figure is organized into four columns and five rows, exploring different neuron dynamics and synapse models. *First Column (A-E)*: Evaluations by using LIF neuron dynamics connected with Delta synapse models. *Second Column (F-J)*: Evaluations by using LIF neuron dynamics connected with Exponential synapse models. *Third Column (K-O)*: Evaluations by using ALIF neuron dynamics connected with Delta synapse models. *Fourth Column (P-T)*: Evaluations by using ALIF neuron dynamics connected with Exponential synapse models. *First Row (A, F, K, P)*: The cosine similarity of long-term hidden Jacobian $\frac{\partial \mathbf{h}^T}{\partial \mathbf{h}^1}$. *Second Row (B, G, L, Q)*: The average cosine similarity of one-step hidden Jacobian $1/T \sum_{j=1}^{T} \frac{\partial \mathbf{h}^j}{\partial \mathbf{h}^{j-1}}$. *Third Row (C, H, M, R)*: The firing rate of the simulated single-layer SNN network. *Fourth Row (D, I, N, S)*: The cosine similarity of the weight gradients between BPTT and D-RTRL. *Fifth Row (E, J, O, T)*: The cosine similarity of the weight gradients between BPTT and ES-D-RTRL.

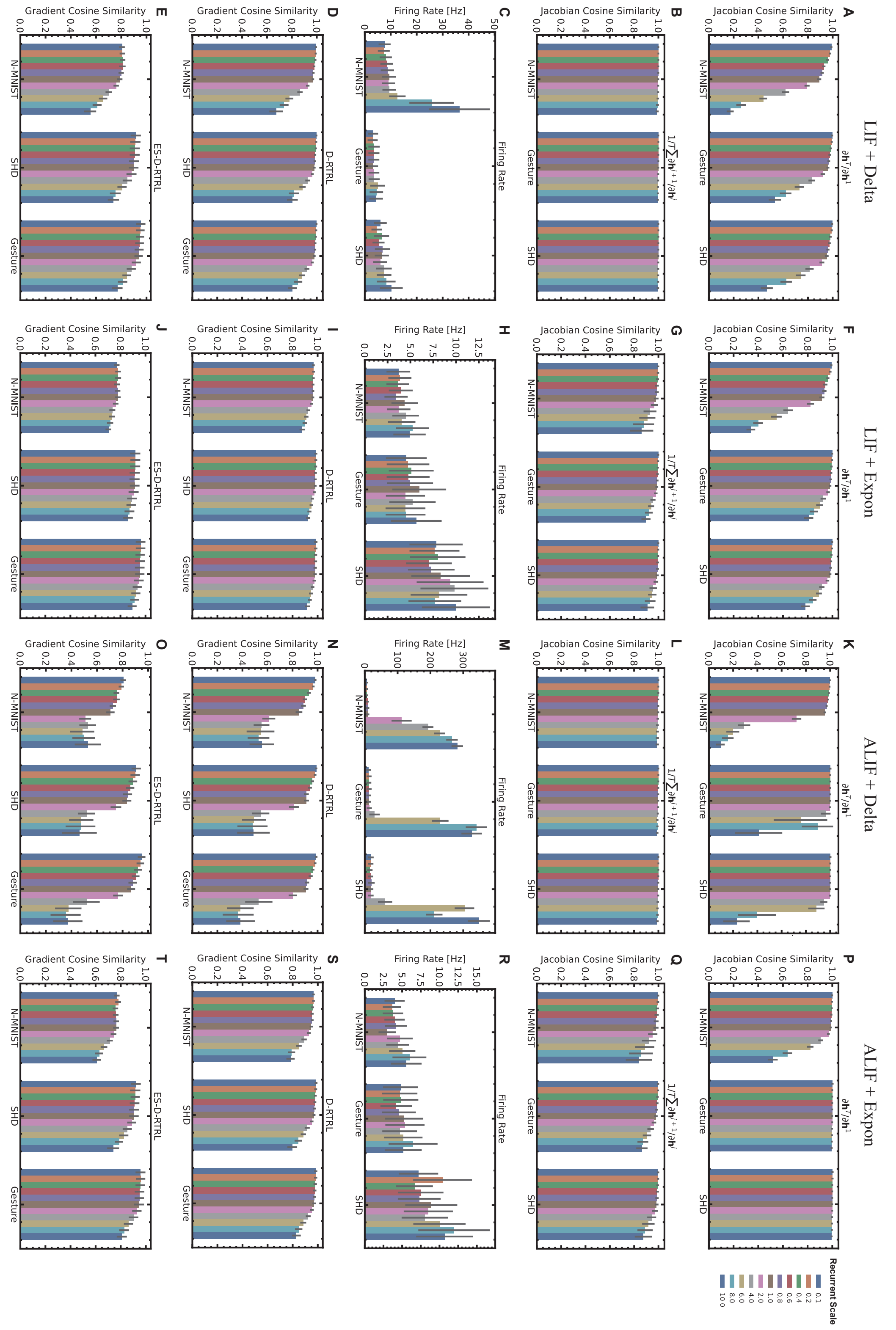

Figure S11: **Impact of Recurrent Connection Size on Cosine Similarity of Hidden Jacobian and Weight Gradients in Recurrent SNNs.**

Figure S11: This figure presents a comprehensive analysis of how recurrent connection size affects the cosine similarity of hidden Jacobian and weight gradients in various SNN configurations. Experiments were conducted using single recurrent layer architectures across different recurrent connection scales, utilizing the N-MNIST [95], Gesture [36], and SHD [44] datasets. Each data point in the box plots represents the average of repeated experiments conducted on each dataset composed of 500 sampled data points, with 95% confidence intervals shown. The figure is organized into four columns and five rows, exploring different neuron dynamics and synapse models. *First Column (A-E)*: Evaluations by using LIF neuron dynamics connected with Delta synapse models. *Second Column (F-J)*: Evaluations by using LIF neuron dynamics connected with Exponential synapse models. *Third Column (K-O)*: Evaluations by using ALIF neuron dynamics connected with Delta synapse models. *Fourth Column (P-T)*: Evaluations by using ALIF neuron dynamics connected with Exponential synapse models. *First Row (A, F, K, P)*: The cosine similarity of long-term hidden Jacobian $\frac{\partial \mathbf{h}^T}{\partial \mathbf{h}^1}$. *Second Row (B, G, L, Q)*: The average cosine similarity of one-step hidden Jacobian $1/T \sum_{j=1}^{T} \frac{\partial \mathbf{h}^j}{\partial \mathbf{h}^{j-1}}$. *Third Row (C, H, M, R)*: The firing rate of the simulated single-layer SNN network. *Fourth Row (D, I, N, S)*: The cosine similarity of the weight gradients between BPTT and D-RTRL. *Fifth Row (E, J, O, T)*: The cosine similarity of the weight gradients between BPTT and ES-D-RTRL.

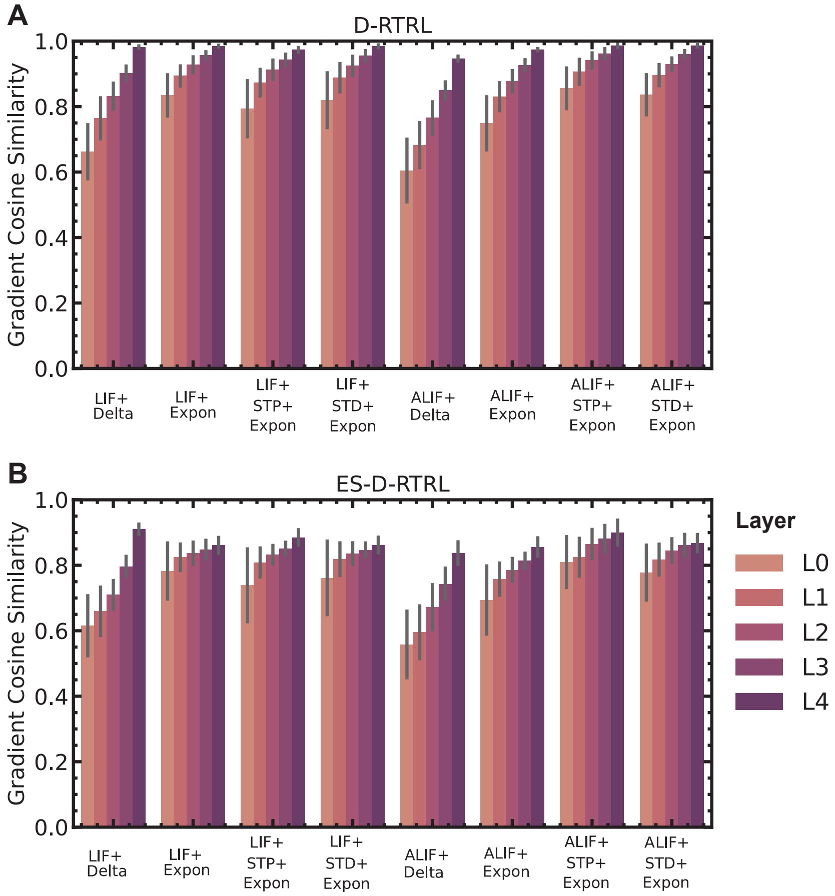

Figure S12: **Evaluation of Gradient Approximation Accuracy Over Different Layers**. We examined the impact of the depth of recurrent layers on the approximation of weight gradients in deep recurrent layers. We assessed the layer-wise cosine similarity between online gradient estimates and offline BPTT gradients using the SHD dataset [44]. Specifically, we evaluated a five-layer SNN network with 200 hidden units per layer, considering a variety of neuronal and synaptic dynamics (SI. B). `L0` represents the bottom layer, and `L4` denotes the top layer. Consistently, we observed a decrease in cosine similarity as the layer depth increased (see panel A for the D-RTRL algorithm and panel B for the ES-D-RTRL algorithm). Lower layers exhibited lower cosine similarity to their exact gradients, corresponding to the fact that more recurrence information of hidden states in top layers is ignored. (A) Layer-wise cosine similarity between weight gradients computed by D-RTRL and BPTT algorithms. (B) Layer-wise cosine similarity between weight gradients computed by ES-D-RTRL and BPTT algorithms.

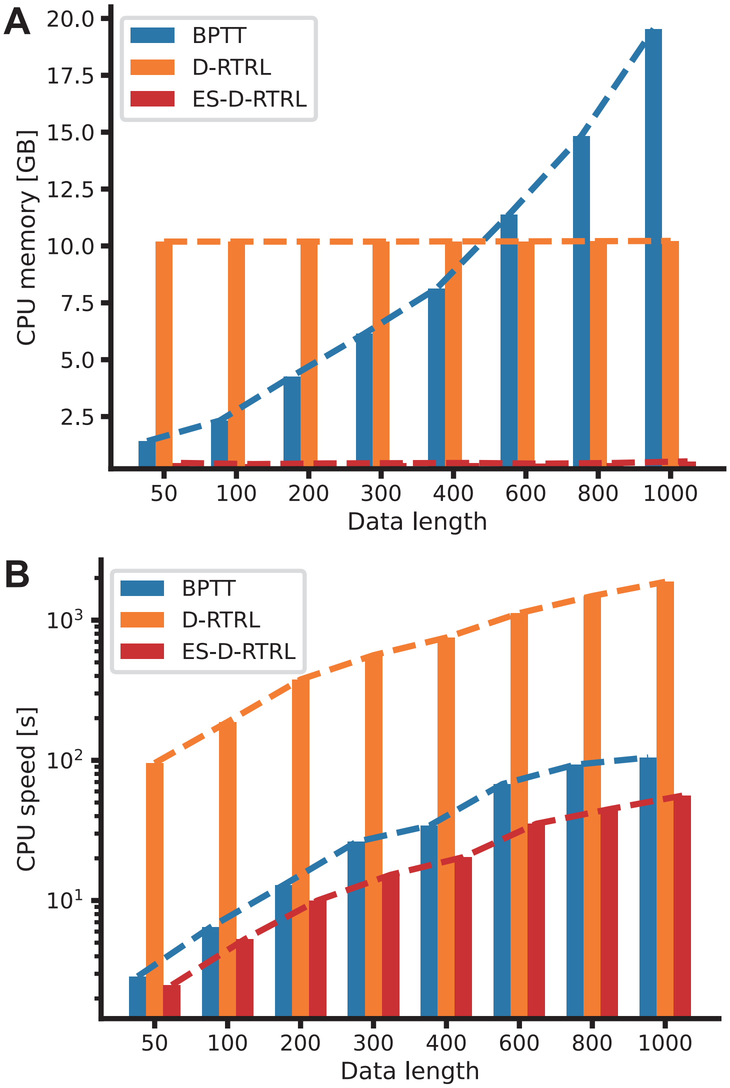

Figure S13: **Comparison of Memory Consumption and Computational Speed on Intel CPU Devices**. All experiments were conducted on the Gesture dataset [36] with varying sequence lengths from 50 to 1000 time steps. Same as Fig. 4 I–J, the batch size here is set to 128. (A) Memory consumption per batch for three training algorithms: BPTT, D-RTRL, and ES-D-RTRL. (B) Computation time per batch for the same three algorithms on CPU devices.

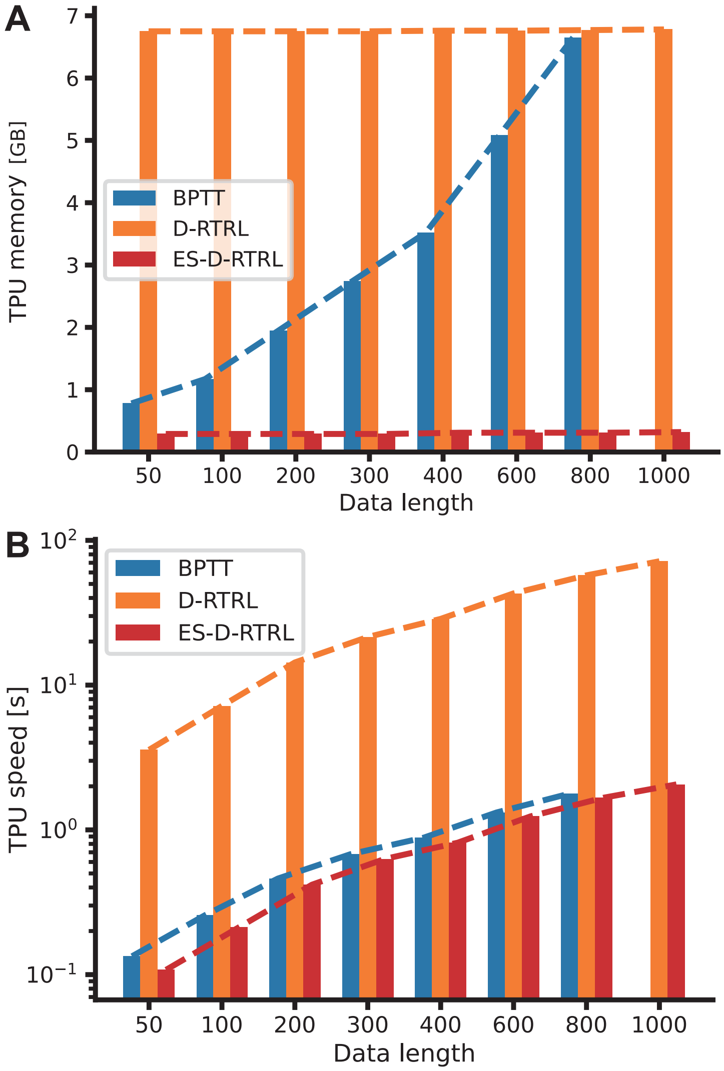

Figure S14: **Comparison of Memory Consumption and Computational Speed on TPU-v3 Devices**. All experiments were conducted on the Gesture dataset [36] with varying sequence lengths from 50 to 1000 time steps. Unlike Fig. 4 I–J, the batch size here is set to 32 due to the 8 GB memory limitation of a single TPU-v3 chip. Note that BPTT encounters an out-of-memory error when processing sequences longer than 800 time steps. (A) Memory consumption per batch for three training algorithms: BPTT, D-RTRL, and ES-D-RTRL. (B) Computation time per batch for the same three algorithms on TPU-v3.

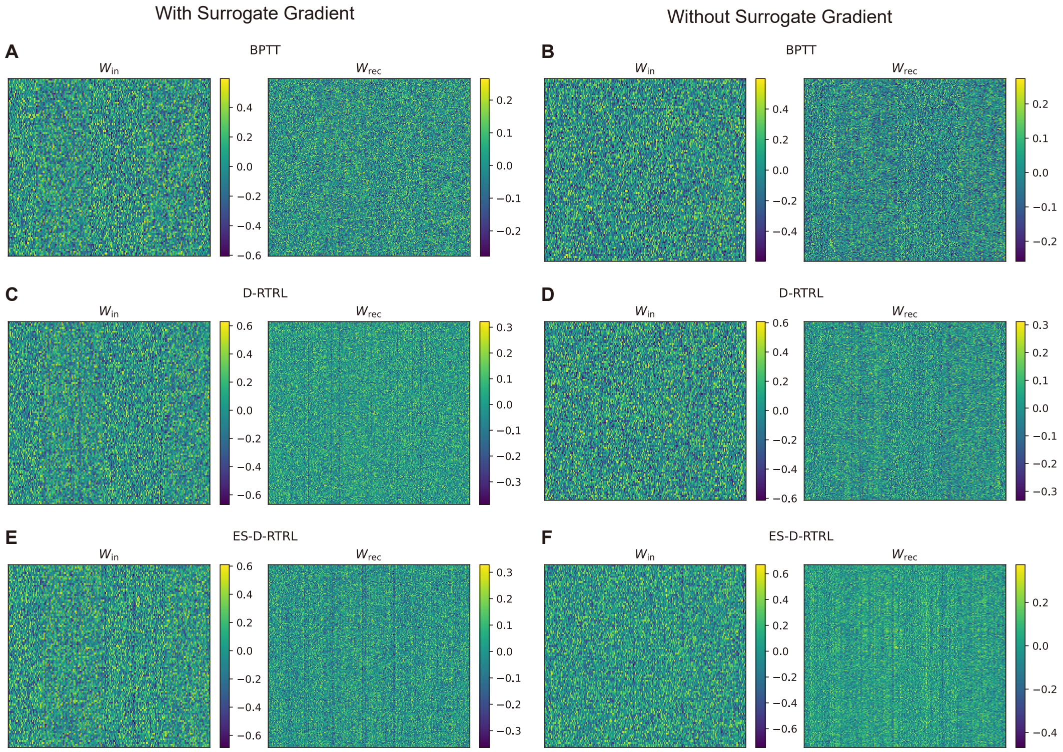

Figure S15: **Comparison of Weight Distributions in Spiking Networks Trained on the DMTS Task using Different Algorithms**. This figure presents weight visualizations for spiking networks trained on the DMTS task using three distinct algorithms: BPTT, D-RTRL, and ES-D-RTRL. The comparison is made both with and without the use of surrogate gradients. (A-B) Weights trained using BPTT. (C-D) Weights trained using D-RTRL. (E-F) Weights trained using ES-D-RTRL. Left column (A, C, E): Networks trained with surrogate gradients. Right column (B, D, F): Networks trained without surrogate gradients. Each panel shows a heatmap or color-coded representation of weight matrices. This visualization allows for a qualitative assessment of how different training algorithms and the use of surrogate gradients affect the final weight configurations in spiking neural networks trained on the DMTS task.

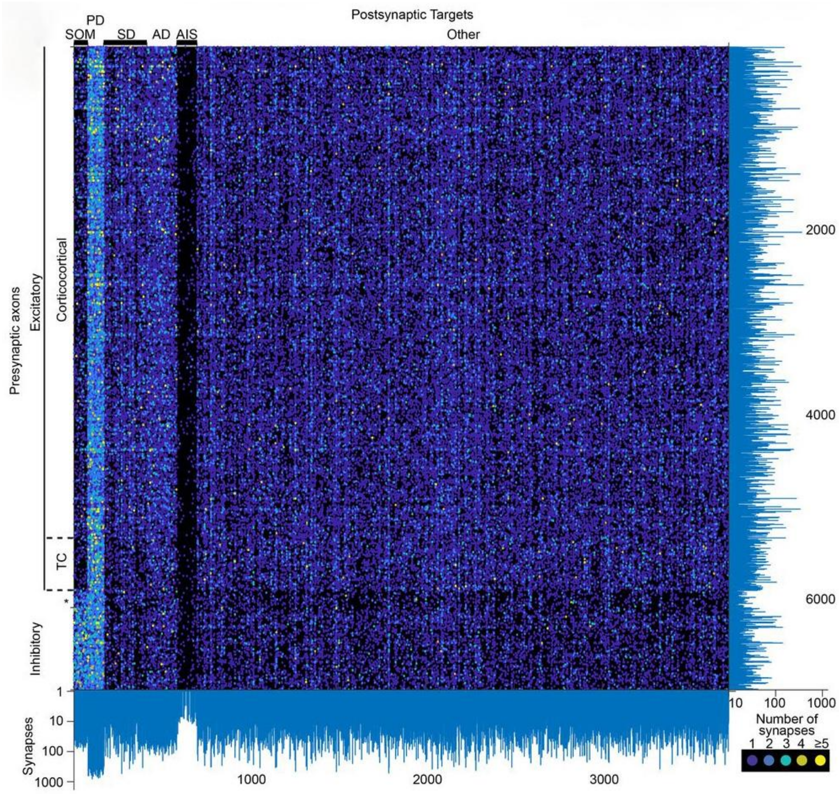

Figure S16: Layer 4 somatosensory cortical connectome between axons and postsynaptic targets [96]. This panel, adapted from Fig. 3E in [96], illustrates the connectome between all axons (n = 6,979) and postsynaptic targets (n = 3,719) within the volume, each with at least 10 synapses. A total of 153,171 synapses are established out of the 388,554 synapses detected in the volume. Please compare this panel with the data shown in Fig. S15.

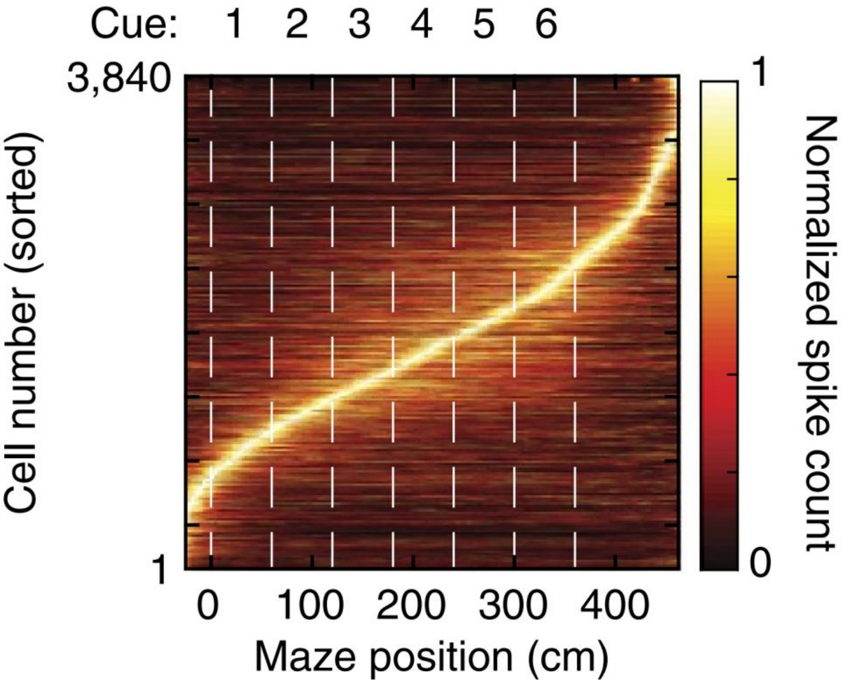

Figure S17: Sorted neuron activity in a head-restrained mouse performing a navigation-based evidence accumulation task [48]. This panel, adapted from Fig. 2a in [48], shows the normalized mean activity across all trials for all neurons in the posterior parietal cortex, pooled across all data sets (n = 3,840 cells from 5 mice). Traces were normalized to the peak of each cell's activity, averaged, and sorted by the peak's maze position. Please compare this panel with the data shown in Fig. 5E-F.

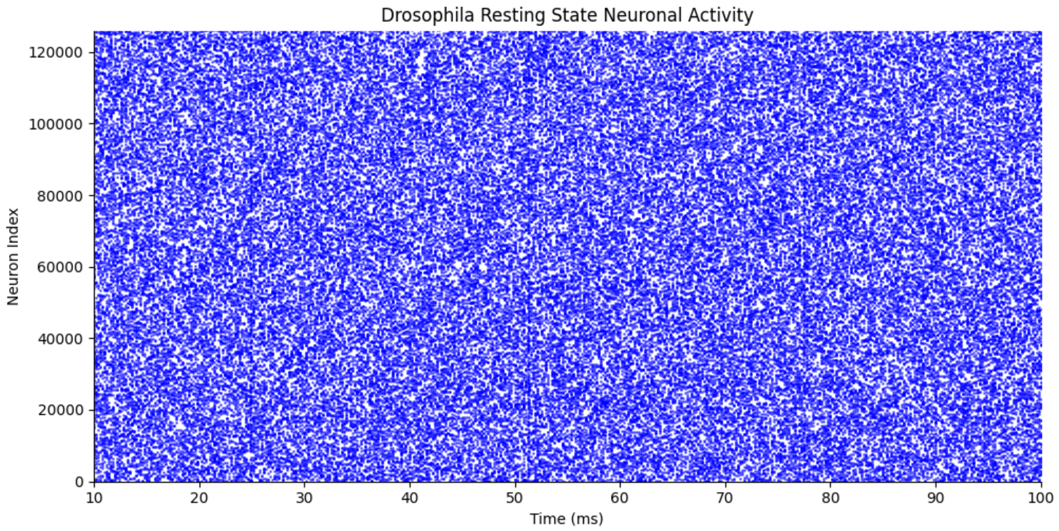

Figure S18: **Spontaneous spiking activity in an untrained neural network model during the resting state**. The raster plot shows individual neuron spike times exhibiting irregular, fluctuating firing patterns characteristic of spontaneous activity in the absence of external input or training.

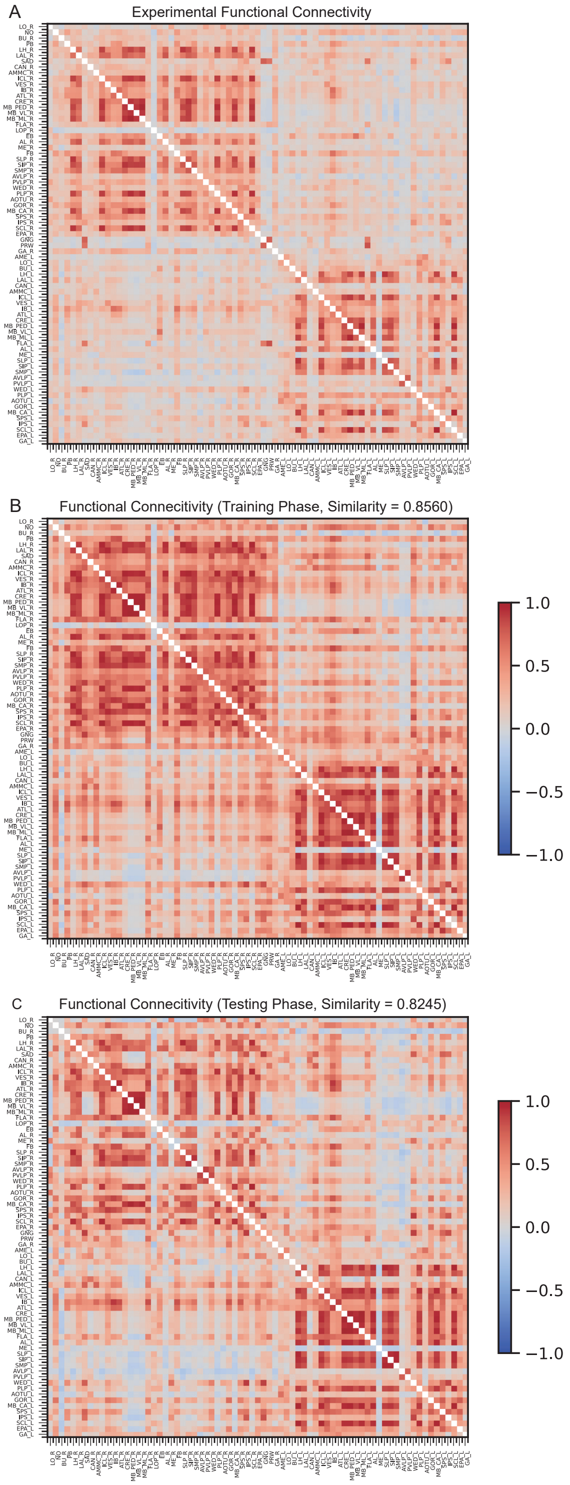

Figure S19: **Comparison of functional connectivity patterns between experimental data and model simulations.** Functional connectivity matrices showing pairwise correlations between brain regions for: (A) Experimental recordings from *Drosophila* whole-brain calcium imaging [49, 50]. (B) Training phase simulation (cosine similarity = 0.8560). (C) Testing phase simulation (cosine similarity = 0.8245). The color scale represents the correlation strength, ranging from -1 to 1.

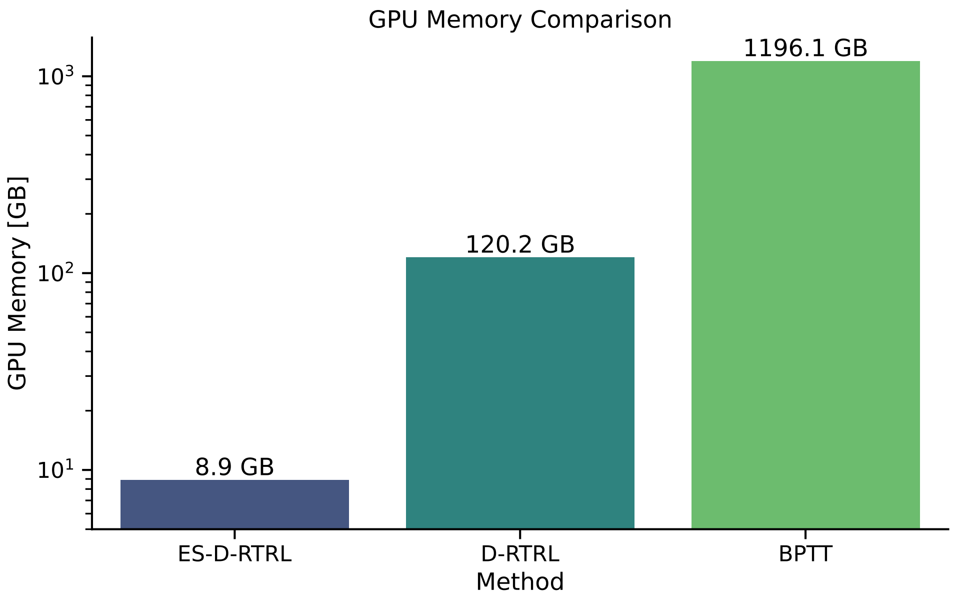

Figure S20: **Comparison of GPU Memory Usage During Whole-Brain *Drosophila* Model Fitting across Three Learning Algorithms**.

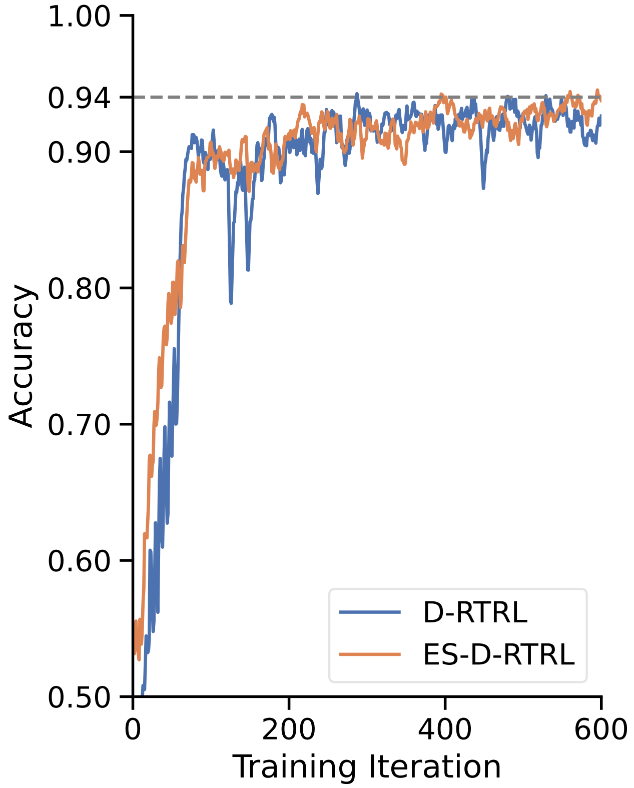

Figure S21: **Learning Performance of ES-D-RTRL and D-RTRL on the Evidence Accumulation Task over Long Time-scale Training**. In contrast to Fig. 5E, the models here were trained for significantly more epochs to examine the convergence behavior of the two algorithms on this task. As we can see, both algorithms demonstrate consistent improvement in training accuracy with increasing training epochs.

#### R Supplementary tables

Here we summarize the hyperparameters we used in all our experiments. For details, please refer to our open-source code for experimental result reproducibility (see Data availability and Code availability sections).

| Parameter | Value |
|---|---|
| Learning rule | 0.001 |
| Learning optimizer | Adam [97] |
| Batch size | 128 |
| Time step $\Delta t$ | 0.2 ms (Fig. 6)<br>1.0 ms (all other experiments) |

Table S3: Parameter settings for the learning hyperparameters.

| Parameter | Value |
|---|---|
| $\tau$ | 10.0 ms |
| $V_{\text{reset}}$ | 0. |
| $V_{\text{rest}}$ | 0. |
| $V_{\text{th}}$ | 1. |
| Hidden Size | single layer, 1000 neurons (Fig. 2)<br>single layer, 200 neurons (Fig. 4 A-D)<br>three layers, 512 neurons (Fig. 4 I-L)<br>three layers, 1024 neurons (Table 1) |
| Input Size | 1000 neurons (Fig. 2)<br>dataset dependent (Fig. 4 A-D)<br>dataset dependent (Fig. 4 I-L)<br>dataset dependent (Table 1) |

Table S4: Parameter settings for the `LIF+Delta` network in Eq. S44. The corresponding network has been used in Fig. 2, Fig. 4, and Table 1.

| Parameter | Value |
|---|---|
| $\tau$ | 10.0 ms |
| $\tau_{\mathbf{a}}$ | 100.0 ms |
| $V_{\text{reset}}$ | 0.0 |
| $V_{\text{rest}}$ | 0.0 |
| $V_{\text{th}}$ | 1.0 |
| $A$ | -0.1 |
| Hidden Size | single layer, 1000 neurons (Fig. 2)<br>single layer, 200 neurons (Fig. 4 A-D)<br>three layers, 1024 neurons (Fig. 4 I-L)<br>three layers, 1024 neurons (Table 1) |
| Input Size | 1000 neurons (Fig. 2)<br>dataset dependent (Fig. 4 A-D)<br>dataset dependent (Fig. 4 I-L)<br>dataset dependent (Table 1) |

Table S5: Parameter settings for the `ALIF+Delta` network in Eq. S45. The corresponding network has been used in Fig. 2, Fig. 4, and Table 1.

| Parameter | Value |
|---|---|
| Neuronal parameters | |
| $\tau$ | 10.0 ms |
| $V_{\text{reset}}$ | 0. |
| $V_{\text{rest}}$ | 0. |
| $V_{\text{th}}$ | 1. |
| Synaptic parameters | |
| $\tau_{\mathbf{g}}$ | 10.0 ms |
| Hidden Size | single layer, 1000 neurons (Fig. 2)<br>single layer, 200 neurons (Fig. 4 A-D) |
| Input Size | 1000 neurons (Fig. 2)<br>dataset dependent (Fig. 4 A-D) |

Table S6: Parameter settings for the `LIF+Expon` network in Eq. S46. The corresponding network has been used in Fig. 2 and Fig. 4.

| Parameter | Value |
|---|---|
| Neuronal parameters | |
| $\tau$ | 10.0 ms (Fig. 2 and Fig. 4 A-D) |
| $\tau$ | 100.0 ms (Fig. 4 E-H) |
| $\tau_{\mathbf{a}}$ | 100.0 ms (Fig. 2 and Fig. 4 A-D) |
| $\tau_{\mathbf{a}}$ | 1500.0 ms (Fig. 4 E-H) |
| $V_{\text{reset}}$ | 0. |
| $V_{\text{rest}}$ | 0. |
| $V_{\text{th}}$ | 1. |
| $A$ | -0.1 (Fig. 2 and Fig. 4 A-D) |
| $A$ | -1.0 (Fig. 4 E-H) |
| Synaptic parameters | |
| $\tau_{\mathbf{g}}$ | 10.0 ms (Fig. 2 and Fig. 4 A-D) |
| $\tau_{\mathbf{g}}$ | 100.0 ms (Fig. 4 E-H) |
| Hidden Size | single layer, 1000 neurons (Fig. 2)<br>single layer, 200 neurons (Fig. 4 A-D)<br>single layer, 200 neurons (Fig. 4 E-H) |
| Input Size | 1000 neurons (Fig. 2)<br>dataset dependent (Fig. 4 A-D)<br>100 neurons (Fig. 4 E-H) |

Table S7: Parameter settings for the `ALIF+Expon` network in Eq. S47. The corresponding network has been used in Fig. 2 and Fig. 4.

| Parameter | Value |
|---|---|
| Neuronal parameters | |
| $\tau$ | 10.0 ms |
| $V_{\text{reset}}$ | 0. |
| $V_{\text{rest}}$ | 0. |
| $V_{\text{th}}$ | 1. |
| Synaptic parameters | |
| $\tau_{\mathbf{g}}$ | 10.0 ms |
| $\tau_d$ | 500.0 ms |
| Hidden Size | single layer, 1000 neurons (Fig. 2)<br>single layer, 200 neurons (Fig. 4 A-D) |
| Input Size | 1000 neurons (Fig. 2)<br>dataset dependent (Fig. 4 A-D) |

Table S8: Parameter settings for the `LIF+STD+Expon` network in Eq. S49. The corresponding network has been used in Fig. 2 and Fig. 4.

| Parameter | Value |
|---|---|
| Neuronal parameters | |
| $\tau$ | 10.0 ms |
| $\tau_{\mathbf{a}}$ | 100.0 ms |
| $V_{\text{reset}}$ | 0. |
| $V_{\text{rest}}$ | 0. |
| $V_{\text{th}}$ | 1. |
| $A$ | -0.1 |
| Synaptic parameters | |
| $\tau_{\mathbf{g}}$ | 10.0 ms |
| $\tau_d$ | 500.0 ms |
| Hidden Size | single layer, 1000 neurons (Fig. 2)<br>single layer, 200 neurons (Fig. 4 A-D) |
| Input Size | 1000 neurons (Fig. 2)<br>dataset dependent (Fig. 4 A-D) |

Table S9: Parameter settings for the `ALIF+STD+Expon` network in Eq. S50. The corresponding network has been used in Fig. 2 and Fig. 4.

| Parameter | Value |
|---|---|
| Neuronal parameters | |
| $\tau$ | 10.0 ms |
| $V_{\text{reset}}$ | 0. |
| $V_{\text{rest}}$ | 0. |
| $V_{\text{th}}$ | 1. |
| Synaptic parameters | |
| $\tau_{\mathbf{g}}$ | 10.0 ms |
| $\tau_d$ | 100.0 ms |
| $\tau_f$ | 500.0 ms |
| Hidden Size | single layer, 1000 neurons (Fig. 2)<br>single layer, 200 neurons (Fig. 4 A-D) |
| Input Size | 1000 neurons (Fig. 2)<br>dataset dependent (Fig. 4 A-D) |

Table S10: Parameter settings for the `LIF+STP+Expon` network in Eq. S51. The corresponding network has been used in Fig. 2 and Fig. 4.

| Parameter | Value |
| --- | --- |
| Neuronal parameters | |
| $\tau$ | 10.0 ms |
| $\tau_{\mathbf{a}}$ | 100.0 ms |
| $V_{\text{reset}}$ | 0. |
| $V_{\text{rest}}$ | 0. |
| $V_{\text{th}}$ | 1. |
| $A$ | -0.1 |
| Synaptic parameters | |
| $\tau_{\mathbf{g}}$ | 10.0 ms |
| $\tau_d$ | 100.0 ms |
| $\tau_f$ | 500.0 ms |
| Hidden Size | single layer, 1000 neurons (Fig. 2)<br>single layer, 200 neurons (Fig. 4 A-D) |
| Input Size | 1000 neurons (Fig. 2)<br>dataset dependent (Fig. 4 A-D) |

Table S11: Parameter settings for the `ALIF+STP+Expon` network in Eq. S52. The corresponding network has been used in Fig. 2 and Fig. 4.

| Parameter | Value |
| --- | --- |
| Neuronal parameters | |
| $\tau_1$ | 50.0 ms |
| $\tau_2$ | 2000.0 ms |
| $\tau_{\mathtt{th}}$ | 100.0 ms |
| $\tau$ | 200.0 ms |
| $V_{\text{reset}}$ | 0. |
| $V_{\text{rest}}$ | 0. |
| $V_{\text{th}}$ | 0.9 |
| $V_{\text{th},\infty}$ | 1.0 |
| $A_1$ | 0.01 |
| $A_2$ | -1.0 |
| $R_1$ | 1.0 |
| $R_2$ | 1.0 |
| $R$ | 1.0 |
| $A_{\mathtt{th}}$ | -1.0 |
| Synaptic parameters | |
| $\tau_{\text{e}}$ | 10.0 ms |
| $\tau_{\text{i}}$ | 10.0 ms |
| $E_{\text{e}}$ | 5.0 |
| $E_{\text{i}}$ | -10.0 |
| Network parameters | |
| Input Size | 100 neurons |
| Hidden Size | single layer, 800 neurons |
| EI ratio | 4:1 |
| Recurrent connection probability | 0.1 |

Table S12: Parameter settings for the E/I network in Eq. S55. The corresponding network has been used in Fig. 5.

[10] Guillaume Bellec, Darjan Salaj, Anand Subramoney, Robert Legenstein, and Wolfgang Maass. Long short-term memory and learning-to-learn in networks of spiking neurons. Advances in neural information processing systems, 31, 2018.

[28] Anand Subramoney, Khaleelulla Khan Nazeer, Mark Schöne, Christian Mayr, and David Kappel. Efficient recurrent architectures through activity sparsity and sparse back-propagation through time. In The Eleventh International Conference on Learning Representations, 2023.

[29] Kyunghyun Cho, Bart van Merrienboer, Çaglar Gülçehre, Dzmitry Bahdanau, Fethi Bougares, Holger Schwenk, and Yoshua Bengio. Learning phrase representations using rnn encoder–decoder for statistical machine translation. In Conference on Empirical Methods in Natural Language Processing, 2014.

[35] Asier Mujika, Florian Meier, and Angelika Steger. Approximating real-time recurrent learning with random kronecker factors. Advances in Neural Information Processing Systems, 31, 2018.

[36] Arnon Amir, Brian Taba, David Berg, Timothy Melano, Jeffrey McKinstry, Carmelo Di Nolfo, Tapan Nayak, Alexander Andreopoulos, Guillaume Garreau, Marcela Mendoza, et al. A low power, fully event-based gesture recognition system. In Proceedings of the IEEE Conference on Computer Vision and Pattern Recognition, pages 7243–7252, 2017.

[37] Paul J. Werbos. Generalization of backpropagation with application to a recurrent gas market model. Neural Networks, 1:339–356, 1988.

[38] Michael C. Mozer. A focused backpropagation algorithm for temporal pattern recognition. Complex Syst., 3, 1989.

[39] Ronald J. Williams and Jing Peng. An efficient gradient-based algorithm for on-line training of recurrent network trajectories. Neural Computation, 2:490–501, 1990.

[40] Anthony J Robinson and Frank Fallside. The utility driven dynamic error propagation network, volume 1. University of Cambridge Department of Engineering Cambridge, 1987.

[41] Ronald J Williams and David Zipser. Experimental analysis of the real-time recurrent learning algorithm. Connection science, 1(1):87–111, 1989.

[42] Ronald J Williams and David Zipser. A learning algorithm for continually running fully recurrent neural networks. Neural computation, 1(2):270–280, 1989.

[43] Thomas Bohnstingl, Stanisław Woźniak, Angeliki Pantazi, and Evangelos Eleftheriou. Online spatio-temporal learning in deep neural networks. IEEE Transactions on Neural Networks and Learning Systems, 2022.

[44] Benjamin Cramer, Yannik Stradmann, Johannes Schemmel, and Friedemann Zenke. The heidelberg spiking data sets for the systematic evaluation of spiking neural networks. IEEE Transactions on Neural Networks and Learning Systems, 2020.

[59] James Bradbury, Roy Frostig, Peter Hawkins, Matthew James Johnson, Chris Leary, Dougal Maclaurin, George Necula, Adam Paszke, Jake VanderPlas, Skye Wanderman-Milne, and Qiao Zhang. JAX: composable transformations of Python+NumPy programs, 2018.

[69] Răzvan V Florian. The chronotron: A neuron that learns to fire temporally precise spike patterns. 2012.

[70] Robert Gütig and Haim Sompolinsky. The tempotron: a neuron that learns spike timing–based decisions. Nature neuroscience, 9(3):420–428, 2006.

[71] Corentin Tallec and Yann Ollivier. Unbiased online recurrent optimization. arXiv preprint arXiv:1702.05043, 2017.

[72] Frederik Benzing, Marcelo Matheus Gauy, Asier Mujika, Anders Martinsson, and Angelika Steger. Optimal kronecker-sum approximation of real time recurrent learning. In International Conference on Machine Learning, pages 604–613. PMLR, 2019.

[73] Jacob Menick, Erich Elsen, Utku Evci, Simon Osindero, Karen Simonyan, and Alex Graves. Practical real time recurrent learning with a sparse approximation. In International conference on learning representations, 2021.

[74] Mingqing Xiao, Qingyan Meng, Zongpeng Zhang, Di He, and Zhouchen Lin. Online training through time for spiking neural networks. Advances in Neural Information Processing Systems, 35:20717–20730, 2022.

[75] Thomas M Summe, Clemens JS Schaefer, and Siddharth Joshi. Estimating post-synaptic effects for online training of feed-forward snns. In 2024 International Conference on Neuromorphic Systems (ICONS), pages 264–271. IEEE, 2024.

[76] Haiyan Jiang, Giulia De Masi, Huan Xiong, and Bin Gu. Ndot: neuronal dynamics-based online training for spiking neural networks. In Forty-first International Conference on Machine Learning, 2024.

[77] Marco Paul E. Apolinario and Kaushik Roy. S-TLLR: STDP-inspired temporal local learning rule for spiking neural networks. Transactions on Machine Learning Research, 2025.

[78] Nicholas T Carnevale and Michael L Hines. The NEURON book. Cambridge University Press, 2006.

[79] Dan FM Goodman and Romain Brette. The brian simulator. Frontiers in neuroscience, 3:643, 2009.

[80] Trevor Bekolay, James Bergstra, Eric Hunsberger, Travis DeWolf, Terrence C Stewart, Daniel Rasmussen, Xuan Choo, Aaron Russell Voelker, and Chris Eliasmith. Nengo: a python tool for building large-scale functional brain models. Frontiers in neuroinformatics, 7:48, 2014.

[81] Chris Eliasmith and Charles H Anderson. Neural engineering: Computation, representation, and dynamics in neurobiological systems. MIT press, 2003.

[82] Chris Eliasmith, Terrence C Stewart, Xuan Choo, Trevor Bekolay, Travis DeWolf, Yichuan Tang, and Daniel Rasmussen. A large-scale model of the functioning brain. science, 338(6111):1202–1205, 2012.

[83] Jason K Eshraghian, Max Ward, Emre Neftci, Xinxin Wang, Gregor Lenz, Girish Dwivedi, Mohammed Bennamoun, Doo Seok Jeong, and Wei D Lu. Training spiking neural networks using lessons from deep learning. Proceedings of the IEEE, 111(9):1016–1054, 2023.

[84] Christian Pehle and Jens Egholm Pedersen. Norse - A deep learning library for spiking neural networks, January 2021. Documentation: https://norse.ai/docs/.

[85] Sadique Sheik, Gregor Lenz, Felix Bauer, and Nogay Kuepelioglu. SINABS: A simple Pytorch based SNN library specialised for Speck.

[91] Kade M Heckel and Thomas Nowotny. Spyx: A library for just-in-time compiled optimization of spiking neural networks. arXiv preprint arXiv:2402.18994, 2024.

[97] Diederik P Kingma and Jimmy Ba. Adam: A method for stochastic optimization. arXiv preprint arXiv:1412.6980, 2014.
